# LCK-targeting molecular glues overcome resistance to inhibitor-based therapy in T-cell acute lymphoblastic leukemia

**DOI:** 10.1101/2025.07.03.663042

**Authors:** Satoshi Yoshimura, Marisa Actis, Justin T. Seffernick, Jamie A. Jarusiewicz, Anup Aggarwal, Angelina Li, Yong Li, DongGeun Lee, Lei Yang, Anand Mayasundari, Zoran Rankovic, Marcus Fischer, Gisele Nishiguchi, Jun J. Yang

## Abstract

Drug resistance is a major challenge in cancer therapy, especially in the context of kinase inhibitors. While targeted protein degradation (TPD) was a distinct mode of action compared to inhibition-based therapeutic targeting, the potential value of TPD in drug-resistant cancer remains unclear. Here, we report the discovery of cereblon-recruiting molecular glue degraders (MGDs) targeting LCK, an oncogenic kinase in T-cell acute lymphoblastic leukemia (T-ALL). By high-throughput screening and medicinal chemistry optimization, we developed a series of MGDs that induced CRBN-dependent degradation of LCK as well as potent cytotoxicity in T-ALL *in vitro*. Structure-activity relationship analysis and ternary complex modeling revealed a non-canonical degron at the LCK-CRBN interface involving the G-loop, whose mutation disrupts this interaction. Unlike inhibitors and inhibitor-based PROTACs, these MGDs engage LCK in regions distal to the ATP binding site and thus their activities in T-ALL are not affected by gate-keeper LCK mutations that drive resistance to inhibitor-based therapeutics. Taken together, our data underscore the potential of LCK-targeting MGDs as a strategy to overcome kinase inhibitor resistance in T-ALL, highlighting a potentially generalizable strategy in cancer therapy.

## Introduction

Targeted protein degradation (TPD) has garnered growing popularity in drug discovery thanks to several key advantages over protein inhibition, including a catalytic mechanism of action, greater efficacy and the ability to target non-functional binding sites^1,2^. The two most explored chemical approaches in TPD are PROteolysis TArgeting Chimeras (PROTACs) and Molecular Glue Degraders (MGDs)^3^. Both modalities harness the endogenous cellular ubiquitin proteasome system by bringing a protein of interest (POI) into proximity to an E3 ligase to induce ubiquitination and subsequent degradation of the former by the proteasome^3^. PROTACs can be rationally designed as bi-functional molecules, i.e., chemically linking a small molecule binder of a POI (e.g., a known inhibitor of a therapeutic target) to an E3 ligase ligand^1^. While the irreversible target degradation induced by PROTACs may be translated into greater efficacy than inhibitors, drug resistance is a significant potential challenge. For example, any mutation in the POI that compromises the binding of the inhibitor would, in theory, result in resistance to PROTACs designed around this inhibitor. For instance, in the case of leukemia, gate-keeper mutations in ABL (e.g. T315I) that cause resistance to dasatinib would greatly reduce the activity of all dasatinib-based PROTACs^4–7^.

By contrast, MGDs induce conformational changes at the surface of the E3 ligase that are complemented by a POI to form the required ternary complex; this proximity leads to target protein ubiquitination and eventually proteasomal degradation^8^. The key feature of a MGD is that it usually engages a POI through surface interaction, which can be distant from the binding pocket deep inside the protein that is required for inhibitor-target interaction. Because of this, we reason that MGDs may be less impacted by mutations in the inhibitor binding pocket and should remain active in cells resistant to inhibitor or inhibitor-based PROTACs. However, there is a paucity of examples clearly demonstrating the potential of MGDs to overcome PROTAC resistance, partly because very few therapeutic targets have been evaluated with both types of protein degradation modalities^9^.

LCK, lymphocyte-specific tyrosine kinase, is constitutively activated in ∼40% of T-cell acute lymphoblastic leukemia (T-ALL)^10–13^. Therapeutic targeting of LCK with small molecule inhibitors such as dasatinib can be highly effective in inducing leukemia cell death both in the lab and in patients^10,12,14,15^. Dasatinib-based PROTACs induce rapid degradation of LCK in T-ALL, with cytotoxicity far exceeding dasatinib itself as well as greatly superior pharmacodynamic effects *in vivo*^16,17^. However, an LCK mutation has been identified (T316M or T316I) that abolishes dasatinib binding^14,18^, with expected resistance to a dasatinib-based PROTAC. We hypothesized that we could overcome resistance associated with kinase domain mutations by developing LCK-targeted MGDs. In fact, LCK contains the common β-hairpin G-loop motif, known as degron, present in the most frequently reported neosubstrates (GSPT1, Casein kinase 1α (CK1α), and IKZF1) of cereblon (CRBN) ligase, providing a rationale for this approach.

Herein, we report the discovery and optimization of a series of LCK-targeting MGDs that shows superior cytotoxicity in dasatinib-resistant T-ALL cells compared to dasatinib and dasatinib-based PROTAC^16^. Importantly, our data points to the potential of switching degradation modalities from bivalent to monovalent as a means of overcoming PROTAC resistance.

## RESULTS

### High-throughput screening of molecular glue library identifies LCK degraders

To identify MGDs that target LCK, we designed a screening strategy where the St. Jude proprietary molecular glue library was assayed in a HiBiT-tagged system and evaluated for degradation of LCK (**Fig. 1A, B**)^19^.

**Fig. 1:**
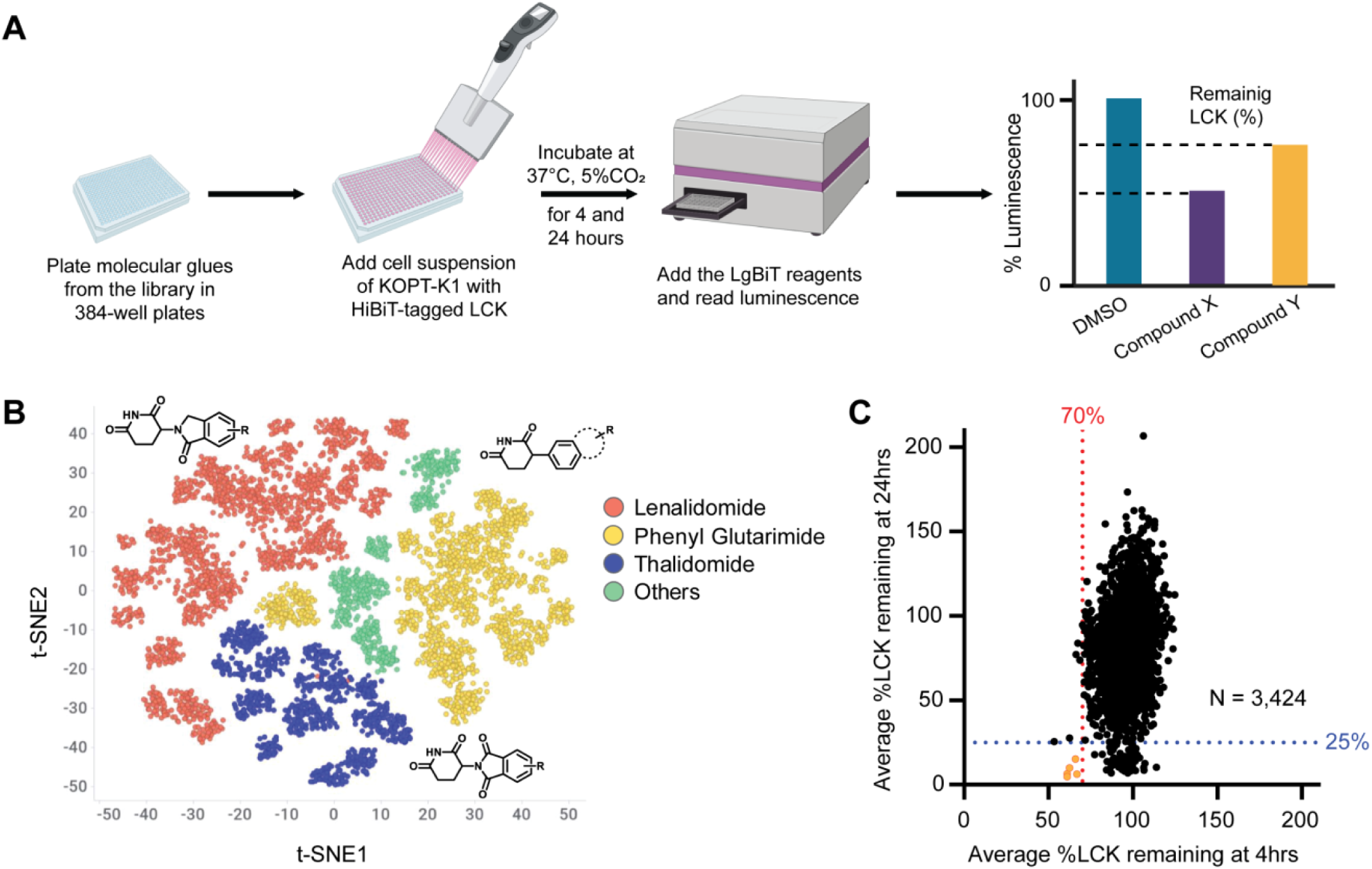
High-throughput screening to identify LCK-targeting MGDs **A**, Schematic workflow of high-throughput screening of LCK-targeting MGDs. **B**, Visualization of the chemical scaffold diversity of the molecular glue library. In this t-SNE plot, each dot represents a compound and the proximity between points indicates their chemical structure similarities. **C**, LCK degradation induced by each MGD at early (4 hrs) and late (24 hrs) time points. Each dot represents the mean percentage of LCK remaining, calculated from duplicate measurements for each compound. Five compounds (orange) met the threshold of ≥30% and ≥75% LCK-HiBiT reduction at 4 hrs and 24 hrs, respectively.

Using CRISPR editing, we first engineered a T-ALL cell line model (KOPT-K1) in which the constitutively-activated LCK is fused with a HiBiT tag^17^, enabling quantification of LCK degradation using luminescence. Then the St. Jude proprietary molecular glue library^20^, consisting of 3,424 compounds designed for potential cereblon engagement and comprising of chemically diverse chemotypes, was screened in the T-ALL cell line using the LCK HiBiT assay at a single concentration (40 µM) (**Fig. 1A**). For each compound, the LCK-HiBiT protein level in treated cells was measured and normalized to that of non-treated cells. Because LCK is essential for T-ALL survival, degraders are expected to induce apoptosis^16^. To account for the effect of cell death on LCK-HiBiT protein level, we performed the assay after both 4 and 24 hrs of drug exposure. Of the 3,424 compounds screened, there was a greater degree of reduction in LCK protein at 24 hrs than at 4 hrs (*P* < 0.0001), and more compounds degraded LCK to the same degree at 24 hrs than at 4 hrs. Five hits (**SJ1473**, **1481**, **1581**, **1652**, and **2917**) induced ≥30% and ≥75% of LCK-HiBiT degradation at 4 and 24 hrs, respectively, and were selected for further characterization (**Fig. 1C**).

Next, these five compounds were tested in a dose-response format using the LCK-HiBiT assay at serial concentrations ranging from 2 nM–40 µM of drug exposure. Compound **1** (SJ1581) exhibited the greatest potency (**Fig. 2A**,**B**), while the other 4 induced only modest degradation. We also directly measured each compound’s ability to promote the formation of the ternary complex with CRBN and LCK, using an *in vitro* AlphaScreen assay^16^, as the induced proximity of the two is required for ubiquitination and subsequent proteasomal degradation. The AlphaScreen signal was measured for each molecular glue at varying concentrations. As a positive control, a previously described dasatinib-based LCK PROTAC (**SJ11646**) was included^16^, giving the characteristic bell-shaped curve (**Fig. 2C**). Of the five compounds, **1** was most effective at forming a ternary complex with a half-maximum concentration (EC_50_) of 123.1 nM (**Table 1**), in line with its potency in degrading LCK quantified by the HiBiT assay.

**Fig. 2:**
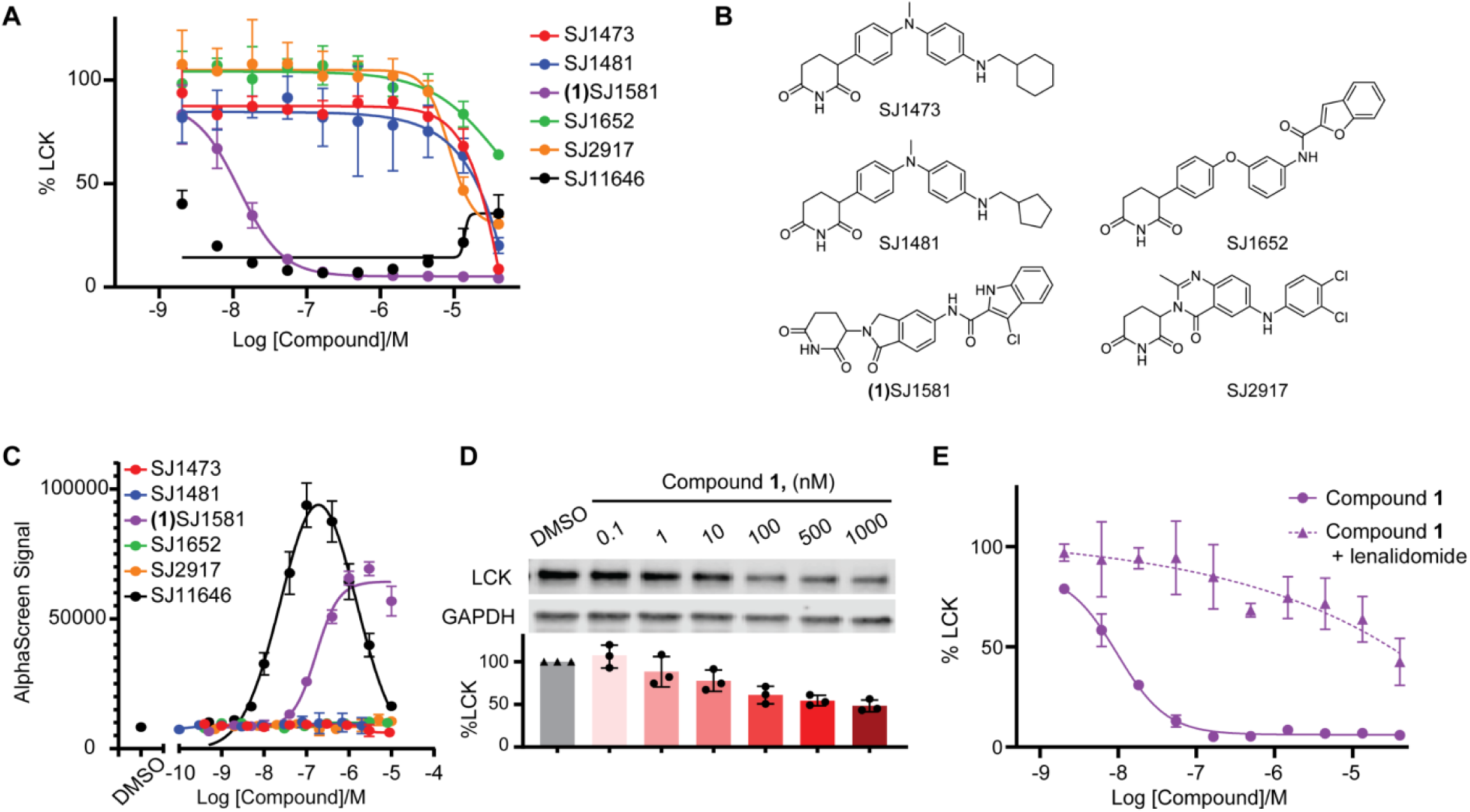
Compound 1 is validated as a hit MGD identified through compound screening. **A**, Dose-dependent LCK degradation by the five compounds identified from the screening, measured using HiBiT assay after 24 hrs of treatment. Compound **1** exhibits the greatest potency among them. Data represents mean ± SD from three biological replicates. **B**, Structures of the five compounds identified from the screening. **C**, MGD-induced formation of a ternary complex between GST-LCK and His-CRBN-DDB1 proteins measured using an *in vitro* AlphaScreen assay. Data is shown as mean ± SD from three technical replicates. **D**, Representative immunoblots showing LCK degradation in KOPT-K1_LCK_HiBiT cells treated with serial concentrations of compound **1** for 24 hrs. Quantified LCK degradation normalized to the DMSO control from three biological replicates are also shown. **E**, Dose-dependent LCK degradation by compound **1** after 24 hr-treatment demonstrated by HiBiT assay, which was attenuated by co-treatment with lenalidomide 10 µM. Data represents mean ± SD from three biological replicates.

**Table 1.** Properties of LCK MGDs 1–8.

| ID | X | $R_1$ | $R_2$ | $R_3$ | Ternary $EC_{50}$ (nM)<sup>a</sup> | LCK $DC_{50}$ 4 h (nM)<sup>b</sup> | LCK $DC_{50}$ 24 h (nM)<sup>b</sup> |
| --- | --- | --- | --- | --- | --- | --- | --- |
| **1** | CO | Cl | H | H | 123.1 (103.2 – 153.7) | >40000 | $8.7 \pm 2.2$ |
| **2** | CO | Me | H | H | 109.9 (97.8 – 123.3) | >40000 | $21.3 \pm 19.2$ |
| **3** | CO | H | H | H | 234.7 (163.5 – 347.9) | >40000 | $268.4 \pm 112.9$ |
| **4** | CO | Cl | Me | H | >1000 | >40000 | $26749.2 \pm 22951.1$ |
| **5** | CO | Me | H | Me | >1000 | >40000 | >40000 |
| **6** | CH<sub>2</sub> | Me | H | H | >1000 | >40000 | >40000 |
| **7** | CO | Et | H | H | 97.4 (85.4 – 110.9) | $20.4 \pm 6.8$ | $2.1 \pm 0.1$ |
| **8** | CO | Pr | H | H | 194.5 (182.7 – 207.2) | >40000 | $10.9 \pm 3.7$ |

<sup>a</sup>AlphaScreen ternary complex formation half-maximum concentration reported as the mean with 95% confidence interval (CI) for three technical replicates. <sup>b</sup>Degradation measured at the indicated time points using HiBiT-LCK, reported as the mean ± standard deviation for three biological replicates.

LCK degradation by compound **1** was also confirmed using immunoblotting (**Fig. 2D**, **Fig. S1**). Pre-treatment with excess lenalidomide abolished CRBN-LCK ternary complex formation (**Fig. S2**) and reversed the compound **1**-induced LCK degradation in T-ALL cells (DC_50_ > 40 µM), confirming these effects as CRBN-dependent (**Fig. 2E**).

These results qualified high-throughput screening hit compound **1** as a potent and CRBN-dependent LCK degrader for further optimization.

### Medicinal chemistry optimization to improve potency and solubility of LCK MGDs

To understand the structure-activity relationship (SAR) for LCK activity, we first generated a series of analogs of compound **1** probing key structural features, i.e., the halide at the 3-position of the indole (R_1_), the NH’s of the indole and amide (R_2_ and R_3_, respectively), and the carbonyl (X) linking the CRBN binding moiety to the indole (**Table 1**). Using the AlphaScreen assay, we observed a loss in the ability to form a ternary complex when removing the halide (compound **3,** EC_50_ = 234.7 nM vs. compound **1**, EC_50_ = 123.1 nM), which also led to reduced degradation potency as determined using the LCK-HiBiT assay (DC_50_ = 268.4 nM at 24 hrs treatment). Replacing the chloro at R_1_ with a methyl substitution (**2**) slightly improved its ability to form the ternary complex (EC_50_ = 109.9 nM), while incorporating an ethyl substitution provided further improvement (**7**, EC_50_ = 97.4 nM) and greatly increased LCK-HiBiT degradation activity, providing the first analog with activity at 4 hrs of drug treatment (DC_50_ = 20.4 nM). These improvements diminished when incorporating a larger propyl group at R_1_ (**8**), suggesting there is a limit to the size tolerated at that position.

We next explored the impact of the indole N-H and amide functional group. As shown in **Table 1**, methylating R_2_ or R_3_ gave compounds (**4** and **5**, respectively) with loss of activity across assays, highlighting the importance of hydrogen bond donors in the formation of the ternary complex. Removing the carbonyl of the amide functional group and losing this hydrogen bond acceptor resulted in inactive compound **6**. The SAR pointed to an essential network of H-bond donors and acceptors for ternary complex formation.

Based on these results, we selected compound **7** for further structural modifications with the goal of systematically exploring the impact of indole substitutions on ternary complex formation and LCK degradation activity, using DC_50_ values at 4hrs as the endpoint to prioritize the most potent degraders (**Table 2**).

**Table 2.** Properties of LCK MGDs 9–15.

| ID | R<sub>1</sub> | R<sub>2</sub> | Ternary EC<sub>50</sub> (nM)<sup>a</sup> | LCK DC<sub>50</sub> 4 h (nM)<sup>b</sup> | Sol. (μM)<sup>c</sup> |
| --- | --- | --- | --- | --- | --- |
| **9** | 5-Br | H | 84.7 (75.8 – 94.7) | 22.1 ± 2.5 | 0.77 ± 0.05 |
| **10** | 6-Br | H | 128.6 (120.0 – 137.9) | 2614.1 ± 855.9 | 0.56 ± 0.13 |
| **11** | 7-Br | H | 62.1 (58.0 – 66.4) | 68.6 ± 35.7 | 1.06 ± 0.03 |
| **12** | 5-OMe | H | 61.0 (55.8 – 66.7) | 33.2 ± 14.2 | 6.77 ± 0.24 |
| **13** | 6-OMe | H | 52.9 (36.6 – 75.8) | 23.1 ± 6.3 | 5.38 ± 1.79 |
| **14** | 7-OMe | H | 101.1 (96.8 – 105.6) | 15.9 ± 3.4 | 3.21 ± 0.98 |
| **15** | 7-Br | Me | >1000 | >40000 | 0.44 ± 0.03 |

<sup>a</sup>AlphaScreen ternary complex formation half-maximum concentration reported as the mean with 95% confidence interval (CI) for three technical replicates. <sup>b</sup>Degradation measured using HiBiT-LCK reported as the mean ± standard deviation for three biological replicates. <sup>c</sup>Kinetic aqueous solubility measured at pH 7.4 and reported as the mean ± standard deviation for three replicates.

Firstly, focusing on bromo substitutions, we found that the 5-Br (**9**) and 7-Br (**11**) analogues were favored for inducing proximity in the AlphaScreen assay (EC_50_ = 84.7 and 62.1 nM, respectively) over the 6-Br substitution (**10,** EC_50_ = 128.6 nM). In agreement with the biochemical data, compounds **9** and **11** were highly effective in LCK degradation (DC_50_ = 22.1 and 68.6 nM, respectively). Incorporating a methoxy group at these positions provided compounds **12**, **13** and **14** with comparable potencies in LCK:CRBN ternary complex assay and LCK degradation (DC_50_ = 33.2, 23.1 and 15.9 nM, respectively). This representative set of compounds suggested that indole substitutions were tolerated, and they could be exploited to improve on ADME properties later in the optimization process. To mechanistically validate the activity of these molecules, compound **15** was prepared, featuring a methyl substitution on the glutarimide nitrogen, which blocks key interactions in the thalidomide binding pocket of CRBN. Compound **15** completely lost its ability to form a ternary complex, which correlated with loss of activity for LCK degradation (**Table 2**). These results support a CRBN-dependent degradation mechanism.

We next focused on physicochemical physical properties and profiled compounds in **Table 2** against our aqueous solubility assay. All compounds displayed poor to modest aqueous solubility (0.44 -6.77 µM) and to overcome this liability, we explored adding polar substitutions at the 7-position of the indole ring. The addition of an amide group in compounds **16** and **18** yielded glues with strong LCK degradation (**Table 3**), while only compound **16** gave improved solubility (22.6 µM) compared to **17** (3.7 µM). Incorporating a primary amine (**17**) or alcohol (**19**) significantly improved solubility (67.9 and 27.0 µM, respectively) while retaining LCK degradation activity (DC_50_ = 103.4 and 32.4 nM, respectively). Encouraged by the profiles of compounds **17** and **19**, we pursued further characterization of these chemical tools, probing their binding modes via molecular modeling, neosubstrate selectivity and activity in dasatinib-resistant cell lines.

**Table 3.** Properties of LCK MGDs 16–19.

| ID | R<sub>1</sub> | Ternary<br>EC<sub>50</sub><br>(nM)<sup>a</sup> | LCK<br>DC<sub>50</sub> 4 h<br>(nM)<sup>b</sup> | KOPT-K1<br>LC<sub>50</sub><br>(nM)<sup>c</sup> | KOPT-K1<br>LCK T316I<br>LC<sub>50</sub> (nM)<sup>c</sup> | Sol. ($\mu\text{M}$)<sup>d</sup> |
| --- | --- | --- | --- | --- | --- | --- |
| **16** | NHCOMe | 67.3 (63.9 – 71.0) | 43.2 ± 16.0 | 32.2 ± 3.9 | 85.6 ± 16.1 | 22.6 ± 4.1 |
| **17** | CH<sub>2</sub>NH<sub>2</sub> | 77.1 (70.2<br>– 84.5) | 103.4 ±<br>20.6 | 32.9 ±<br>5.2 | 63.0 ± 3.3 | 67.9 ± 5.0 |
| **18** | CH<sub>2</sub>NHCOMe | 52.9 (48.7<br>– 57.4) | 65.0 ±<br>21.6 | 17.9 ±<br>0.1 | 56.9 ± 0.5 | 3.7 ± 0.1 |
| **19** | CH<sub>2</sub>OH | 65.1 (61.2<br>– 69.2) | 32.4 ±<br>14.5 | 11.5 ±<br>0.6 | 27.9 ± 2.4 | 27.0 ± 0.3 |

<sup>a</sup>AlphaScreen ternary complex formation half-maximum concentration reported as the mean with 95% confidence interval (CI) for three technical replicates. <sup>b</sup>Degradation measured using HiBiT-LCK reported as the mean ± standard deviation for three biological replicates. <sup>c</sup>Cytotoxicity measured using CTG assay reported as the mean ± standard deviation for three biological replicates. <sup>d</sup>Kinetic aqueous solubility measured at pH 7.4 and reported as the mean ± standard deviation for three replicates.

### Modeling of CRBN-MGD-LCK interaction rationalizes structure-activity relationship

Generally, IMiD-based MGDs are thought to degrade target protein through a glycine containing β-hairpin loop recognition motif, known as the “degron”. However, this view was recently challenged in work by Petzold *et al*.^21^ which described non-β-hairpin glycine containing degrons recognized by CRBN to form the required ternary complex. In particular, the authors reported an MGD-mediated interaction of CRBN with inactive LCK through its helical G-loop motif, specifically at G415.

To validate the importance of the G415-containing degron, we generated a mutant LCK protein (G415N), with which we screened a panel of LCK MGDs for their ability to form a ternary complex with CRBN. None of the LCK MGDs tested (**16 – 19**) induced the formation of the ternary complex between CRBN and mutant LCK (**Fig. 3A**). By contrast, the mutant LCK can still interact with CRBN in the presence of LCK PROTAC **SJ11646** because this interaction does not involve the G415 degron. This was further supported by immunoblotting, which showed that LCK degradation by MGDs was impaired in KOPT-K1 with LCK G415N (**Fig. 3B**).

**Fig. 3:**
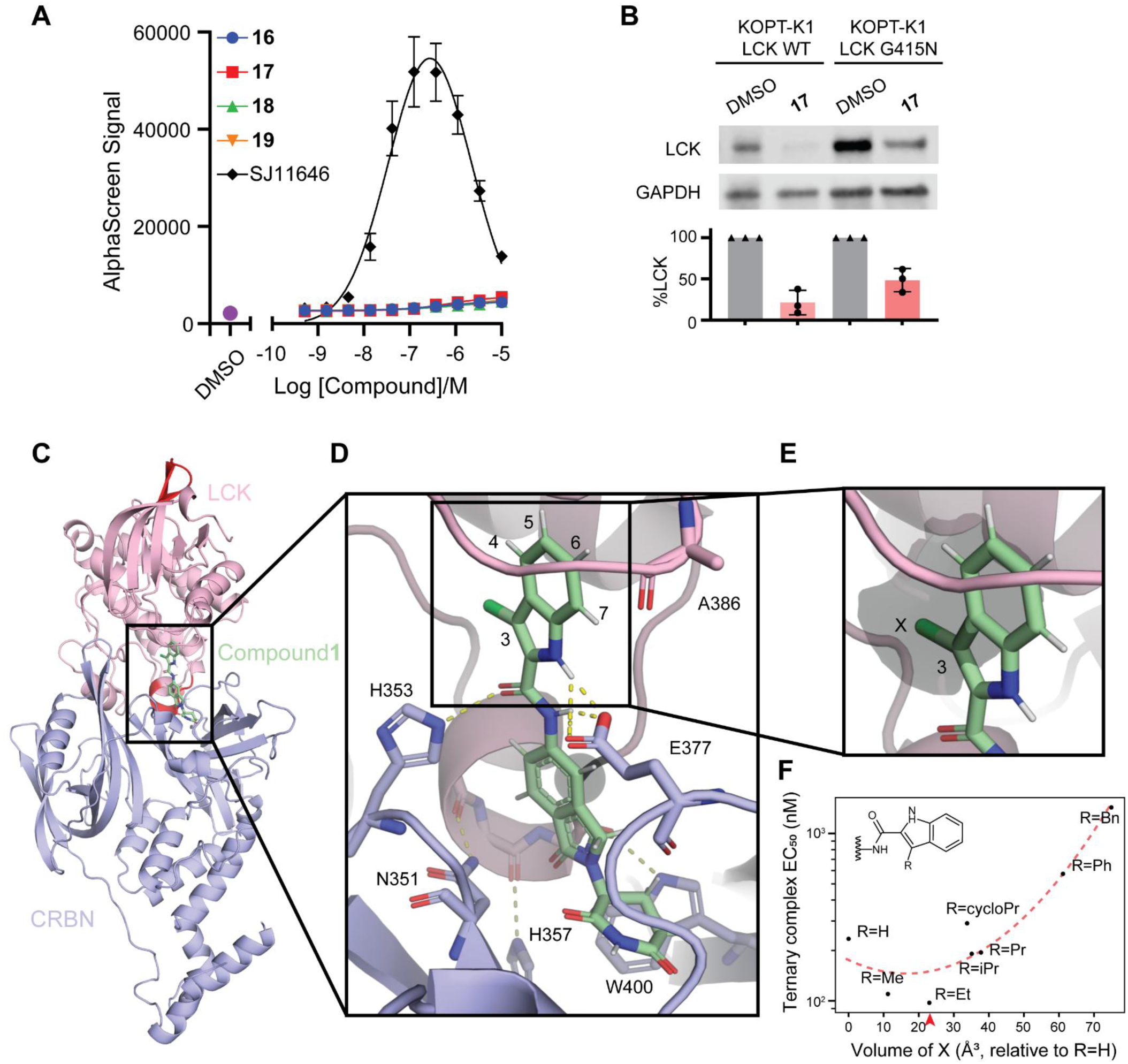
Modeling of CRBN-MGD-LCK interaction rationalizes structure-activity relationship. **A**, *In vitro* AlphaScreen assay demonstrating ternary complex formation between His-CRBN-DDB1 and GST-LCK (G415N) in the presence of MGD. **B**, Representative immunoblots showing attenuated LCK degradation KOPT-K1 with LCK G415N mutant treated with compound **17** at 1000 nM for 24hrs, compared to cells expressing wildtype LCK. Quantified LCK degradation normalized to the DMSO control from three biological replicates are also shown. **C-E**, Model of the ternary complex, CRBN-**1**-LCK, with the helical G-loop motif G415 of LCK: **C**, Wide view of the modeled ternary complex. β-hairpin loop (top) and G-loop (middle) are highlighted in red. **D**, Close-up view of the interactions between CRBN, compound **1**, and LCK with G415 shown in black with C-alpha as sphere. **E**, Binding pocket of LCK corresponding to the 3 position in the indole ring (X = Cl). The pocket shown (black) is a representative size from the volume calculations. **F**, Ternary complex EC_50_ values for eight different aliphatic R groups (R = H, Me, Et, Pr, iPr, cycloPr, Ph, and Bn). The relative volume of the ethyl (Et) group most closely matched the volume calculated from the structure and was associated with the most robust ternary complex formation. The red arrowhead indicates the average modeled pocket volume (23.2Å³). The red dashed line represents a quadratic trendline.

To provide structural insight into the observed activities of different LCK MGDs in the absence of a co-crystal structure, we developed models of the ternary complex for various compounds in **Tables 1–3**. **Fig. 3C** shows a model of CRBN-**1**-LCK. We prioritized models where the three backbone carbonyls (G-3, G-2, and G-1) on the helical G-loop motif on LCK form hydrogen bonds (H-bonds) to N351, H357, and W400 in CRBN, respectively (**Fig. 3D**), to understand the SARs in **Tables 1–3**. First, our models contained H-bonds between the two N-Hs (R_2_ and R_3_ in **Table 1**) on the MGD to E377 on CRBN, matching the SAR of N-H over N-methyl preferred at that position (**Table 1**). Similarly, we observed an H-bond between the carbonyl (X in **Table 1**) and H353 in CRBN, that has been shown to be involved in other quaternary complexes^20^. In our models, substitutions at the 3-position of the indole point in the direction of LCK and engage a small, hydrophobic pocket. Based on an ensemble of 250 models of CRBN-**1**-LCK, the volume of the pocket engaged by substitutions at this position was on average 23.2 Å^3^ (**Fig. 3E**). The pocket size agrees well with the SAR observed in **Table 1** where the ethyl group in compound **7** (with an approximate volume of 23 Å^3^) outperforms all other analogous compounds (larger and smaller), as shown in **Fig. 3F** for eight aliphatic groups (**Table S3**).

While our model explains this SAR, to accommodate binding at the 4-7 positions of the MGD indole ring, the flexible LCK activation loop, known to play a role in inactive LCK, requires a conformational change (**Table 2**). This suggests that substitutions at the 7-position are directed towards a more solvent exposed region (also towards CRBN) may be preferable as these may not need to pay the energetic cost to reorganize the activation loop. Our model suggests two possible interactions for the substitutions at the 7-position in **Table 3**: the backbone carbonyl in A386 in the LCK activation loop within ∼3 Å, and E377 in CRBN at ∼4.5 Å; both distances are consistent with a ∼2 bond-length group, such as those in **17** and **19**.

### LCK MGDs overcome leukemia resistance to LCK inhibitor and inhibitor-based PROTAC

LCK is a selective therapeutic target in T-ALL and inhibitors such as dasatinib are highly effective^12,15^. However, leukemia can develop drug resistance by acquiring mutations in the LCK gene, especially at the gatekeeper position T316^14,18^. The mutant LCK protein is expected to lose its ability to bind dasatinib and this should also extend to dasatinib-based PROTACs. To test this, we established isogenic clones of KOPT-K1 cells with either mutant or wildtype LCK. As shown in **Fig. 4A**, T316I mutation led to a 4.8-and10.0-fold increase in AUC of dasatinib and SJ11646, respectively, compared to cells with wildtype LCK. By contrast, compounds **16-19** retained strong cytotoxicity even in KOPT-K1 cells harboring LCK mutation (LC_50_ 27.9–85.96 nM; **Fig. 4B**). Consistent with this, LCK degradation by the PROTAC was greatly compromised in the mutant cell line relative to parental control, whereas MGDs induced a comparable level of LCK degradation in KOPT-K1 cells regardless of LCK genotype (**Fig. 4C**). We reason that this can be attributed to the fact that MGDs engage LCK in a region distal to the ATP binding pocket used by dasatinib and dasatinib-derived PROTACs. (**Fig. 4D**). Therefore, LCK MGDs can overcome small molecule inhibitor resistance, which is a liability to treatment with an inhibitor-based PROTAC.

**Fig. 4:**
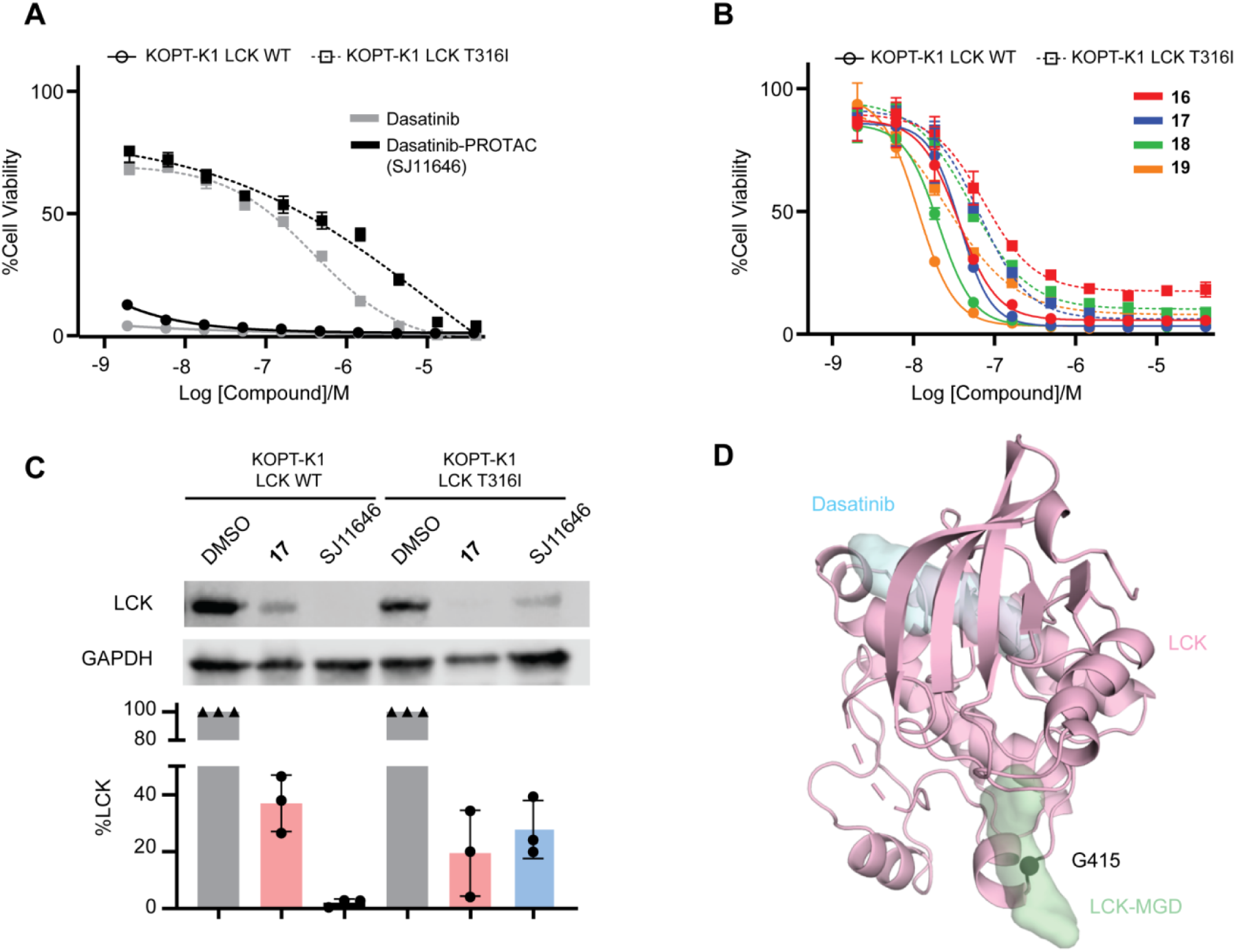
LCK MGDs overcome leukemia resistance to LCK inhibitors and PROTAC. **A**,**B**, Dose-dependent cytotoxicity in parental and dasatinib-resistant KOPT-K1 harboring a heterozygous LCK gate-keeper mutation T316I by dasatinib or a dasatinib-PROTAC SJ11646 (**A**), or LCK-MGDs (**B**), measured by CTG assay. The cytotoxicity was retained by LCK-MGDs, but greatly shifted by dasatinib or the dasatinib-PROTAC in LCK T316I cells compared to LCK wildtype cells. Cells were treated with compounds for 72 hrs. Data represents mean ± SD from three biological replicates. **C**, Representative immunoblots showing LCK protein levels in KOPT-K1 with LCK WT and with LCK T316I after treatment with compound **17** at 1000 nM or SJ11646 at 10 nM for 24 hrs. Quantification of LCK degradation from three biological replicates normalized to DMSO-treated controls, in parental and dasatinib-resistant KOPT-K1 cells are shown below. **D**, LCK structural model showing that binding locations to dasatinib (cyan) and to LCK-MGDs (e.g., compound **1**, green) are different. The C-alpha of G415 is highlighted as a dark green sphere.

### Effects of LCK MGD on common IMiD neosubstrates

To explore the selectivity of our LCK MGDs, we used HEK293 cell line models in which HiBiT was tagged with either of two common neosubstrates, namely GSPT1 and CK1α^20^. Degradation of GSPT1 was not observed for compounds **16–19** (**Table S1**), while CK1α was degraded with DC_50_ ranging from 10 to 40 nM (**Table S1**). Immunoblot analyses also confirmed substantial CK1α degradation in KOPT-K1 cells treated with MGDs **17** and **19** (**Fig. 5A**). As shown in **Fig. 5B**, there is indeed significant structural similarity between LCK and CK1α. On the basis of DC_50_ for either LCK or CK1α, our series of MGDs clustered into two groups (LCK DC₅₀ ≥ or <100 nM, **Fig. 5C**). Importantly, compounds that degraded both LCK and CK1α (DC_50_ <100nM for both targets, n = 14) exhibited significantly greater cytotoxicity against KOPT-K1 cells compared to those preferentially degrading CK1α (n = 5, **Fig. 5D**).

**Fig. 5:**
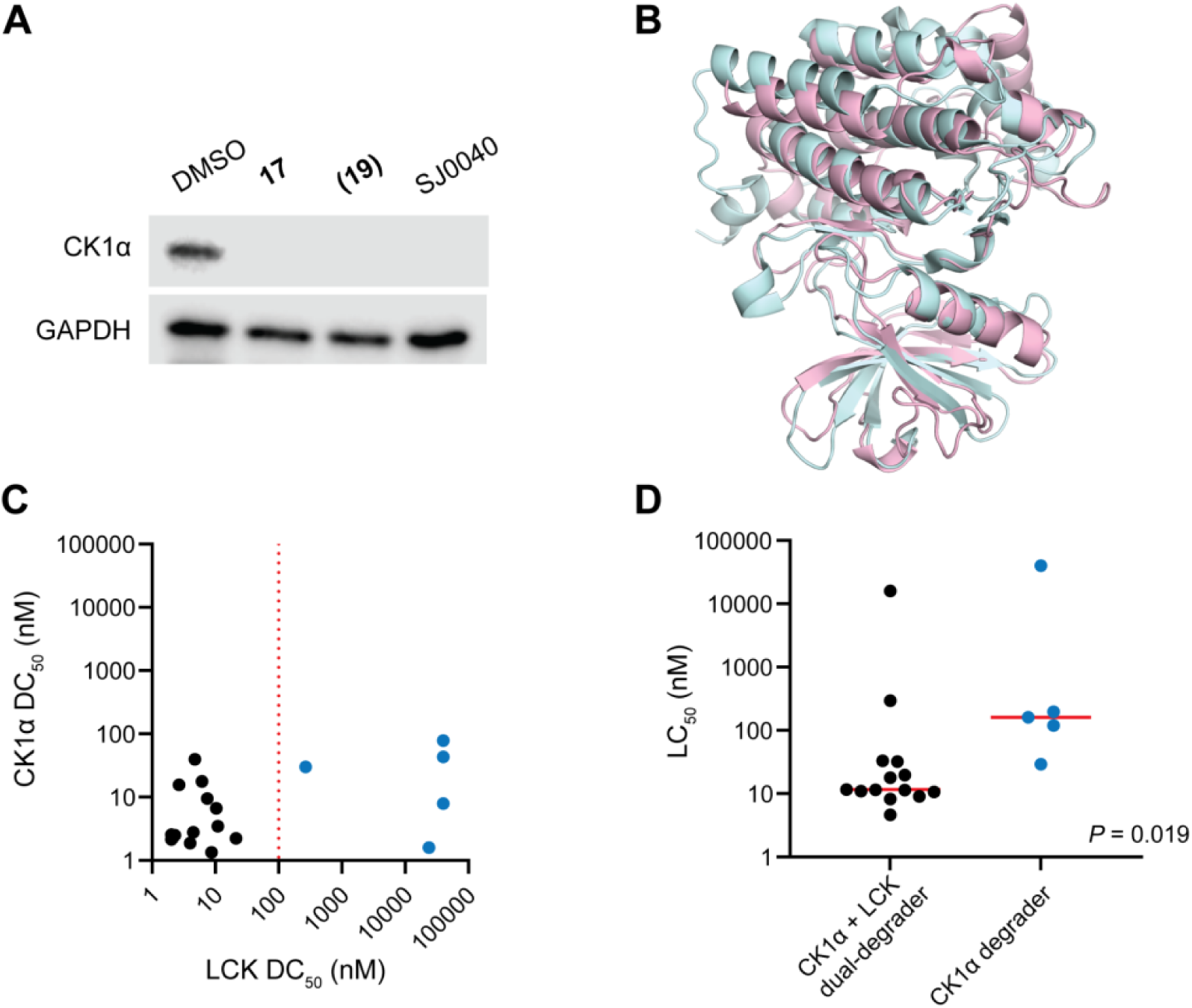
MGDs also show activity against common IMiD neosubstrates. **A**, Representative immunoblots showing CK1α degradation in KOPT-K1 treated with compounds **17** and **19**, or a selective CK1α degrader, SJ0040 (1000 nM, 24 hrs). **B**, Structural overlay illustrating similarity between the active form of LCK (pink) and CK1α (cyan). **C**, Degradation activity against LCK and CK1α by MGDs (**1-19**, excluding compounds **6** and **15**) and CK1α-selective degraders (SJ0040 and SJ3149), as determined by DC_50_ values at 24 hrs. The red dashed line indicates LCK DC_50_ threshold of 100 nM. **D**, Comparison of cytotoxicity in KOPT-K1 between MGDs capable of degrading both LCK and CK1α (DC_50_ <100 nM for both, n = 14) and those degrading CK1α (DC_50_ <100 nM for CK1α only, n = 5). Statistical significance was assessed by Mann-Whitney U test.

## DISCUSSION

Drug resistance is a formidable challenge in oncology. While targeted therapeutics offer improved therapeutic indices, they are often undermined by rapid emergence of resistance, often through mutations in the drug target^4,22^. Degradation-based agents, such as PROTACs and MGDs, have superior pharmacodynamic effects thanks to catalyst-like behavior^2,8,23^, but the patterns of drug resistance in this context remain unclear. In this study, we report on the identification and optimization of cereblon-recruiting MGDs that degrade LCK, an essential oncogenic kinase in T-ALL. Importantly, we provide a direct comparison of drug resistance for inhibitor, PROTAC, and MGD for the same target. The T316I mutation in LCK abrogates dasatinib binding and confers resistance both to the inhibitor itself and to dasatinib-based PROTACs. By contrast, our lead MGDs engage LCK through a distinct surface interaction at the helical G-loop degron and maintain activity in T316I-expressing cells. This confirms that MGDs can bypass resistance mechanisms that limit the efficacy of both small-molecule inhibitors and PROTACs based on the same scaffold.

Our data adds important evidence for degron space targetable by MGDs beyond the canonical β-hairpin G-loop motif, as proposed by Petzold et. al^21^. Despite their similarity in sequence and structure, the region in LCK corresponding to the β-hairpin degron of CK1α adopts a different inverted-loop conformation that is predicted to sterically prevent ternary complex formation (**Fig. 5B**). By contrast, our modeling analyses described a helical G-loop that likely mediates the LCK-MGD-CRBN interaction, as confirmed by the G415N mutation studies (**Fig. 3A**). These results will motivate further exploration of “non-canonical” degron motifs (such as helical G-loop) that are targetable by MGDs, leading to the expansion of neosubstrate space for TPD. Given the importance of G-loop engagement in determining specificity, refinement of MGDs to fit this interface may enable selective recognition of different neosubstrates^24^.

There are significant structural similarities between LCK and CK1α (**Fig. 5B**), and therefore it was not entirely surprising that our lead MGDs can also potently degrade CK1α, a therapeutic target in hematologic malignancies in its own right (dependency data from DepMap in **Fig. S3**)^25,26,27,28^. We reason that there may be several potential advantages for a dual degrader over selectively targeting each one individually. First, CK1α dependency is more common than the dependency on LCK in ALL^28^, therefore compounds targeting both would be more broadly active than LCK-specific agents (**Fig. 5D**). Second, in ALL cases for which both LCK and CK1α are essential, they represent two independent therapeutic vulnerabilities and simultaneously targeting non-overlapping pathways is advantageous because acquiring resistance mutations in two unrelated targets is much more difficult than developing resistance for one target^29,30^. Future optimization may focus on LCK selectivity, targeted delivery (e.g., antibody conjugation), or leveraging biomarker-guided strategies to mitigate toxicity while maintaining anti-leukemic potency.

In summary, we demonstrate that LCK-targeting MGDs represent a viable strategy to overcome clinically relevant drug resistance in T-ALL. This work highlights the potential of degrader-based therapeutics to complement or replace conventional kinase inhibitors, especially in settings where mutation-driven resistance is common.

## METHODS

### Cell Lines

T-ALL cell line KOPT-K1 was gifted by Dr. Takaomi Sanda at National University of Singapore. KOPT-K1 with HiBiT-tagged LCK was generated using CRISPR-Cas9 technology, as previously reported^17,31^. Dasatinib-resistant clones with a heterozygous LCK T316I mutation was generated as described previously^14^. LCK G415I knock-in clones were generated in KOPT-K1 cells by CRISPR-Cas9 editing for which genotyping primers and gRNAs are shown in **Table S2**. All T-ALL cell lines were cultured in RPMI-1640 supplemented with 10% FBS.

### Compound screening

3,424 compounds from the St. Jude Children’s Research Hospital proprietary molecular glue library were screened against an LCK-activated T-ALL line, KOPT-K1, expressing HiBiT-tagged LCK, as previously described^17^. Compounds were prepared in 384-well plates at a concentration of 10 mM in 100 nL. The cells were suspended at a density of 300,000 cells/mL, and 25 μL (7,500 cells) of the suspension were plated into each well using a VIAFLO 384 electronic pipette (Integra). Following incubation of 4 or 24 hrs at 37°C/5% CO_2_, the levels of HiBiT tagged-LCK protein were quantified by the Nano-Glo HiBiT lytic detection system (Promega, #N3030) per the manufacturer’s instruction. Luminescence was measured using a Synergy H4 Hybrid Microplate reader (BioTek). %LCK degradation was calculated by normalizing luminescence values to DMSO-treated controls and subtracting background signal from parental KOPT-K1 cells.

### LCK MGDs synthesis

Unless otherwise specified, reagents and solvents were obtained from commercial suppliers and used without further purification. Reactions were typically set up under ambient conditions and conducted under a nitrogen atmosphere. Thin-layer chromatography was performed using either Merck Millipore Silica Gel 60G F₂₅₄ glass plates or Biotage KP-NH plates and visualized under 254-nm ultraviolet light. Automated flash column chromatography was conducted on a Biotage SP1 system using Sfar Silica, Sfar Amino, or SNAP C18 columns. Solvent removal was carried out using a Büchi Rotovapor R-205. Nuclear magnetic resonance spectra were recorded on either a Bruker 400-MHz, 500-MHz, or 600-MHz spectrometer in the solvents indicated. Spectra were processed using MestReNova (v14), with chemical shifts reported in parts per million (ppm) relative to the residual solvent peak. Details for the synthesis of all LCK MGDs are provided in **Supplementary Materials**. Previously reported selective CK1α degraders SJ3149 and 0040^20^, as well as the dasatinib-based PROTAC SJ11646, were included in comparative assays^16^.

### LCK HiBiT assay following the screening

Compounds were prepared in 384-well plates in a dose-response format, with 100 nL of each compound dispensed at serial concentrations ranging from 0.5081 to10,000 μM. Cell incubation with compounds and detection of HiBiT-LCK levels were performed as described above. The DC_50_ was determined by the compound concentration at which %LCK degradation was 50%, using a four-parameter dose-response model. Each compound was tested independently in triplicates. When necessary, DC₅₀ values falling outside the tested concentration range were imputed as either the lowest or highest concentration tested.

### Ternary complex assay (LCK/CRBN-DDB1)

*Protein Production.* GST-tagged LCK construct (residues Gly2–Pro509) was designed, with the GST tag added to the N-terminus of LCK using a ‘SDGGGGS’ linker. The codon-optimized LCK gene fragment was synthesized and cloned into the pFastBac1 vector for expression in Sf9 insect cells. The protein was subsequently produced by Genscript. The GST-LCK(G415N) mutant protein construct was generated by site-directed mutagenesis of the GST-LCK construct, and the protein was also produced by Genscript. The purified proteins were stored in a final buffer containing 50 mM Tris-HCl, 500 mM NaCl, 5% glycerol, pH 8.0. His-tagged CRBN-DDB1 protein was prepared following the procedure reported by Matyskiela et al^32^.

*AlphaScreen Assay.* All reagents were diluted in assay buffer comprising 25 mM HEPES, pH 7.4, 100 mM NaCl, 0.1% BSA, and 0.05% Tween20. An ECHO 650 (Labcyte Inc.) acoustic dispenser was used to generate a 10-point dilution curve from DMSO stocks of the compounds directly into a 384-well OptiPlate (PerkinElmer, cat# 6007290) giving a final DMSO concentration of 0.1%. The assay mixture contained 100 nM His-tagged CRBN-DDB1, and 75 nM GST-tagged LCK (or GST-tagged LCK(G415N)). AlphaScreen glutathione coated donor and AlphaScreen nickel chelate acceptor beads were purchased from PerkinElmer (cat# 6765300 and 6760141 respectively). Briefly, to a 384-well OptiPlate containing 5× compound in triplicate was added 5 µL of a 5 × solution of His-CRBN-DDB1 and GST-LCK (or GST-LCK(G415N)) and then incubated at rt for 1 h. After incubation, 10 μL nickel chelate acceptor (20 µg/mL final concentration) and 10 μL glutathione donor beads (20 µg/mL final concentration) were added under subdued lighting. The plate was sealed and mixed on a MixMate (eppendorf) for 1 h at rt and then luminescence detection was collected on an Envision plate reader (PerkinElmer). A four-parameter dose-response model was used to fit the data using GraphPad Prism software (v10.3.1).

### Cell viability assay for KOPT-K1

Parental KOPT-K1, a previously reported dasatinib-resistant KOPT-K1 line harboring a heterozygous LCK T316I mutation^14^ were plated at a density of 120,000 cells/mL in 25μL onto 384 well plates containing compounds as described in the “**LCK HiBiT Assay following the screening”** section above. After 72 hrs of incubation at 37°C/5% CO_2_, cell viability was assessed using CellTiter-Glo® (CTG) assay (Promega, #G9243). An equal volume of CTG reagent was added to each well, followed by a brief incubation at room temperature. Luminescence was then measured using a Synergy H4 Hybrid Microplate reader (BioTek). Viability was calculated by normalizing luminescence values from compound-treated cells to that of DMSO-treated controls. Each compound was tested independently in triplicate, and LC_50_ values (the concentrations lethal to 50% of cells) were calculated by a four-parameter dose-response model.

### Immunoblotting

T-ALL cell lines were incubated for one hour and then DMSO or each compound diluted in DMSO was added. After being treated for 24 hrs, cells were harvested, washed once with ice-cold PBS and lysed with RIPA Lysis and Extraction Buffer (Thermo Scientific, #89901) supplemented with protease and phosphatase inhibitor cocktail (Thermo Scientific, #78440). Protein lysates were incubated on ice with gentle shaking for 15 minutes before being centrifuged at 4°C, 14,000 rpm for 15 minutes. Supernatants were added to an equal volume of 2 × Laemmli sample buffer (Bio-Rad, #1610737) supplemented with 2-mercaptoethanol (Bio-Rad, #1610710). Protein samples were heated at 98°C for 10 minutes before being stored at -20°C or western blotting. Equal amounts of protein samples were separated by precast 4–15% Tris-glycine Mini-PROTEAN TGX gels (Bio-Rad, #4561086). Resolved proteins were transferred onto Immobilon-FL PVDF membranes (Millipore, #IPFL00010) for detection of LCK and onto TransBlot Turbo LF PVDF membranes (Bio-Rad, #10026834) for detection of CK1α. For LCK, membranes were blocked with Intercept® (TBS) blocking buffer (LI-COR, #927-60001) for two hours at room temperature and then probed with primary antibodies at an optimal concentration in the same blocking buffer supplemented with 0.2% Tween 20 (Fisher BioReagents, #BP337-500) overnight at 4°C: anti-LCK (Cell Signaling Technology, #2657; 1:2000) and anti-GAPDH (Cell Signaling Technology, #2118; 1:5000) for loading controls. The membranes were then washed with TBS-T three times and incubated with the IRDye® 800CW goat anti-rabbit (LI-COR, #926-32211; 1:5000) and 680CW goat anti-mouse IgG secondary antibodies (LI-COR, #926-68070; 1:5000) in the blocking buffer with 0.2% Tween 20 and 0.02% sodium dodecyl sulfate at room temperature for two hours. Excessive antibodies were washed out with TBS-T, and the membranes were exposed in LI-COR Odyssey imaging system. Fluorescent intensity was quantified and analyzed with Image Studio software (lite version 5.2). For CK1α detections, membranes were blocked in 5% non-fat milk in TBS-T and probed with anti-CK1α (Abcam, #AB206652; 1:2000) followed by incubation with anti-rabbit HRP-lined IgG (Cell Signaling Technology, #7074; 1:5000). Chemiluminescence was detected using the LI-COR Odyssey imaging system. After detection, membranes were stripped from the membranes with stripping buffer and re-probed with anti-GAPDH as a loading control.

### Modeling of ternary complex and pocket volume analysis

Compounds (e.g., (**1**)SJ1581) were docked into the MG binding pocket to CRBN only extracted from the crystal structure of DDB1-CRBN-SJ3149-CK1α (8G66)^20^. Ligands were prepared using Schrödinger LigPrep^33^ (OPLS3e^34^ force field for minimization and Epik^35^ for protonation with pH=7±2); the receptor grid was prepared using Schrödinger protein preparation workflow^36^ (OPLS3e force field for minimization and grid based on SJ3149 position); and docking was performed using Glide SP^37^. To incorporate LCK into the model, an inactive structure of LCK (4C3F) was superimposed into the hypothesized position^38^ by aligning the 3 backbone carbonyl oxygens from the LCK helical degron (G[415]-1, G-2, G-3) to the CK1α loop degron backbone carbonyl oxygens (G[40]-1, G-2, G-3). Rosetta^39^ was then used to relax (minimize) the overall structure 250 times, resulting in an ensemble of potential models that were used to understand SAR. Finally, the pocket volume (centered around the Cl coordinate in 1) was calculated using POVME^40^ in each of the 250 relaxed models.

### CK1α HiBiT assay

For the CK1α HiBiT assay, CSNK1A1 (CK1α)-HiBiT HEK293 LgBiT KI cells (Promega, Cat. #CS3023104) were seeded at 5,000 cells/well in white 384-well assay plates (Corning, Cat. # 3570BC) and incubated overnight at 37 °C, 5% CO_2_. After overnight incubation, compounds were added in a dose-response format using an Echo 655 Acoustic Liquid Handler (Beckman Coulter). After a 24-hr incubation with compounds, the level of HiBiT tagged-CK1α was assessed by adding Nano-Glo Live Cell Assay reagent (Promega, Cat. #N2013) per manufacturer instructions. Plates were incubated for at least 10 minutes before reading the luminescence signal on an EnVision plate reader (PerkinElmer). The assay was conducted with two biological replicates. Luminescence data were analyzed using an in-house pipeline, Robust Investigation of Screening Experiments (RISE), based on the Pipeline Pilot platform (Accelrys, v.8.5.0), to determine compound DC_50_ values using a four-parameter dose-response model.

### GSPT1 HiBiT assay

The GSPT1 HiBiT assay used HEK293 hGSPT1 HiBiT tagged cells. Cells were seeded at 1,000 cells/well in white 384-well assay plates (Corning, Cat. # 3570BC) in duplicate and incubated overnight at 37 °C and 5% CO_2_. After overnight incubation, compound treatment was performed using an Echo 655 Acoustic Liquid Handler (Beckman Coulter) in a dose-response format. Following a 24-hr incubation, HiBiT-GSPT1 levels were measured by adding the Nano-Glo HiBiT lytic detection system (Promega, Cat. #N3050) according to manufacturer instructions. Plates were incubated for at least 10 minutes before reading the luminescence signal using an EnVision plate reader (PerkinElmer). Luminescence data analysis was performed similarly to the CK1α HiBiT data using the RISE data analysis platform to determine compound DC_50_ values with a four-parameter dose-response model.

### Statistical analysis

The association of LCK DC_50_ values between 4hrs and 24hrs was evaluated by Wilcoxon matched-pairs signed rank test. The comparison of LC_50_ values in KOPT-K1 between MGDs capable of degrading both LCK and CK1α and those degrading CK1α only was assessed by Mann-Whitney U test. *P*-values were considered significant if <0.05. All analyses were performed with GraphPad Prism (version 10.1.2).

## ACKNOWLEDGEMENTS

We thank the Hartwell Center for Bioinformatics and Biotechnology and the Center for Advanced Genome Engineering (S. Pruett-Miller and M. Reynolds-Gagliano) at St. Jude Children’s Research Hospital for their technical support in performing experiments included in this study. This work was in part supported by the National Institutes of Health (P30 CA021765, R01CA264837), Alex’s Lemonade Stand Foundation Crazy8 grant, and American Lebanese Syrian Associated Charities.

J.J.Y. is the Endowed Chair of Pharmacogenomics at St. Jude Children’s Research Hospital.

## AUTHOR CONTRIBUTIONS

J.J.Y., Z.R, and G.N. conceptualized and supervised the project. M.F. led the structural and computational biology efforts. S.Y. conducted the compound library screening. M.A. and J.A.J. synthesized all MGDs based on the hit from the screening. S.Y., M.A., A.L., A.A., and A.M. performed *in vitro* characterizations of MGDs following the initial screening. J.T.S. performed structural modeling. Y.L., D.G.L., and L.Y. characterized the physicochemical properties of MGDs. J.J.Y., G.N., S.Y., M.A., M.F., and J.T.S. wrote the manuscript. All authors reviewed and approved the final version of the manuscript.

