## Supplementary Materials for "LCK-targeting molecular glues overcome resistance to inhibitor-based therapy in T-cell acute lymphoblastic leukemia"

**Table of Contents**

Supplementary Tables	2

Supplementary Figures	3

*In vitro* ADME assay methods	5

Synthetic methods and schemes	6

^1^H and ^13^C spectra figures for new key compounds	30

LC chromatograms for purity assessment of compounds used *in vitro*	68

Supplementary References	86

**SUPPLEMENTARY TABLE**

**Table S1.** Degradation potency of compounds **(16)**–(**19)** for GSPT1 and CK1α in comparison with LCK

| **ID** | **GSPT1 DC_50_ (nM)^a^** | **CK1α DC_50_ (nM)^a^** | **LCK DC_50_ (nM)^b^** |
| --- | --- | --- | --- |
| (**16**) | >10000 | 39.7 ± 1.9 | 4.7 ± 1.6 |
| (**17**) | >10000 | 17.8 ± 1.6 | 6.2 ± 1.5 |
| (**18**) | >10000 | 9.5 ± 1.4 | 7.5 ± 1.6 |
| (**19**) | >10000 | 15.6 ± 0.9 | 2.7 ± 0.6 |

^a^DC_50_ potency was calculated using HiBiT assays after 24 hrs of drug-exposure.

**Table S2.** Primers used in this study

| LCK-G415-gDNA-F | ATTCATCGTGACCTTCGGGC |
| --- | --- |
| LCK-G415-gDNA-R | ATACGGAGATGGAAGCCAGC |
| gRNA-LCK-G415N | GGTGAATGTCCCGTAGTTAA |

**Table S3.** Ternary complex formation and R group volume for 3-substituted indoles.

| **ID** | **Ternary EC_50_ (nM)^a^** | 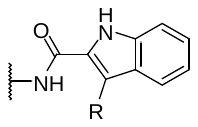  **Volume of R group (Å^3^, relative to R=H)** |
| --- | --- | --- |
| **2** | 109.9 (97.8 – 123.3) | 11.2 (R = Me) |
| **3** | 234.7 (163.5 – 347.9) | 0 (R = H) |
| **7** | 97.4 (85.4 – 110.9) | 23.0 (R = Et) |
| **8** | 194.5 (182.7 – 207.2) | 37.8 (R = Pr) |
| **20** | 191.1 (154.1 – 240.4) | 35.1 (R = iPr) |
| **21** | 290.6 (237.3 – 355.5) | 33.8 (R = cycloPr) |
| **22** | 574 (470.5 – 702.9) | 61.1 (R = Ph) |
| **23** | 1431 (839.2 – 4472) | 74.9 (R = Bn) |

^a^AlphaScreen ternary complex formation half-maximum concentration reported as the mean with 95% confidence interval (CI) for three technical replicates.

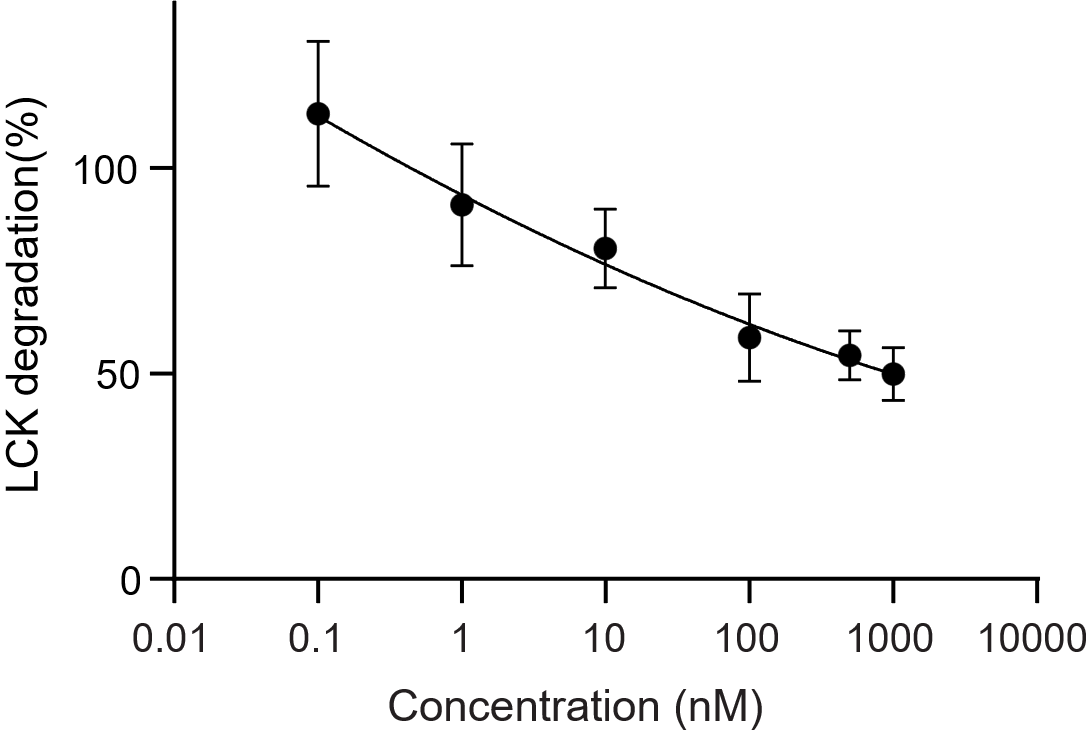
**SUPPLEMENTARY FIGURES**

**Figure S1.** LCK degradation by compound **1** determined by immunoblotting. KOPT-K1 LCK HiBiT cells were treated with compound **1** at the indicated serial concentrations for 24hrs, followed by immunoblotting. LCK levels were quantified by fluorescent intensity and LCK/GAPDH ratios were normalized to those of DMSO-treated control. The DC_50_ for LCK degradation was determined as the compound concentration at which %LCK degradation was 50%, calculated using a four-parameter dose-response model.

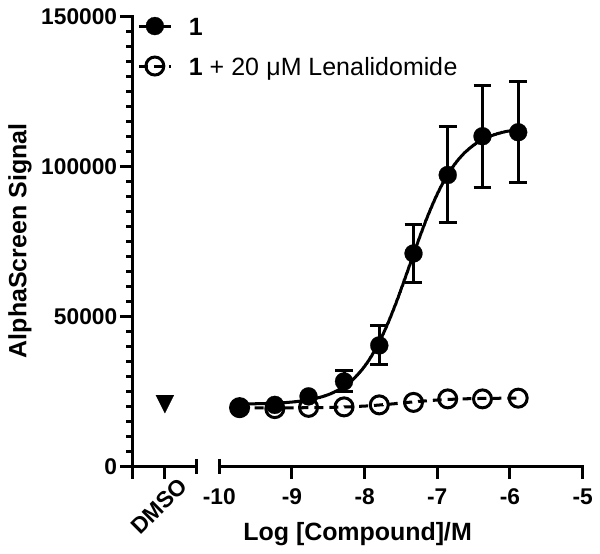

**Figure S2.** Ternary complex formation of compound **1** in the presence and absence of lenalidomide. MGD-induced formation of a ternary complex between GST-LCK and His-CRBN-DDB1 proteins measured using an *in vitro* AlphaScreen assay. Data is shown as mean ± SD from three technical replicates.

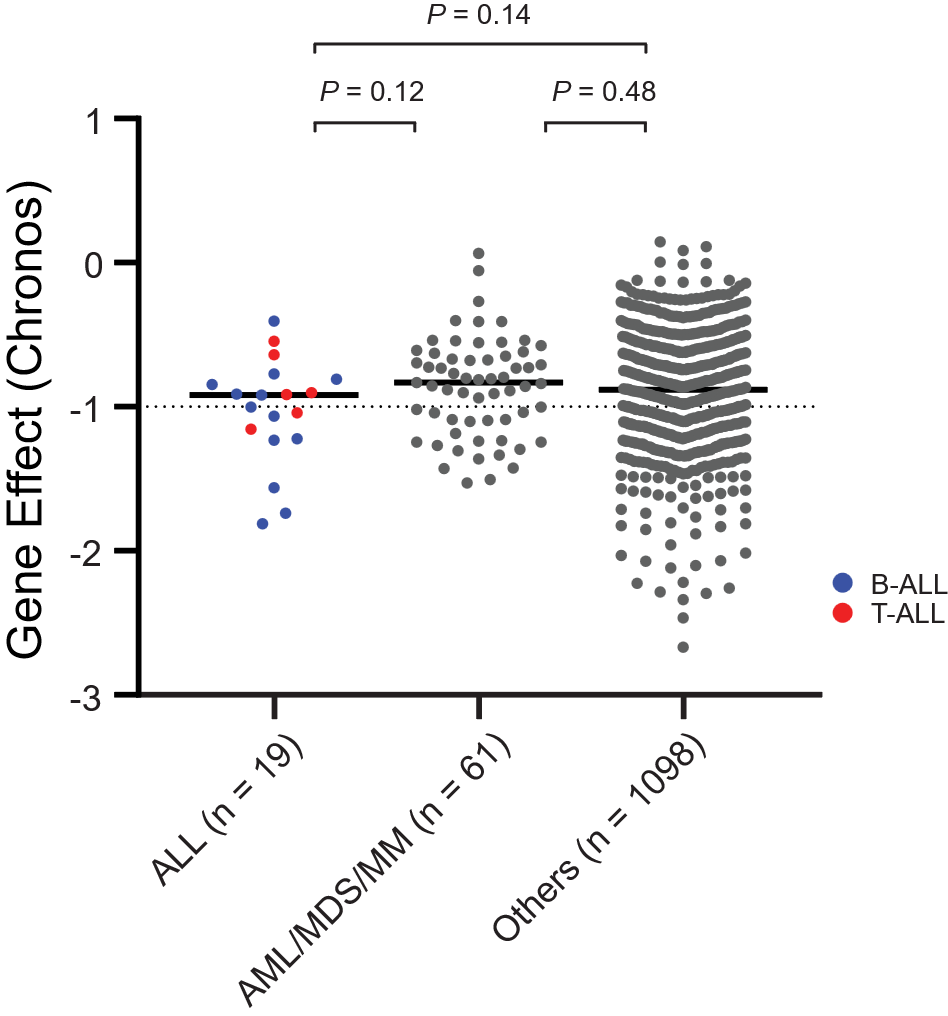

**Figure S3.** Dependency on CK1α in ALL cell lines in comparison with AML, MDS, or MM. Gene Effect (Chronos) scores across a total of 1178 cell lines were obtained on the DepMap portal^1^: https://depmap.org/portal. For ALL cell lines (n=19), B-ALL are indicated in blue, while red dots represent T-ALL cell lines. 53.8% of B-ALL (n = 7 of 13) and 33.3% of T-ALL lines (n = 2 of 6) were considered dependent on CSNK1A (Gene Effect < −1.0). Statistical significance was assessed by Mann-Whitney U test using Prism GraphPad.

***IN VITRO* ADME ASSAY METHODS**

Instrumentation

**UPLC/MS/UV System:** Chromatographic separation was conducted using a Waters Acquity ultra-performance liquid chromatography system coupled to an SQ mass spectrometer. The UPLCSQDMS conditions were set similarly to those reported previously^2^.

**UHPLC-MS/MS System:** Chromatographic separation was conducted using an Acquity UPLC BEH C18 column (1.7 µm, 2.1 x 50 mm; Waters Corporation, Milford, MA) on a Sciex ExionLC™ system with a 6500+ Qtrap mass spectrometer (SCIEX, Forster City, CA). The UPLC column was maintained at 55 °C. Mobile phase A was 0.1% formic acid in MilliQ H_2_O and Mobile phase B was 0.1% formic acid in acetonitrile. The flow rate was 0.9 mL/min with a gradient of 0–0.2 min, B% 1–1%; 0.2–0.5min, B% 1–50%; 0.5–1.6 min, B% 50–95%; 1.6–1.95 min, B% 95–95%; 1.95–1.96 min, B% 95–1%; and 1.96–2.2 min, B% 1–1%. The first 0.4 minutes of eluate were desalted to waste via an integrated Valco valve. The remaining eluate was directed to the triple quadrupole mass spectrometer equipped with an electrospray ionization source. LC-MS/MS was performed in positive polarity (5000 V), with a source temperature of 550 °C. Nitrogen was used for Gas 1 and Gas 2, set at 60. The curtain gas and collision gas, also nitrogen, were set to 40 and 6, respectively. Multiple reaction monitoring transitions and energy parameters for each compound and control were selected using Discovery Quant Software. Data acquisition was managed with SCIEX Analyst 1.7.3 and data processing was performed with SCIEX MS-OS 2.1.1 software.

Aqueous solubility

Solubility assays were conducted using a Biomek FX lab automation workstation (Beckman Coulter, Inc., Fullerton, CA) and μSOL Evolution software (pION Inc., Woburn, MA). In a 96- well microplate (Cat. No: 3363, Corning Incorporated, Salt Lake, UT), 10 μL of a 10 mM test compound stock in DMSO was added to 190 μL 1-propanol (Thermo Scientific, spectroscopy, Cat. No: 434360010, Fair Lawn, NJ) to create a reference stock plate. From this reference stock plate, 5 μL solutions were mixed with 70 μL 1-propanol and 75 μL Dulbecco's Phosphate Buffered Saline (DPBS, 1X, Gibco™, Cat. No: 14190-144, Thermo Fisher Scientifics, Waltham, MA) to generate the reference plate. In a 96-well storage plate (Cat. No: 201276-100, Agilent, Santa Clara, CA), 6 μL of a 10 mM test compound stock was added to 600 μL buffer, mixed, sealed, and incubated at room temperature for 18 hours. Following incubation, the suspension was centrifuged at 4000 rpm under room temperature (Eppendorf 5910R Refrigerated Centrifuge) for 20 minutes. The supernatant (75 μL) was combined with 75 μL 1-propanol to create the sample plate. Concentrations in reference and sample plates were assessed by UPLC/MS/UV system. Solubility (μM) were determined via the equation, Solubility = (Peak Area_Sample_/3) x C_0_/ Peak Area_reference_. C_0_ is the concentration of DMSO stock solution divided by 100. All compounds were tested in triplicate wells.

**SYNTHETIC METHODS**

Abbreviations used: DCM, dichloromethane; DIPEA, diisopropylethylamine; DMF, dimethylformamide; HATU, hexafluorophosphate azabenzotriazole tetramethyl uranium; MTBE, methyl *tert*-butyl ether; NMP, *N*-methyl-2-pyrrolidone; TCFH, chloro-*N,N,N′,N′*-tetramethylformamidinium hexafluorophosphate; THF, tetrahydrofuran; XPhos, 2-dicyclohexylphosphino-2′,4′,6′-triisopropylbiphenyl.

**General Methods and Synthesis.** No unexpected or unusually high safety hazards were encountered while carrying out the experimental procedures described. Reagents and solvents were used as received from commercial suppliers without purification unless described otherwise. Reactions were set up under air and carried out under nitrogen atmosphere in a fume hood unless specifically specified otherwise. Evaporation under reduced pressure was carried out using a Büchi Rotovapor R-205 with a B-490 heating bath base and V-700 vacuum pump. Thin layer chromatography to monitor reaction progress was performed using Merck Millipore silica gel 60G F254 glass plates or Biotage KP-NH plates and a handheld 254 nm UV lamp for detection. Automated flash column chromatography was carried out on a Biotage Isolera One flash chromatography system using Biotage Sfar silica, amino, or C18 columns. NMR spectra were acquired on Bruker 400 MHz, 500 MHz, or 600 MHz spectrometers in the solvents indicated. NMR spectra were processed using MestReNova v14. Chemical shifts (*δ*) are reported in parts per million (ppm) referenced to the indicated solvent peak with signals designated as: s, singlet; br d, broad doublet; br dd broad doublet of doublets; br s, broad singlet; d, doublet; dd, doublet of doublets; dt, doublet of triplets; dq, doublet of quartets; ddd, doublet of doublet of doublets; t, triplet; td, triplet of doublets; tdd, triplet of doublet of doublets; q, quartet; qd, quartet of doublets; p, pentet; m, multiplet. Coupling constants (*J*) are in Hertz (Hz). Compound purity was assessed using a Waters Corp. UPLC-MS (Acquity PDA detector, Acquity SQ detector and Acquity UPLC BEH-C18 1.7 µm, 2.1 x 50 mm column) with a mobile phase of 0.1% formic acid in water and acetonitrile. All compounds are >95% pure by UPLC by average of UV/ELSD signals, unless otherwise specified.

**High Resolution Mass Spectrometry Analysis.** Chromatographic separation was carried out using an ACQUITY UPLC™ BEH C18 column (2.1 × 50 mm, 1.7 µm particle size; Waters Corporation, Milford, MA) on an ACQUITY UPLC system. Data acquisition was performed using Waters_Connect Service version 7.9.0 software, with the system interfaced to a Waters Xevo Multi-Reflecting Time-of-Flight (MRT) mass spectrometer. The mobile phase flow rate was set at 0.6 mL/min, and the column temperature was maintained at 63 °C. A gradient elution was applied as follows: 99% solvent A (0.1% formic acid in water) was held for 0.3 min, followed by a linear ramp to 99% solvent B (0.1% formic acid in acetonitrile) over 1.7 min. This condition was held for 0.85 min, then returned to 99% A over 0.05 min and held for an additional 0.1 min. To minimize salt interference, the sample flow was diverted to waste during the first 0.1 min, after which it was directed to the mass spectrometer until 2.9 min. Mass spectrometry was conducted in positive-ion mode using electrospray ionization (ESI). The following parameters were applied: capillary voltage, 0.5 kV; cone voltage, 20 V; source temperature, 120 °C; desolvation temperature, 600 °C; desolvation gas flow, 1000 L/h; and cone gas flow, 50 L/h. Data acquisition was performed in MSe mode with a low collision energy of 6 V and a high-energy range from 20 to 55 V. Intelligent data capture was enabled and set to "high." Mass spectra were collected over an *m/z* range of 50–1200 with a scan time of 0.2 s. Leucine-enkephalin was used as a lock-mass reference for internal calibration. Lock-mass acquisition was performed in single mode with a capillary voltage of 3.0 kV, cone voltage of 20 V, and collision energy of 4 V, with scans acquired every 0.25 min. A mass accuracy of <5 ppm was maintained throughout the analysis.

**Scheme S1. Preparation of Compounds 1–5, 7, 21-23^a^**

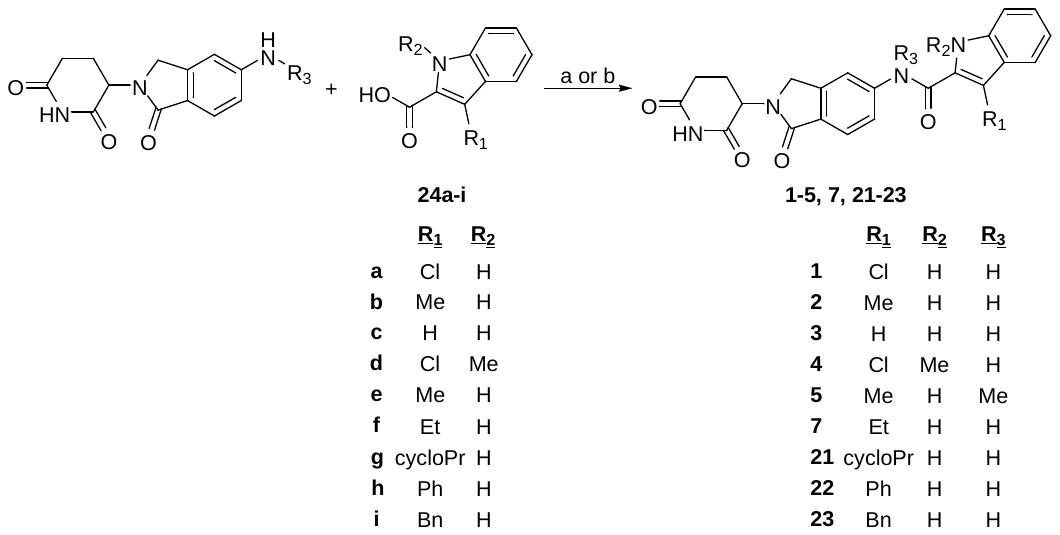

^a^Reagents and conditions: a) appropriate carboxylic acid **24** (1.0 equiv), HATU (1.5 equiv), DIPEA (2.0 equiv), DMF, rt, 16 h; b) appropriate carboxylic acid **24** (1.0 equiv), TCFH (2.0 equiv), *N*-methylimidazole (3.0 equiv), MeCN or NMP, rt, 3–4 h.

***Synthesis of 3-chloro-N-(2-(2,6-dioxopiperidin-3-yl)-1-oxoisoindolin-5-yl)-1H-indole-2-carboxamide*** ***1***. 3-(5-Amino-1-oxoisoindolin-2-yl)piperidine-2,6-dione (30 mg, 0.116 mmol) was added to a stirring solution of 3-chloro-1*H*-indole-2-carboxylic acid **24a** (22.6 mg, 0.116 mmol), HATU (66.0 mg, 0.174 mmol), and DIPEA (40.4 µL, 0.231 mmol) in DMF (500 µL). The reaction mixture was stirred for 16 h and then diluted with water (5 mL) and extracted with EtOAc (3 × 5 mL). The combined organic layers were washed with brine, dried over anhydrous Na_2_SO_4_, filtered, and concentrated. The crude product was purified using automated flash chromatography (Sfar Silica HC D 10 g column, methanol/dichloromethane gradient mobile phase). Product-containing fractions were concentrated to obtain the desired product (2 mg, 4% yield). HRMS (ESI) for C_22_H_18_ClN_4_O_4_ [M + H]^+^ calc’d 437.1016, obtained 437.1011. ^1^H NMR (500 MHz, DMSO-*d_6_*) δ 12.17 (s, 1H), 11.00 (s, 1H), 10.41 (s, 1H), 8.11 (d, *J* = 1.7 Hz, 1H), 7.85 – 7.78 (m, 1H), 7.75 (d, *J* = 8.3 Hz, 1H), 7.63 (d, *J* = 8.1 Hz, 1H), 7.52 (d, *J* = 8.3 Hz, 1H), 7.36 (t, *J* = 7.7 Hz, 1H), 7.22 (t, *J* = 7.5 Hz, 1H), 5.10 (dd, *J* = 13.3, 5.1 Hz, 1H), 4.49 (d, *J* = 17.3 Hz, 1H), 4.35 (d, *J* = 17.2 Hz, 1H), 2.92 (ddd, *J* = 18.3, 13.7, 5.5 Hz, 1H), 2.61 (d, *J* = 17.4 Hz, 1H), 2.41 (tt, *J* = 14.4, 7.3 Hz, 1H), 2.05 – 1.97 (m, 1H). ^13^C NMR (126 MHz, DMSO-*d_6_*) δ 173.01, 171.17, 167.86, 158.80, 143.37, 141.76, 134.63, 127.12, 126.94, 125.37, 124.90, 123.83, 121.10, 119.78, 118.72, 114.27, 112.94, 106.11, 51.67, 47.30, 31.28, 22.57.

***Synthesis of N-(2-(2,6-dioxopiperidin-3-yl)-1-oxoisoindolin-5-yl)-3-methyl-1H-indole-2-carboxamide*** ***2***. 3-(5-Amino-1-oxoisoindolin-2-yl)piperidine-2,6-dione (30 mg, 0.116 mmol) was added to a stirring solution of 3-methyl-1*H*-indole-2-carboxylic acid **24b** (20.3 mg, 0.116 mmol), HATU (66.0 mg, 0.174 mmol), and DIPEA (40.4 µL, 0.231 mmol) in DMF (500 µL). The reaction mixture was stirred for 16 h and then diluted with water (5 mL) and extracted with EtOAc (3 × 5 mL). The combined organic layers were washed with brine, dried over anhydrous Na_2_SO_4_, filtered, and concentrated. The crude product was purified using automated flash chromatography (Sfar Silica HC D 10 g column, methanol/dichloromethane gradient mobile phase). Product-containing fractions were concentrated to obtain the desired product (11 mg, 22% yield). HRMS (ESI) for C_23_H_21_N_4_O_4_ [M + H]^+^ calc’d 417.1563, obtained 417.1563. ^1^H NMR (500 MHz, DMSO-*d_6_*) δ 11.45 (s, 1H), 11.00 (s, 1H), 10.24 (s, 1H), 8.13 (d, *J* = 1.8 Hz, 1H), 7.78 (dd, *J* = 8.3, 1.7 Hz, 1H), 7.73 (d, *J* = 8.2 Hz, 1H), 7.66 (d, *J* = 8.0 Hz, 1H), 7.45 (d, *J* = 8.2 Hz, 1H), 7.26 (t, *J* = 7.6 Hz, 1H), 7.09 (t, *J* = 7.5 Hz, 1H), 5.10 (dd, *J* = 13.3, 5.1 Hz, 1H), 4.48 (d, *J* = 17.2 Hz, 1H), 4.34 (d, *J* = 17.2 Hz, 1H), 2.92 (ddd, *J* = 18.0, 13.6, 5.4 Hz, 1H), 2.67 – 2.58 (m, 1H), 2.56 (s, 3H), 2.40 (qd, *J* = 13.2, 4.5 Hz, 1H), 2.04 – 1.94 (m, 1H). ^13^C NMR (126 MHz, DMSO-*d_6_*) δ 173.01, 171.20, 167.95, 160.98, 143.34, 142.37, 135.74, 128.02, 127.43, 126.63, 124.43, 123.74, 120.02, 119.57, 119.45, 115.99, 114.01, 112.12, 51.65, 47.28, 31.29, 22.59, 9.86.

***Synthesis of N-(2-(2,6-dioxopiperidin-3-yl)-1-oxoisoindolin-5-yl)-1H-indole-2-carboxamide*** ***3***. 3-(5-Amino-1-oxoisoindolin-2-yl)piperidine-2,6-dione (30 mg, 0.116 mmol) was added to a stirring solution of 1*H*-indole-2-carboxylic acid **24c** (18.7 mg, 0.116 mmol), HATU (66.0 mg, 0.174 mmol), and DIPEA (40.4 µL, 0.231 mmol) in DMF (500 µL). The reaction mixture was stirred for 16 h and then diluted with water (5 mL) and extracted with EtOAc (3 × 5 mL). The combined organic layers were washed with brine, dried over anhydrous Na_2_SO_4_, filtered, and concentrated. The crude product was purified using automated flash chromatography (Sfar Silica HC D 10 g column, methanol/dichloromethane gradient mobile phase). Product-containing fractions were concentrated to obtain the desired product (27 mg, 58% yield). HRMS (ESI) for C_22_H_19_N_4_O_4_ [M + H]^+^ calc’d 403.1406, obtained 403.1405. ^1^H NMR (500 MHz, DMSO-*d_6_*) δ 11.79 (s, 1H), 11.00 (s, 1H), 10.51 (s, 1H), 8.15 (d, *J* = 1.7 Hz, 1H), 7.89 (dd, *J* = 8.3, 1.8 Hz, 1H), 7.74 (d, *J* = 8.3 Hz, 1H), 7.70 (d, *J* = 8.0 Hz, 1H), 7.51 – 7.46 (m, 2H), 7.24 (t, *J* = 7.7 Hz, 1H), 7.08 (t, *J* = 7.5 Hz, 1H), 5.10 (dd, *J* = 13.3, 5.1 Hz, 1H), 4.49 (d, *J* = 17.1 Hz, 1H), 4.35 (d, *J* = 17.1 Hz, 1H), 2.92 (ddd, *J* = 18.0, 13.6, 5.4 Hz, 1H), 2.65 – 2.58 (m, 1H), 2.40 (qd, *J* = 13.3, 4.5 Hz, 1H), 2.02 (ddd, *J* = 10.9, 5.6, 3.4 Hz, 1H). ^13^C NMR (126 MHz, DMSO-*d_6_*) δ 173.02, 171.19, 167.95, 160.07, 143.26, 142.33, 137.05, 131.14, 127.02, 126.65, 124.14, 123.70, 121.95, 120.12, 119.77, 114.25, 112.52, 104.55, 51.64, 47.28, 31.29, 22.60.

***Synthesis of 3-chloro-N-(2-(2,6-dioxopiperidin-3-yl)-1-oxoisoindolin-5-yl)-1-methyl-1H-indole-2-carboxamide*** ***4***. 3-(5-Amino-1-oxoisoindolin-2-yl)piperidine-2,6-dione (30 mg, 0.116 mmol) was added to a stirring solution of 3-chloro-1-methyl-1*H*-indole-2-carboxylic acid **24d** (24.3 mg, 0.116 mmol), HATU (66.0 mg, 0.174 mmol), and DIPEA (40.4 µL, 0.231 mmol) in DMF (500 µL). The reaction mixture was stirred for 16 h and then diluted with water (5 mL) and extracted with EtOAc (3 × 5 mL). The combined organic layers were washed with brine, dried over anhydrous Na_2_SO_4_, filtered, and concentrated. The crude product was purified using automated flash chromatography (Sfar Silica HC D 10 g column, methanol/dichloromethane gradient mobile phase). Product-containing fractions were concentrated to obtain the desired product (12 mg, 23% yield). HRMS (ESI) for C_23_H_20_ClN_4_O_4_ [M + H]^+^ calc’d 451.1173, obtained 451.1167. ^1^H NMR (500 MHz, DMSO-*d_6_*) δ 11.00 (s, 1H), 10.97 (s, 1H), 8.14 (d, *J* = 1.7 Hz, 1H), 7.79 (dd, *J* = 8.3, 1.8 Hz, 1H), 7.75 (d, *J* = 8.2 Hz, 1H), 7.65 (dd, *J* = 15.2, 8.2 Hz, 2H), 7.42 (ddd, *J* = 8.4, 6.9, 1.2 Hz, 1H), 7.27 (t, *J* = 7.5 Hz, 1H), 5.12 (dd, *J* = 13.3, 5.1 Hz, 1H), 4.49 (d, *J* = 17.3 Hz, 1H), 4.35 (d, *J* = 17.3 Hz, 1H), 3.89 (s, 3H), 2.92 (ddd, *J* = 17.1, 13.6, 5.4 Hz, 1H), 2.65 – 2.57 (m, 1H), 2.40 (qd, *J* = 13.2, 4.4 Hz, 1H), 2.02 (dtd, *J* = 12.7, 5.3, 2.3 Hz, 1H). ^13^C NMR (126 MHz, DMSO-*d_6_*) δ 172.91, 171.08, 167.74, 158.76, 143.38, 141.65, 136.00, 130.09, 127.29, 124.91, 123.84, 123.68, 121.21, 119.50, 118.55, 114.03, 111.14, 104.64, 51.61, 47.22, 31.46, 31.22, 22.53.

***Synthesis of N-(2-(2,6-dioxopiperidin-3-yl)-1-oxoisoindolin-5-yl)-N,3-dimethyl-1H-indole-2-carboxamide*** ***5***. *N*-Methylimidazole (26.3 µL, 0.329 mmol) was added to a stirring solution of 3-(5-(methylamino)-1-oxoisoindolin-2-yl)piperidine-2,6-dione (30 mg, 0.110 mmol), 3-methyl-1*H*-indole-2-carboxylic acid **24e** (19.2 mg, 0.110 mmol), and TCFH (61.6 mg, 0.220 mmol) in acetonitrile (500 µL). The reaction mixture was stirred for 4 h and then diluted with water (5 mL) and extracted with EtOAc (3 × 5 mL). The combined organic layers were washed with brine, dried over anhydrous Na_2_SO_4_, filtered, and concentrated. The crude product was purified using automated flash chromatography (Sfar Silica HC D 10 g column, methanol/dichloromethane gradient mobile phase). Product-containing fractions were concentrated to obtain the desired product (23 mg, 49% yield). HRMS (ESI) for C_24_H_23_N_4_O_4_ [M + H]^+^ calc’d 431.1719, obtained 431.1718. ^1^H NMR (500 MHz, DMSO-*d_6_*) δ 11.16 (s, 1H), 10.97 (s, 1H), 7.55 (d, *J* = 8.2 Hz, 1H), 7.51 (d, *J* = 1.8 Hz, 1H), 7.40 (d, *J* = 7.9 Hz, 1H), 7.29 (dd, *J* = 8.2, 0.9 Hz, 1H), 7.23 (dd, *J* = 8.2, 2.0 Hz, 1H), 7.12 (ddd, *J* = 8.2, 6.9, 1.1 Hz, 1H), 6.95 (ddd, *J* = 8.0, 6.9, 1.0 Hz, 1H), 5.05 (dd, *J* = 13.3, 5.1 Hz, 1H), 4.35 (d, *J* = 17.5 Hz, 1H), 4.23 (d, *J* = 17.5 Hz, 1H), 3.48 (s, 3H), 2.87 (ddd, *J* = 17.3, 13.6, 5.4 Hz, 1H), 2.61 – 2.53 (m, 1H), 2.34 (qd, *J* = 13.2, 4.5 Hz, 1H), 1.98 – 1.92 (m, 4H). ^13^C NMR (126 MHz, DMSO-*d_6_*) δ 172.97, 171.05, 167.46, 164.67, 147.38, 142.77, 135.92, 128.81, 128.56, 127.27, 125.79, 123.31, 123.26, 120.59, 119.54, 119.00, 111.80, 111.20, 54.97, 51.69, 47.12, 37.76, 31.23, 22.44, 9.44.

***Synthesis of N-(2-(2,6-dioxopiperidin-3-yl)-1-oxoisoindolin-5-yl)-3-ethyl-1H-indole-2-carboxamide*** ***7***. To a solution of 3-ethyl-1*H*-indole-2-carboxylic acid **24f** (0.022 g, 0.12 mmol) in DMF (0.5 mL) at rt was added DIPEA (0.060 mL, 0.35 mmol), HATU (0.048 g, 0.13 mmol) and 3-(5-amino-1-oxoisoindolin-2-yl)piperidine-2,6-dione (0.030 g, 0.12 mmol). The reaction mixture was stirred at room temperature for 16 h then was diluted with ethyl acetate (3 mL) and washed sequentially with brine (2 × 3 mL) followed by 5% aqueous LiCl solution (3 mL). The organic phase was dried over sodium sulfate, filtered and concentrated *in vacuo*. The crude was purified using automated flash chromatography (Biotage Sfar Amino D 11g column, methanol/dicholoromethane gradient mobile phase). Product-containing fractions were evaporated *in vacuo* to obtain a tan solid (8.3 mg, 17%). HRMS (ESI) for C_24_H_23_N_4_O_4_ [M+H]^+^ calc’d 431.1719, obtained 431.1733. ^1^H NMR (500 MHz, DMSO-*d_6_*) δ 11.44 (s, 1H), 10.99 (s, 1H), 10.29 (s, 1H), 8.15 (s, 1H), 7.79 – 7.70 (m, 2H), 7.68 (d, *J* = 8.1 Hz, 1H), 7.46 (d, *J* = 8.2 Hz, 1H), 7.31 – 7.23 (m, 1H), 7.13 – 7.03 (m, 1H), 5.11 (dd, *J* = 13.3, 5.1 Hz, 1H), 4.54 – 4.26 (m, 2H), 3.09 (q, *J* = 7.4 Hz, 2H), 2.97 – 2.86 (m, 1H), 2.66 – 2.57 (m, 1H), 2.46 – 2.34 (m, 1H), 2.06 – 1.97 (m, 1H), 1.22 (t, *J* = 7.5 Hz, 3H). ^13^C NMR (126 MHz, DMSO-*d_6_*) δ 173.39, 171.61, 168.32, 161.21, 143.77, 142.82, 136.14, 127.50, 127.04, 124.76, 124.14, 123.75, 120.36, 119.91, 114.35, 112.65, 52.04, 47.67, 31.71, 23.02, 17.91, 16.18.

***Synthesis of 3-cyclopropyl-N-(2-(2,6-dioxopiperidin-3-yl)-1-oxoisoindolin-5-yl)-1H-indole-2-carboxamide 21.*** A solution of 3-(5-amino-1-oxoisoindolin-2-yl)piperidine-2,6-dione (30 mg, 0.116 mmol), 3-cyclopropyl-1*H*-indole-2-carboxylic acid **24g** (23.8 mg, 0.116 mmol), TCFH (35.7 mg, 0.127 mmol), and 1-methyl-1*H*-imidazole (28.5 mg, 0.347 mmol) was stirred in NMP (500 µL) for 16 h. The reaction mixture was purified using automated reversed-phase flash chromatography (Biotage Sfar C18 30 g column, acetonitrile/water gradient mobile phase with 0.1% formic acid additive). Product-containing fractions were lyophilized to obtain the desired product (5 mg, 9% yield). ^1^H NMR (500 MHz, DMSO*-d_6_*) δ 11.62 (s, 1H), 11.00 (s, 1H), 10.20 (s, 1H), 8.15 (s, 1H), 7.84 (d, *J* = 8.3 Hz, 1H), 7.74 (d, *J* = 8.3 Hz, 1H), 7.69 (d, *J* = 8.1 Hz, 1H), 7.43 (d, *J* = 8.2 Hz, 1H), 7.22 (t, *J* = 7.6 Hz, 1H), 7.06 (t, *J* = 7.6 Hz, 1H), 5.11 (dd, *J* = 13.3, 5.1 Hz, 1H), 4.49 (d, *J* = 17.2 Hz, 1H), 4.35 (d, *J* = 17.2 Hz, 1H), 2.92 (ddd, *J* = 18.4, 13.7, 5.6 Hz, 1H), 2.65 – 2.55 (m, 1H), 2.46 – 2.34 (m, 2H), 2.02 (dd, *J* = 12.6, 6.3 Hz, 1H), 1.06 (d, *J* = 9.3 Hz, 2H), 0.78 (d, *J* = 5.1 Hz, 2H). ^13^C NMR (126 MHz, DMSO*-d_6_*) δ 172.95, 171.15, 167.85, 160.59, 143.35, 142.14, 135.67, 129.34, 127.25, 126.70, 123.97, 123.75, 120.47, 119.53, 119.43, 113.83, 112.42, 51.59, 47.21, 31.25, 22.55, 6.78, 6.65.

***Synthesis of N-(2-(2,6-dioxopiperidin-3-yl)-1-oxoisoindolin-5-yl)-3-phenyl-1H-indole-2-carboxamide 22.*** To 3-phenyl-1*H*-indole-2-carboxylic acid **24h** (0.027 g, 0.12 mmol) in DMF (0.5 mL) at room temperature was added DIPEA (0.060 mL, 0.35 mmol), HATU (0.048 g, 0.13 mmol) and 3-(5-amino-1-oxoisoindolin-2-yl)piperidine-2,6-dione (0.030 g, 0.12 mmol). The reaction mixture was stirred at room temperature for 2 h then diluted with ethyl acetate (3 mL) and washed sequentially with brine (2 × 3 mL) and 5% LiCl (3 mL). The organic phase was dried over sodium sulfate, filtered, and concentrated *in vacuo*. The crude was purified using automated flash chromatography (Biotage Sfar Amino D 11g column, methanol/dichloromethane gradient mobile phase). Product-containing fractions were evaporated *in vacuo* to obtain a white solid (4.3 mg, 8% yield). ^1^H NMR (500 MHz, DMSO*-d_6_*) δ 12.00 (s, 1H), 10.98 (s, 1H), 10.10 (s, 1H), 7.97 (s, 1H), 7.68 (d, *J* = 8.3 Hz, 1H), 7.62 (d, *J* = 8.2 Hz, 1H), 7.57 – 7.50 (m, 4H), 7.46 (t, *J* = 7.7 Hz, 2H), 7.35 (t, *J* = 7.4 Hz, 1H), 7.31 (t, *J* = 7.6 Hz, 1H), 7.14 (t, *J* = 7.5 Hz, 1H), 5.09 (dd, *J* = 13.3, 5.1 Hz, 1H), 4.50 – 4.24 (m, 2H), 2.91 (ddd, *J* = 17.5, 13.7, 5.4 Hz, 1H), 2.67 – 2.56 (m, 1H), 2.38 (qd, *J* = 13.4, 4.4 Hz, 1H), 2.04 – 1.96 (m, 1H). ^13^C NMR (126 MHz, DMSO*-d_6_*) δ 173.38, 171.58, 168.22, 161.30, 143.75, 142.43, 136.13, 134.14, 130.30, 128.90, 128.67, 127.22, 126.84, 124.64, 124.16, 121.00, 120.49, 119.77, 118.94, 114.18, 112.90, 52.02, 47.64, 31.69, 22.99.

***Synthesis of 3-benzyl-N-(2-(2,6-dioxopiperidin-3-yl)-1-oxoisoindolin-5-yl)-1H-indole-2-carboxamide 23.*** A mixture of 3-(5-amino-1-oxoisoindolin-2-yl)piperidine-2,6-dione (0.021 g, 0.080 mmol), 3-benzyl-1*H*-indole-2-carboxylic acid **24i** (0.020 g, 0.080 mmol), TCFH (0.025 g, 0.088 mmol) and NMI (0.019 mL, 0.24 mmol) was stirred at room temperature in NMP (0.5 mL) for 2 h. The reaction mixture was diluted with ethyl acetate (3 mL) and washed with brine (3 × 3 mL). The organic phase was dried over sodium sulfate, filtered and concentrated *in vacuo*. The crude was purified using automated flash chromatography (Biotage Sfar Amino D 11g column, methanol/dichloromethane gradient mobile phase). Product-containing fractions were evaporated *in vacuo* to obtain a white solid (7.4 mg, 19% yield). ^1^H NMR (400 MHz, DMSO*-d_6_*) δ 11.62 (s, 1H), 10.99 (s, 1H), 10.41 (s, 1H), 8.15 (s, 1H), 7.80 – 7.68 (m, 2H), 7.63 (d, *J* = 8.1 Hz, 1H), 7.47 (d, *J* = 8.2 Hz, 1H), 7.31 (d, *J* = 7.2 Hz, 2H), 7.26-7.18 (m, 3H), 7.12-7.03 (m, 2H), 5.11 (dd, *J* = 13.3, 5.1 Hz, 1H), 4.54 – 4.29 (m, 4H), 3.00 – 2.85 (m, 1H), 2.69 – 2.57 (m, 1H), 2.46 – 2.31 (m, 1H), 2.04-1.99 (m, 1H). ^13^C NMR (101 MHz, DMSO*-d_6_*) δ 172.87, 171.09, 167.80, 160.77, 143.26, 142.22, 141.54, 135.67, 128.29, 128.12, 127.29, 127.24, 126.63, 125.59, 124.33, 123.64, 120.24, 119.63, 119.51, 113.95, 112.20, 51.55, 47.18, 31.20, 29.75, 22.52.

**Scheme S2. Preparation of Compound 6^a^**

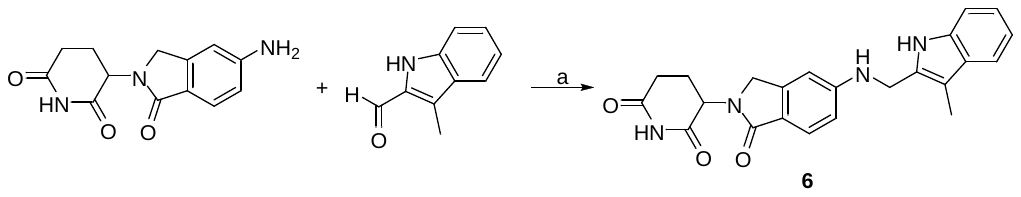

^a^Reagents and conditions: a) sodium triacetoxyborohydride (6.0 equiv), DMF/AcOH (3:1), rt, 16 h.

***Synthesis of 3-(5-(((3-methyl-1H-indol-2-yl)methyl)amino)-1-oxoisoindolin-2-yl)piperidine-2,6-dione*** ***6***. Sodium triacetoxyborohydride (147 mg, 0.694 mmol) was added to a stirring solution of 3-(5-amino-1-oxoisoindolin-2-yl)piperidine-2,6-dione (30 mg, 0.116 mmol) and 3-methyl-1*H*-indole-2-carbaldehyde (36.8 mg, 0.231 mmol) in anhydrous DMF (1 mL) and AcOH (0.3 mL), stirring at rt under N_2_. The reaction mixture was stirred for 16 h and then diluted with water (5 mL) and extracted with EtOAc (3 × 5 mL). The combined organic layers were washed with brine, dried over anhydrous Na_2_SO_4_, filtered, and concentrated. The crude product was purified using automated flash chromatography (Sfar Silica HC D 10 g column, methanol/dichloromethane gradient mobile phase). Product-containing fractions were concentrated to obtain the desired product (3 mg, 6% yield). HRMS (ESI) for C_23_H_23_N_4_O_3_ [M + H]^+^ calc’d 403.1770, obtained 403.1769. ^1^H NMR (500 MHz, DMSO-*d_6_*) δ 10.92 (s, 1H), 10.84 (s, 1H), 7.41 (dd, *J* = 14.6, 8.0 Hz, 2H), 7.28 (d, *J* = 8.0 Hz, 1H), 7.03 (t, *J* = 7.5 Hz, 1H), 6.95 (t, *J* = 7.4 Hz, 1H), 6.80 – 6.70 (m, 3H), 5.01 (dd, *J* = 13.3, 5.1 Hz, 1H), 4.41 (d, *J* = 5.1 Hz, 2H), 4.27 (d, *J* = 16.7 Hz, 1H), 4.14 (d, *J* = 16.7 Hz, 1H), 2.94 – 2.83 (m, 1H), 2.62 – 2.53 (m, 1H), 2.39 – 2.29 (m, 1H), 2.27 (s, 3H), 1.96 – 1.89 (m, 1H). ^13^C NMR (126 MHz, DMSO-*d_6_*) δ 172.96, 171.40, 168.67, 152.01, 144.35, 135.37, 132.03, 128.55, 123.85, 120.78, 119.29, 118.16, 117.90, 112.82, 110.87, 106.63, 104.56, 51.28, 46.78, 38.74, 31.28, 22.63, 8.37.

**Scheme S3. Preparation of Compound 8^a^**

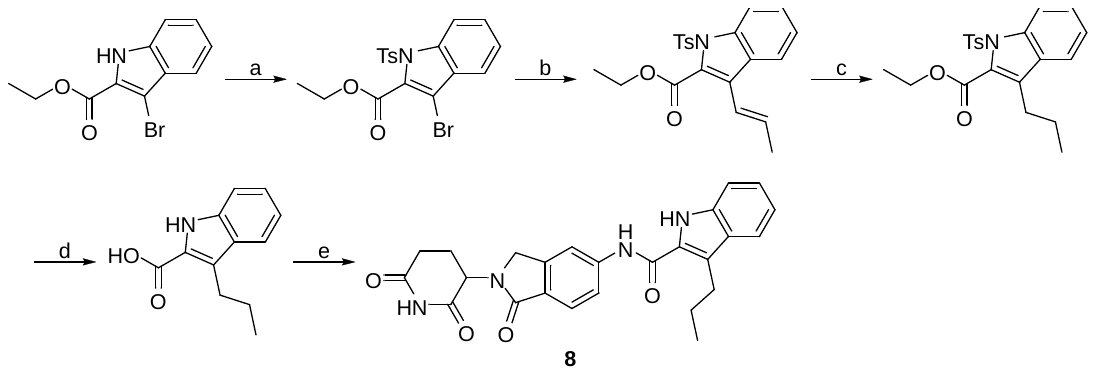

^a^Reagents and conditions: a) 4-methylbenzenesulfonyl chloride, NaH, DMF, 0 °C to rt, 3 h; b) *trans*-1-propenylboronic acid pinacol ester, K_2_CO_3_, Pd(PPh_3_)_4_, Dioxane:H_2_O (3:1), 90 °C, 16 h; c) Pd/C, triethylsilane, EtOAc, rt, 22 h; d) 2M KOH, EtOH, 80 °C, 1 h; e) 3-(5-amino-1-oxoisoindolin-2-yl)piperidine-2,6-dione (1.0 equiv), TCFH (2.0 equiv), *N*-methylimidazole (3.0 equiv), NMP, rt, 3 h.

***(a)*** ***Synthesis of ethyl 3-bromo-1-tosyl-1H-indole-2-carboxylate*.** To a solution of NaH (97 mg, 2.424 mmol) in anhydrous DMF (2 mL) stirring at 0 °C under N_2_ was added a solution of ethyl 3-bromo-1*H*-indole-2-carboxylate (500 mg, 1.865 mmol) in DMF (2 mL) dropwise. The reaction mixture was stirred for 30 min and then 4-methylbenzenesulfonyl chloride (356 mg, 1.865 mmol) dissolved in DMF (1 mL) was added dropwise. The reaction mixture was stirred for 1 h at 0 °C followed by 1 h at rt. The solution was then diluted with water (25 mL) and washed with EtOAc (2 × 25 mL). The combined organic layers were washed with brine, dried with anhydrous Na_2_SO_4_, filtered, and concentrated. The crude mixture was purified by flash column chromatography (Biotage Isolera, 28 g KP-amino Sfar column, 0–25% hexanes:EtOAc) to give the desired product (509 mg, 65% yield). ^1^H NMR (500 MHz, CDCl_3_) δ 8.02 (d, *J* = 8.4 Hz, 1H), 7.84 (d, *J* = 8.1 Hz, 2H), 7.51 (dd, *J* = 7.9, 1.1 Hz, 1H), 7.44 (ddt, *J* = 8.4, 7.4, 1.1 Hz, 1H), 7.33 (t, *J* = 7.6 Hz, 1H), 7.23 (d, *J* = 8.1 Hz, 2H), 4.53 (qd, *J* = 7.1, 0.9 Hz, 2H), 2.34 (s, 3H), 1.47 (td, *J* = 7.2, 0.9 Hz, 3H).

**(b)** ***Synthesis* *of ethyl (E)-3-(prop-1-en-1-yl)-1-tosyl-1H-indole-2-carboxylate.*** A mixture of ethyl 3-bromo-1-tosyl-1*H*-indole-2-carboxylate (0.10 g, 0.24 mmol), *trans*-1-propenylboronic acid pinacol ester (0.047 mL, 0.25 mmol), potassium carbonate (0.065 g, 0.47 mmol), and Pd(PPh_3_)_4_ (0.027 g, 0.024 mmol) in 3:1 dioxane/water (2 mL) was degassed then stirred at 90 °C under nitrogen atmosphere for 1 h, then cooled to room temperature. The reaction mixture was diluted with ethyl acetate (10 mL) and washed with brine (3 × 10 mL). The organic phase was dried over sodium sulfate, filtered, and concentrated *in vacuo*. The crude was purified using automated flash chromatography (Biotage Sfar Silica HC D 25g column, ethyl acetate/hexanes gradient mobile phase). Product-containing fractions were evaporated *in vacuo* to obtain a colorless oil (0.038 g, 42%). LC-MS (ESI) *m/z*: 384.18 [M+H]^+^. ^1^H NMR (400 MHz, CDCl_3_) δ 8.08 – 8.00 (m, 1H), 7.80 – 7.69 (m, 3H), 7.39 (ddd, *J* = 8.4, 7.3, 1.2 Hz, 1H), 7.30 – 7.25 (m, 1H), 7.18 (d, *J* = 8.0 Hz, 2H), 6.58 (dq, *J* = 16.1, 1.5 Hz, 1H), 6.43 (dq, *J* = 16.1, 6.5 Hz, 1H), 4.51 (q, *J* = 7.2 Hz, 2H), 2.33 (s, 3H), 1.95 (dd, *J* = 6.5, 1.6 Hz, 3H), 1.46 (t, *J* = 7.2 Hz, 3H). ^13^C NMR (101 MHz, CDCl_3_) δ 162.82, 144.91, 136.96, 134.28, 131.88, 129.52, 128.47, 127.69, 127.23, 126.58, 124.95, 124.32, 121.64, 120.64, 115.49, 62.30, 21.58, 19.40, 14.09.

**(c)** ***Synthesis* *of ethyl 3-propyl-1-tosyl-1H-indole-2-carboxylate.*** To a microwave vial fitted with a magnetic stir bar was added 10% Pd/C (0.011 g, 9.91 µmol). The vial was capped and purged with nitrogen. Ethyl (*E*)-3-(prop-1-en-1-yl)-1-tosyl-1*H*-indole-2-carboxylate (0.038 g, 0.099 mmol) was diluted in ethyl acetate (1 mL) and added to the microwave vial via syringe. The microwave vial was then degassed and purged with nitrogen. Triethylsilane (0.047 mL, 0.73 mmol) was added to the microwave vial dropwise at rt. After stirring at rt for 22 h the reaction mixture was filtered through celite and the filtrate was concentrated *in vacuo*. The crude was purified using automated flash chromatography (Biotage Sfar HC D 10g column, ethyl acetate/hexanes gradient mobile phase). Product-containing fractions were evaporated *in vacuo* to obtain a colorless oil (0.022 g, 58%). LC-MS (ESI) *m/z*: 386.29 [M+H]^+^. ^1^H NMR (400 MHz, CDCl_3_) δ 8.09 – 8.01 (m, 1H), 7.79 – 7.70 (m, 2H), 7.48 (dt, *J* = 7.9, 0.9 Hz, 1H), 7.39 (ddd, *J* = 8.4, 7.3, 1.2 Hz, 1H), 7.28 – 7.23 (m, 1H), 7.18 (d, *J* = 8.0 Hz, 2H), 4.48 (q, *J* = 7.2 Hz, 2H), 2.81 – 2.68 (m, 2H), 2.33 (s, 3H), 1.68 – 1.61 (m, 2H), 1.45 (t, *J* = 7.2 Hz, 3H), 0.90 (t, *J* = 7.4 Hz, 3H). ^13^C NMR (101 MHz, CDCl_3_) δ 162.62, 144.70, 137.26, 134.32, 130.27, 129.93, 129.34, 128.51, 127.22, 126.65, 124.02, 120.58, 115.78, 61.97, 26.28, 23.04, 21.57, 14.14, 13.83.

**(d)** ***Synthesis* *of*** ***3-propyl-1H-indole-2-carboxylic acid.*** To a solution of ethyl 3-propyl-1-tosyl-1*H*-indole-2-carboxylate (0.022 g, 0.057 mmol) in ethanol (1.0 mL) was added 2M KOH (0.23 mL, 0.46 mmol). The reaction mixture was stirred at 80 °C for 1 h, then cooled to rt. The crude was concentrated *in vacuo* to remove volatiles, then was purified using automated reversed-phase flash chromatography (Biotage Sfar C18 12g column, acetonitrile/water gradient mobile phase with 0.1% formic acid additive). Product-containing fractions were lyophilized to obtain a white solid (8 mg, 69%). LC-MS (ESI) *m/z*: 204.20 [M+H]^+^. ^1^H NMR (500 MHz, DMSO-*d_6_*) δ 11.29 (s, 1H), 7.62 (d, *J* = 8.1 Hz, 1H), 7.37 (d, *J* = 8.3 Hz, 1H), 7.27 – 7.12 (m, 1H), 7.07 – 6.91 (m, 1H), 3.05 – 2.98 (m, 2H), 1.61 (h, *J* = 7.4 Hz, 2H), 0.89 (t, *J* = 7.4 Hz, 3H). ^13^C NMR (126 MHz, DMSO-*d_6_*) δ 164.06, 136.40, 127.96, 124.91, 124.65, 122.73, 120.68, 119.49, 112.73, 26.42, 24.39, 14.45.

***(e) Synthesis of N-(2-(2,6-dioxopiperidin-3-yl)-1-oxoisoindolin-5-yl)-3-propyl-1H-indole-2-carboxamide*** ***8***. A mixture of 3-(5-amino-1-oxoisoindolin-2-yl)piperidine-2,6-dione (0.009 g, 0.034 mmol), 3-propyl-1*H*-indole-2-carboxylic acid (0.007 g, 0.034 mmol), TCFH (0.011 g, 0.038 mmol) and *N*-methylimidazole (0.008 mL, 0.10 mmol) was stirred at rt in NMP (0.5 mL) for 3 h. The reaction mixture was diluted with ethyl acetate (3 mL) and washed with brine (3 × 3 mL). The organic phase was dried over sodium sulfate, filtered and concentrated *in vacuo*. The crude was purified using automated flash chromatography (Biotage Sfar Amino D 11g column, methanol/dichloromethane gradient mobile phase). Product-containing fractions were evaporated *in vacuo* to obtain a white solid (3.3 mg, 22%). HRMS (ESI) for C_25_H_25_N_4_O_4_ [M+H]^+^ calc’d 445.1876, obtained 445.1871. ^1^H NMR (600 MHz, DMSO-*d_6_*) δ 11.45 (s, 1H), 10.99 (s, 1H), 10.29 (s, 1H), 8.15 (s, 1H), 7.78 – 7.69 (m, 2H), 7.67 (d, *J* = 8.1 Hz, 1H), 7.46 (d, *J* = 8.2 Hz, 1H), 7.30 – 7.21 (m, 1H), 7.12 – 7.03 (m, 1H), 5.11 (dd, *J* = 13.3, 5.1 Hz, 1H), 4.48 (d, *J* = 17.2 Hz, 1H), 4.34 (d, *J* = 17.2 Hz, 1H), 3.10 – 3.00 (m, 2H), 2.92 (ddd, *J* = 17.4, 13.7, 5.4 Hz, 1H), 2.68 – 2.55 (m, 1H), 2.39 (td, *J* = 12.9, 4.5 Hz, 1H), 2.06 – 1.98 (m, 1H), 1.64 (q, *J* = 7.4 Hz, 2H), 0.90 (t, *J* = 7.3 Hz, 3H). ^13^C NMR (151 MHz, DMSO-*d_6_*) δ 173.38, 171.60, 168.32, 161.28, 143.78, 142.84, 136.09, 128.02, 127.53, 127.02, 124.68, 124.13, 121.93, 120.49, 119.90, 119.89, 114.33, 112.61, 52.05, 47.68, 31.71, 26.50, 24.34, 23.03, 14.49.

**Scheme S4. Preparation of Compound 9^a^**

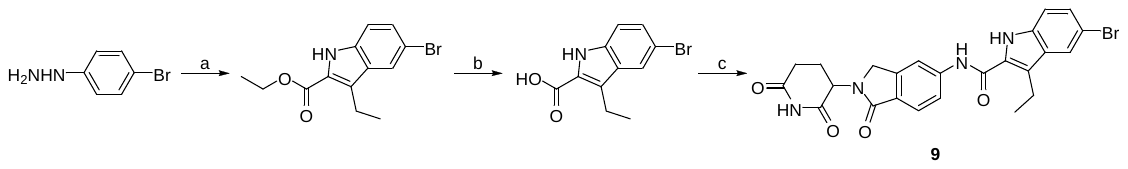

^a^Reagents and conditions: a) ethyl 2-oxopentanoate, *p*-toluenesulfonic acid monohydrate, EtOH, 90 °C; b) 2M KOH, EtOH, 90 °C; c) 3-(5-amino-1-oxoisoindolin-2-yl)piperidine-2,6-dione (1.0 equiv), TCFH (2.0 equiv), *N*-methylimidazole (3.0 equiv), NMP, rt, 16 h.

***(a) Synthesis of ethyl 5-bromo-3-ethyl-1H-indole-2-carboxylate*.** A mixture of 4-bromophenylhydrazine hydrochloride (0.56 g, 2.5 mmol), ethyl-2-oxo-4-valerate (0.36 mL, 2.5 mmol), and *p*-TsOH monohydrate (0.76 g, 4.0 mmol) were stirred in ethanol (20 mL) at 90 °C. After 3 h the reaction mixture was cooled to rt and an additional 3 eq. of *p*-TsOH-H_2_O was added (1.43 g, 7.5 mmol). The reaction mixture was stirred at 90 °C for a total of 18 h, then was cooled to room temperature. The crude reaction mixture was diluted with ethyl acetate (50 mL) and washed with saturated sodium bicarbonate solution (3 × 50 mL). The organic phase was dried over sodium sulfate, filtered, and concentrated *in vacuo*. The crude was purified using automated flash chromatography (Biotage Sfar Silica HC D 50g column, ethyl acetate/hexanes gradient mobile phase). Product-containing fractions were evaporated *in vacuo* to obtain a pale orange solid (0.474 g, 64%). LC-MS (ESI) *m/z*: 296.17 [M+H]^+^. ^1^H NMR (500 MHz, CDCl_3_) δ 8.69 (s, 1H), 7.82 (s, 1H), 7.38 (d, *J* = 8.7 Hz, 1H), 7.24 (s, 1H), 4.42 (q, *J* = 7.1 Hz, 2H), 3.07 (q, *J* = 7.5 Hz, 2H), 1.43 (t, *J* = 7.1 Hz, 3H), 1.26 (t, *J* = 7.5 Hz, 3H). ^13^C NMR (126 MHz, CDCl_3_) δ 162.06, 134.34, 129.35, 128.42, 126.08, 123.83, 123.31, 113.22, 113.14, 60.93, 18.00, 15.39, 14.40.

***(b) Synthesis of 5-bromo-3-ethyl-1H-indole-2-carboxylic acid*.** To a solution of ethyl 5-bromo-3-ethyl-1*H*-indole-2-carboxylate (0.030 g, 0.10 mmol) in EtOH (2 mL) was added 2M KOH (0.41 mL, 0.81 mmol). The reaction mixture was stirred at 80 °C for 1 h, then cooled to rt and concentrated *in vacuo* to remove volatiles. The crude was diluted with water (3 mL) and cooled on an ice bath. A solution of 1 M aqueous HCl was used to adjust the crude to pH 3 and form a white precipitate which was then extracted into ethyl acetate (2 × 5 mL). The combined organics were washed with brine (5 mL), dried over sodium sulfate, filtered, and concentrated *in vacuo* to obtain a white solid (0.027 g, 99%). LC-MS (ESI) *m/z*: 266.00 [M-H]^-^. ^1^H NMR (500 MHz, DMSO-*d_6_*) δ 13.07 (s, 1H), 11.59 (s, 1H), 7.86 (s, 1H), 7.35 (s, 2H), 3.02 (q, *J* = 7.4 Hz, 2H), 1.17 (t, *J* = 7.4 Hz, 3H). ^13^C NMR (126 MHz, DMSO-*d_6_*) δ 163.52, 135.10, 129.04, 127.52, 125.19, 124.48, 122.84, 114.92, 112.19, 17.69, 16.19.

***(c) Synthesis of* *5-bromo-N-(2-(2,6-dioxopiperidin-3-yl)-1-oxoisoindolin-5-yl)-3-ethyl-1H-indole-2-carboxamide*** ***9****.* A mixture of 3-(5-amino-1-oxoisoindolin-2-yl)piperidine-2,6-dione (0.024 g, 0.093 mmol), 5-bromo-3-ethyl-1*H*-indole-2-carboxylic acid (0.025 g, 0.093 mmol), TCFH (0.029 g, 0.10 mmol) and *N*-methylimidazole (0.022 mL, 0.28 mmol) was stirred at room temperature in NMP (0.5 mL) for 3 h. The reaction mixture was diluted with ethyl acetate (3 mL) and washed sequentially with saturated sodium bicarbonate solution (3 × 3 mL) followed by brine (3 × 3 mL). The organic phase was separated, concentrated *in vacuo*, then purified using automated flash chromatography (Sfar Amino D 11g column, MeOH/dichloromethane gradient mobile phase). Product-containing fractions were evaporated *in vacuo* to obtain a white solid (4.9 mg, 10%). HRMS (ESI) for C_24_H_22_BrN_4_O4 [M+H]^+^ calc’d 509.0824, obtained 509.0839. ^1^H NMR (500 MHz, DMSO-*d_6_*) δ 11.71 (s, 1H), 10.98 (s, 1H), 10.41 (s, 1H), 8.14 (s, 1H), 7.88 (d, *J* = 1.6 Hz, 1H), 7.78 – 7.70 (m, 2H), 7.43 (d, *J* = 8.7 Hz, 1H), 7.36 (dd, *J* = 8.7, 1.8 Hz, 1H), 5.11 (dd, *J* = 13.3, 5.1 Hz, 1H), 4.54 – 4.22 (m, 2H), 3.04 (q, *J* = 7.4 Hz, 2H), 2.98 – 2.86 (m, 1H), 2.66 – 2.55 (m, 1H), 2.46 – 2.33 (m, 1H), 2.04-1.98 (m, 1H), 1.19 (t, *J* = 7.5 Hz, 3H). ^13^C NMR (126 MHz, DMSO-*d_6_*) δ 173.40, 171.61, 168.30, 160.95, 143.78, 142.67, 134.69, 129.23, 128.62, 127.19, 124.16, 122.84, 122.60, 119.98, 114.79, 114.44, 112.45, 52.05, 47.68, 31.71, 23.02, 17.71, 16.18.

**Scheme S5. Preparation of Compound 10^a^**

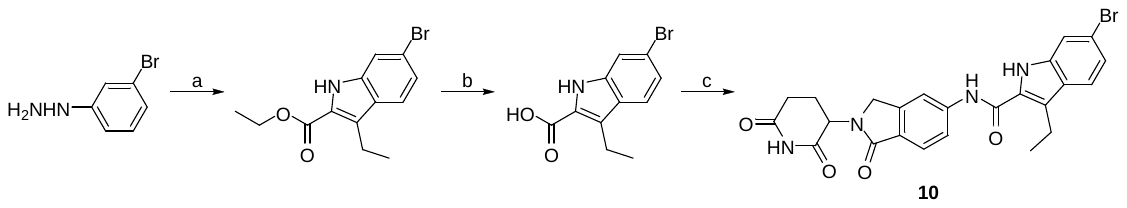

^a^Reagents and conditions: a) ethyl 2-oxopentanoate, *p*-toluenesulfonic acid monohydrate, EtOH, 90 °C; b) 2M KOH, EtOH, 90 °C; c) 3-(5-amino-1-oxoisoindolin-2-yl)piperidine-2,6-dione (1.0 equiv), TCFH (2.0 equiv), *N*-methylimidazole (3.0 equiv), NMP, rt, 3 h.

***(a) Synthesis of ethyl ethyl 6-bromo-3-ethyl-1H-indole-2-carboxylate*.** A mixture of 3-bromophenylhydrazine hydrochloride (0.56 g, 0.22 mmol), ethyl-2-oxo-4-valerate (0.36 mL, 2.50 mmol), and *p*-toluenesulfonic acid monohydrate (0.76 g, 4.00 mmol) were stirred in ethanol (20 mL) at 90 °C. After 3 h the reaction mixture was cooled to rt and an additional 3 eq. of *p*-toluenesulfonic acid monohydrate was added (1.4 g, 7.5 mmol). The reaction mixture was then stirred at 90 °C for a total of 22 h and cooled to rt. The crude was diluted with ethyl acetate (50 mL) and washed with saturated sodium bicarbonate solution (3 × 50 mL). The organic phase was dried over sodium sulfate, filtered, and concentrated. The crude was purified using automated flash chromatography (Biotage Sfar Silica HC D 25 g column, ethyl acetate/hexanes gradient mobile phase). Product-containing fractions were evaporated *in vacuo* to obtain a white solid (0.113 g, 15%). LC-MS (ESI) *m/z*: 296.10 [M+H]^+^. ^1^H NMR (500 MHz, CDCl_3_) δ 8.66 (s, 1H), 7.57 – 7.51 (m, 2H), 7.24 (dd, *J* = 8.5, 1.5 Hz, 1H), 4.42 (q, *J* = 7.1 Hz, 2H), 3.09 (q, *J* = 7.5 Hz, 2H), 1.43 (t, *J* = 7.1 Hz, 3H), 1.26 (t, *J* = 7.5 Hz, 3H). ^13^C NMR (126 MHz, CDCl_3_) δ 162.13, 136.44, 126.87, 126.56, 123.50, 123.28, 122.07, 119.32, 114.56, 60.90, 18.01, 15.42, 14.41.

***(b) Synthesis of 6-bromo-3-ethyl-1H-indole-2-carboxylic acid*.** To a solution of ethyl 6-bromo-3-ethyl-1*H*-indole-2-carboxylate (0.040 g, 0.14 mmol) in EtOH (2 mL) was added 2M KOH (0.54 mL, 1.1 mmol). The reaction mixture was stirred at 80 °C for 1 h, then cooled to rt and concentrated *in vacuo* to remove volatiles. The crude was diluted with water (3 mL), cooled on an ice bath, then acidified to pH 3 using 1M aqueous HCl solution to form a white precipitate which was extracted into ethyl acetate (2 × 5 mL). The combined organics were washed with brine (5 mL), dried over sodium sulfate, filtered, and concentrated *in vacuo* to obtain a white solid (0.032 g, 88%). LC-MS (ESI) *m/z*: 266.00 [M-H]^-^. ^1^H NMR (500 MHz, DMSO-*d_6_*) δ 13.03 (s, 1H), 11.51 (s, 1H), 7.63 (d, *J* = 8.6 Hz, 1H), 7.54 (s, 1H), 7.17 (dd, *J* = 8.6, 1.5 Hz, 1H), 3.03 (q, *J* = 7.5 Hz, 2H), 1.17 (t, *J* = 7.5 Hz, 3H). ^13^C NMR (126 MHz, DMSO-*d_6_*) δ 163.54, 137.21, 126.32, 125.21, 124.65, 122.68, 122.64, 117.83, 115.19, 17.74, 16.18.

***(c) Synthesis of* *6-bromo-N-(2-(2,6-dioxopiperidin-3-yl)-1-oxoisoindolin-5-yl)-3-ethyl-1H-indole-2-carboxamide*** ***10****.* A mixture of 3-(5-amino-1-oxoisoindolin-2-yl)piperidine-2,6-dione (0.029 g, 0.11 mmol), 6-bromo-3-ethyl-1*H*-indole-2-carboxylic acid (0.030 g, 0.11 mmol), TCFH (0.035 g, 0.12 mmol) and *N*-methylimidazole (0.027 mL, 0.34 mmol) was stirred at rt in NMP (0.5 mL) for 4 h. The reaction mixture was then diluted with ethyl acetate (3 mL) and washed sequentially with saturated sodium bicarbonate solution (3 × 3 mL) followed by brine (3 × 3 mL). The organic phase was separated, concentrated, and purified using automated flash chromatography (Biotage Sfar Amino D 11g column, methanol/dichloromethane gradient mobile phase). Product-containing fractions were evaporated *in vacuo* to obtain a white solid (7.4 mg, 13%). HRMS (ESI) for C_24_H_22_BrN_4_O_4_ [M+H]^+^ calc’d 509.0824, obtained 509.0836. ^1^H NMR (500 MHz, DMSO-*d_6_*) δ 11.62 (s, 1H), 10.99 (s, 1H), 10.39 (s, 1H), 8.13 (s, 1H), 7.73 (t, *J* = 6.6 Hz, 2H), 7.69 – 7.61 (m, 2H), 7.21 (d, *J* = 8.6 Hz, 1H), 5.11 (dd, *J* = 13.2, 4.9 Hz, 1H), 4.52 – 4.26 (m, 2H), 3.06 (q, *J* = 7.2 Hz, 2H), 2.92 (ddd, *J* = 17.7, 13.4, 5.4 Hz, 1H), 2.61 (d, *J* = 17.8 Hz, 1H), 2.44 – 2.34 (m, 1H), 2.04-1.99 (m, 1H), 1.20 (t, *J* = 7.4 Hz, 3H). ^13^C NMR (126 MHz, DMSO-*d_6_*) δ 173.39, 171.60, 168.29, 160.96, 143.77, 142.65, 136.82, 128.04, 127.18, 126.49, 124.16, 123.52, 122.95, 122.30, 119.99, 117.45, 115.06, 114.46, 52.04, 47.67, 31.70, 23.01, 17.80, 16.13.

**Scheme S6. Preparation of Compound 11^a^**

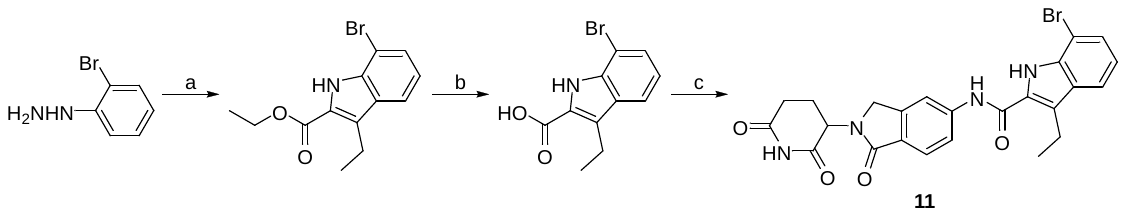

^a^Reagents and conditions: a) ethyl 2-oxopentanoate, *p*-toluenesulfonic acid monohydrate, EtOH, 90 °C; b) 2M KOH, EtOH, 90 °C; c) 3-(5-amino-1-oxoisoindolin-2-yl)piperidine-2,6-dione (1.0 equiv), TCFH (2.0 equiv), *N*-methylimidazole (3.0 equiv), NMP, rt, 4 h.

***(a)*** ***Synthesis of ethyl 7-bromo-3-ethyl-1H-indole-2-carboxylate*.** To a solution of (2-bromophenyl)hydrazine hydrochloride (244 mg, 1.092 mmol), and ethyl 2-oxopentanoate (157 mg, 1.092 mmol), in EtOH (10 mL), was added *p*-toluenesulfonic acid monohydrate (623 mg, 3.28 mmol) and stirred at 90 °C. After 3 h, additional *p*-toluenesulfonic acid monohydrate (623 mg, 3.28 mmol) was added and the reaction mixture stirred for an additional 16 h. The solution was cooled to rt, diluted with saturated NaHCO_3_ (50 mL), and washed with EtOAc (50 mL). The organic layer was washed with brine, dried with anhydrous Na_2_SO_4_, filtered, and concentrated. The crude mixture was purified by flash column chromatography (Biotage Isolera, 25 g Sfar column, 0–50% hexanes:EtOAc) to give the desired product (101 mg, 31% yield). ^1^H NMR (500 MHz, CDCl_3_) δ 8.74 (s, 1H), 7.64 (d, *J* = 8.1 Hz, 1H), 7.47 (d, *J* = 7.5 Hz, 1H), 7.02 (t, *J* = 7.8 Hz, 1H), 4.45 (q, *J* = 7.1 Hz, 2H), 3.10 (q, *J* = 7.5 Hz, 2H), 1.44 (t, *J* = 7.2 Hz, 3H), 1.27 (t, *J* = 7.5 Hz, 3H).

***(b) Synthesis of 7-bromo-3-ethyl-1H-indole-2-carboxylic acid*.** A solution of ethyl 7-bromo-3-ethyl-1*H*-indole-2-carboxylate (60 mg, 0.203 mmol) in 2M KOH (0.8 mL) and EtOH (2 mL) was stirred at 90 °C. After 1 h, the reaction mixture was cooled to rt and then directly purified using automated reversed-phase flash chromatography (Biotage Sfar C18 30 g column, acetonitrile/water gradient mobile phase with 0.1% formic acid additive). Product-containing fractions were lyophilized to obtain a white solid (50 mg, 92%). ^1^H NMR (400 MHz, DMSO-*d_6_*) δ 13.18 (s, 1H), 11.03 (s, 1H), 7.69 (d, *J* = 8.0 Hz, 1H), 7.48 (d, *J* = 7.5 Hz, 1H), 7.01 (t, *J* = 7.7 Hz, 1H), 3.05 (q, *J* = 7.4 Hz, 2H), 1.18 (t, *J* = 7.4 Hz, 3H).

***(c) Synthesis of* *7-bromo-N-(2-(2,6-dioxopiperidin-3-yl)-1-oxoisoindolin-5-yl)-3-ethyl-1H-indole-2-carboxamide*** ***11****.* A solution of 3-(5-amino-1-oxoisoindolin-2-yl)piperidine-2,6-dione (30 mg, 0.116 mmol), 7-bromo-3-ethyl-1*H*-indole-2-carboxylic acid (31.0 mg, 0.116 mmol), TCFH (64.9 mg, 0.231 mmol), and *N*-methylimidazole (27.7 µL, 0.347 mmol) was stirred for 90 min in NMP (500 µL). The reaction mixture was purified using automated reversed-phase flash chromatography (Biotage Sfar C18 30 g column, acetonitrile/water gradient mobile phase with 0.1% formic acid additive). Product-containing fractions were lyophilized to obtain a white solid (18 mg, 31% yield). HRMS (ESI) for C_24_H_22_BrN_4_O_4_ [M+H]^+^ calc’d 509.0824, obtained 509.0818. ^1^H NMR (500 MHz, DMSO-*d_6_*) δ 11.46 (s, 1H), 11.00 (s, 1H), 10.54 (s, 1H), 8.18 (s, 1H), 7.79 (dd, *J* = 8.3, 1.7 Hz, 1H), 7.77 – 7.70 (m, 2H), 7.52 (d, *J* = 7.5 Hz, 1H), 7.06 (t, *J* = 7.7 Hz, 1H), 5.11 (dd, *J* = 13.3, 5.1 Hz, 1H), 4.49 (d, *J* = 17.2 Hz, 1H), 4.35 (d, *J* = 17.2 Hz, 1H), 3.10 (q, *J* = 7.4 Hz, 2H), 2.92 (ddd, *J* = 18.0, 13.6, 5.4 Hz, 1H), 2.67 – 2.57 (m, 1H), 2.40 (qd, *J* = 13.2, 4.4 Hz, 1H), 2.02 (tt, *J* = 6.6, 4.2 Hz, 1H), 1.21 (t, *J* = 7.5 Hz, 3H). ^13^C NMR (126 MHz, DMSO-*d_6_*) δ 172.93, 171.15, 167.84, 159.82, 143.28, 142.11, 134.20, 128.62, 127.31, 127.01, 126.77, 125.60, 123.63, 120.95, 119.75, 119.43, 114.24, 104.64, 51.59, 47.21, 31.24, 22.56, 17.69, 15.48.

**Scheme S7. Preparation of Compound 12^a^**

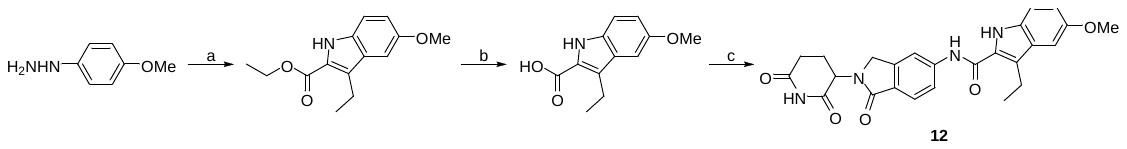

^a^Reagents and conditions: a) ethyl 2-oxopentanoate, *p*-toluenesulfonic acid monohydrate, EtOH, 90 °C; b) 2M KOH, EtOH, 90 °C; c) 3-(5-amino-1-oxoisoindolin-2-yl)piperidine-2,6-dione (1.0 equiv), TCFH (2.0 equiv), *N*-methylimidazole (3.0 equiv), NMP, rt, 90 min.

***(a) Synthesis of ethyl 3-ethyl-5-methoxy-1H-indole-2-carboxylate*.** To a stirring solution of (4-methoxyphenyl)hydrazine hydrochloride (250 mg, 1.432 mmol) and ethyl 2-oxopentanoate (206 mg, 1.432 mmol) in EtOH (8 mL) was added *p*-toluenesulfonic acid monohydrate (817 mg, 4.29 mmol) and heated to 90 °C. After 3 h, the reaction mixture was allowed to cool to rt, diluted with EtOAc (30 mL) and then washed with saturated NaHCO_3_ (30 mL). The organic layer was washed with brine, dried with anhydrous Na_2_SO_4_, filtered, and concentrated. The crude mixture was purified by flash column chromatography (Biotage Isolera, 25 g Sfar column, 0–30% hexanes:EtOAc) to give the desired product (200 mg, 57% yield). ^1^H NMR (500 MHz, CDCl_3_) δ 8.61 (s, 1H), 7.27 (d, *J* = 9.6 Hz, 1H), 7.05 (d, *J* = 2.4 Hz, 1H), 7.03 – 6.97 (m, 1H), 4.41 (q, *J* = 7.1 Hz, 2H), 3.88 (s, 3H), 3.09 (q, *J* = 7.5 Hz, 2H), 1.42 (t, *J* = 7.1 Hz, 3H), 1.28 (t, *J* = 7.5 Hz, 3H).

***(b)*** ***Synthesis of 3-ethyl-5-methoxy-1H-indole-2-carboxylic acid*.** A solution of ethyl 3-ethyl-5-methoxy-1*H*-indole-2-carboxylate (175 mg, 0.708 mmol) in 2M KOH (2.8 mL) and EtOH (3 mL) was stirred at 90 °C. After 4 h, the reaction mixture was cooled to rt and then directly purified using automated reversed-phase flash chromatography (Biotage Sfar C18 30 g column, acetonitrile/water gradient mobile phase with 0.1% formic acid additive). Product-containing fractions were lyophilized to obtain a white solid (199 mg). ^1^H NMR (500 MHz, DMSO-*d_6_*) δ 10.49 (s, 1H), 7.19 (d, *J* = 8.7 Hz, 1H), 6.93 (d, *J* = 2.4 Hz, 1H), 6.70 (dd, *J* = 8.8, 2.4 Hz, 1H), 3.74 (s, 3H), 3.05 (q, *J* = 7.4 Hz, 2H), 1.14 (t, *J* = 7.4 Hz, 3H). ^13^C NMR (126 MHz, DMSO-*d_6_*) δ 165.63, 165.37, 152.64, 129.81, 127.89, 118.16, 112.48, 100.26, 55.28, 17.26, 15.91.

***(c) Synthesis of* *N-(2-(2,6-dioxopiperidin-3-yl)-1-oxoisoindolin-5-yl)-3-ethyl-5-methoxy-1H-indole-2-carboxamide*** ***12****.* A solution of 3-(5-amino-1-oxoisoindolin-2-yl)piperidine-2,6-dione (40 mg, 0.154 mmol), 3-ethyl-5-methoxy-1*H*-indole-2-carboxylic acid (42 mg, 0.192 mmol), TCFH (87 mg, 0.309 mmol), and *N*-methylimidazole (36.9 µL, 0.463 mmol) was stirred for 16 h in NMP (500 µL). The reaction mixture was diluted with water (5 mL) and washed with EtOAc (3 × 5 mL). The combined organic layers were washed with brine, dried with anhydrous Na_2_SO_4_, filtered, and concentrated. The crude mixture was purified by flash column chromatography (Biotage Isolera, 11 g KP-amino Sfar column, 0–8% DCM:MeOH) to give the desired product (4 mg, 6% yield). HRMS (ESI) for C_25_H_25_N_4_O_5_ [M+H]^+^ calc’d 461.1825, obtained 461.1823. ^1^H NMR (500 MHz, DMSO-*d_6_*) δ 11.30 (s, 1H), 10.99 (s, 1H), 10.24 (s, 1H), 8.15 (s, 1H), 7.72 (dq, *J* = 17.5, 8.3 Hz, 2H), 7.36 (d, *J* = 8.8 Hz, 1H), 7.10 (d, *J* = 2.4 Hz, 1H), 6.92 (dd, *J* = 8.9, 2.4 Hz, 1H), 5.11 (dd, *J* = 13.2, 5.1 Hz, 1H), 4.47 (d, *J* = 17.2 Hz, 1H), 4.34 (d, *J* = 17.3 Hz, 1H), 3.80 (s, 3H), 3.06 (q, *J* = 7.4 Hz, 2H), 2.92 (ddd, *J* = 18.4, 13.5, 5.6 Hz, 1H), 2.61 (d, *J* = 18.9 Hz, 1H), 2.44 – 2.34 (m, 1H), 2.06 – 1.96 (m, 1H), 1.21 (t, *J* = 7.4 Hz, 3H). ^13^C NMR (126 MHz, DMSO-*d_6_*) δ 172.92, 171.14, 167.86, 160.67, 153.57, 143.29, 142.41, 130.90, 127.31, 127.02, 126.48, 123.65, 123.06, 119.40, 115.48, 113.82, 113.11, 100.30, 55.32, 51.56, 47.19, 31.23, 22.55, 17.43, 15.57.

**Scheme S8. Preparation of Compound 13^a^**

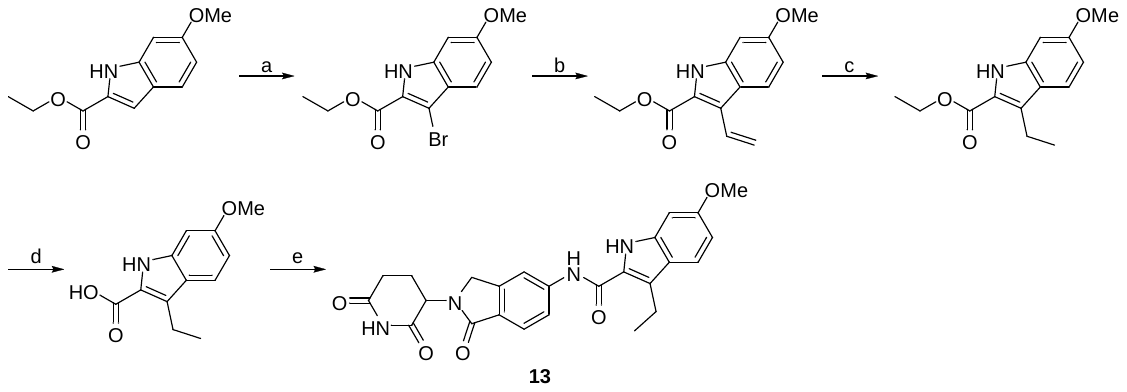

^a^Reagents and conditions: a) *N*-bromosuccinimide, DMF, rt, 16 h; b) 4,4,6-trimethyl-2-vinyl-1,3,2-dioxaborinane, K_2_CO_3_, Pd(PPh_3_)_4_, Dioxane:H_2_O (3:1), 90 °C, 16 h; c) Pd/C, triethylsilane, MeOH, rt, 1 h; d) 2M KOH, H_2_O, 90 °C, 16 h; e) 3-(5-amino-1-oxoisoindolin-2-yl)piperidine-2,6-dione (1.0 equiv), TCFH (2.0 equiv), *N*-methylimidazole (3.0 equiv), NMP, rt, 16 h.

***(a)*** ***Synthesis of ethyl 3-bromo-6-methoxy-1H-indole-2-carboxylate*.** *N-*Bromosuccinimide (170 mg, 0.958 mmol) was added portionwise to a solution of ethyl 6-methoxy-1*H*-indole-2-carboxylate (200 mg, 0.912 mmol) in DMF (3 mL) and for 16 h. The reaction mixture was diluted with saturated NaHCO_3_ (20 mL) and extracted with EtOAc (2 × 20 mL). The combined organic layers were washed with brine, dried with anhydrous Na_2_SO_4_, filtered, and concentrated. The crude product was purified using automated flash chromatography (Sfar Silica HC D 25 g column, hexanes/EtOAc gradient mobile phase). Product-containing fractions were concentrated to obtain the desired product (183 mg, 67% yield). ^1^H NMR (500 MHz, CDCl_3_) δ 8.87 (s, 1H), 7.53 (d, *J* = 8.9 Hz, 1H), 6.92 – 6.86 (m, 1H), 6.79 (d, *J* = 2.2 Hz, 1H), 4.44 (q, *J* = 7.1 Hz, 2H), 3.86 (s, 3H), 1.44 (t, *J* = 7.2 Hz, 3H).

***(b)*** ***Synthesis of ethyl 6-methoxy-3-vinyl-1H-indole-2-carboxylate*.** Pd(PPh_3_)_4_ (58.1 mg, 0.050 mmol) was added to a solution of ethyl 3-bromo-6-methoxy-1*H*-indole-2-carboxylate (150 mg, 0.503 mmol), 4,4,6-trimethyl-2-vinyl-1,3,2-dioxaborinane (0.095 mL, 0.553 mmol), and K_2_CO_3_ (139 mg, 1.006 mmol) in degassed dioxane (3 mL) and water (1 mL). The reaction mixture was stirred for 16 h at 90 °C under N_2_. The cooled reaction mixture was diluted with water (10 mL) and extracted with EtOAc (2 × 10 mL). The combined organic layers were washed with brine, dried with anhydrous Na_2_SO_4_, filtered, and concentrated. The crude product was purified using automated flash chromatography (Sfar Silica HC D 25 g column, hexanes/EtOAc gradient mobile phase). Product-containing fractions were concentrated to obtain the desired product (30 mg, 24% yield). ^1^H NMR (500 MHz, CDCl_3_) δ 8.71 (s, 1H), 7.87 (d, *J* = 8.9 Hz, 1H), 7.51 (dd, *J* = 18.0, 11.5 Hz, 1H), 6.89 – 6.77 (m, 2H), 5.92 (dd, *J* = 18.0, 1.6 Hz, 1H), 5.47 (dd, *J* = 11.6, 1.5 Hz, 1H), 4.42 (q, *J* = 7.3 Hz, 2H), 3.86 (d, *J* = 2.1 Hz, 3H), 1.43 (t, *J* = 7.1 Hz, 3H).

***(c)*** ***Synthesis of ethyl 3-ethyl-6-methoxy-1H-indole-2-carboxylate*.** Triethylsilane (0.073 mL, 0.457 mmol) was added dropwise to a solution of ethyl 6-methoxy-3-vinyl-1*H*-indole-2-carboxylate (28 mg, 0.114 mmol), and 10% Pd/C (12.2 mg, 0.011 mmol) stirring under N_2_ in MeOH. The reaction mixture was stirred for 1 h and then filtered through celite. The filtrate was concentrated and purified using automated flash chromatography (Sfar Silica HC D 10 g column, hexanes/EtOAc gradient mobile phase). Product-containing fractions were concentrated to obtain the desired product (26 mg, 92% yield). ^1^H NMR (500 MHz, CDCl_3_) δ 8.57 – 8.53 (m, 1H), 7.55 (ddt, *J* = 8.4, 3.2, 0.9 Hz, 1H), 6.85 – 6.75 (m, 2H), 4.39 (qd, *J* = 7.1, 4.6 Hz, 2H), 3.86 (d, *J* = 2.4 Hz, 3H), 3.08 (q, *J* = 7.6 Hz, 2H), 1.41 (t, *J* = 7.1 Hz, 3H), 1.27 (t, *J* = 7.5 Hz, 3H).

***(d)*** ***Synthesis of 3-ethyl-6-methoxy-1H-indole-2-carboxylic acid*.** A solution of ethyl 3-ethyl-6-methoxy-1*H*-indole-2-carboxylate (24 mg, 0.097 mmol) in 2M KOH (1 mL) and water (1 mL) was stirred overnight at 90 °C. The reaction mixture was acidified with 1M HCl and washed with EtOAc (3 × 10 mL). The combined organic layers were washed with brine, dried with anhydrous Na_2_SO_4_, filtered, and concentrated. The product was used in the next step without further purification. ^1^H NMR (500 MHz, DMSO-*d_6_*) δ 12.60 (s, 1H), 11.14 (s, 1H), 7.51 (d, *J* = 8.8 Hz, 1H), 6.81 (d, *J* = 2.2 Hz, 1H), 6.69 (dd, *J* = 8.7, 2.3 Hz, 1H), 3.76 (s, 3H), 3.00 (q, *J* = 7.5 Hz, 2H), 1.16 (t, *J* = 7.5 Hz, 3H).

***(e) Synthesis of* *N-(2-(2,6-dioxopiperidin-3-yl)-1-oxoisoindolin-5-yl)-3-ethyl-6-methoxy-1H-indole-2-carboxamide*** ***13****.* A solution of 3-(5-amino-1-oxoisoindolin-2-yl)piperidine-2,6-dione (30 mg, 0.116 mmol), 3-ethyl-6-methoxy-1*H*-indole-2-carboxylic acid (25.4 mg, 0.116 mmol), TCFH (35.7 mg, 0.127 mmol), and *N*-methylimidazole (28.5 mg, 0.347 mmol) was stirred for 16 h in NMP (500 µL). The reaction mixture was purified using automated reversed-phase flash chromatography (Biotage Sfar C18 30g column, acetonitrile/water gradient mobile phase with 0.1% formic acid additive). Product-containing fractions were lyophilized to obtain a white solid (8 mg, 15% yield). HRMS (ESI) for C_25_H_25_N_4_O_5_ [M+H]^+^ calc’d 461.1825, obtained 461.1827. ^1^H NMR (500 MHz, DMSO-*d_6_*) δ 11.25 (s, 1H), 10.99 (s, 1H), 10.15 (s, 1H), 8.14 (d, *J* = 1.7 Hz, 1H), 7.77 – 7.68 (m, 2H), 7.56 (d, *J* = 8.7 Hz, 1H), 6.89 (d, *J* = 2.2 Hz, 1H), 6.74 (dd, *J* = 8.8, 2.3 Hz, 1H), 5.11 (dd, *J* = 13.3, 5.1 Hz, 1H), 4.47 (d, *J* = 17.2 Hz, 1H), 4.34 (d, *J* = 17.2 Hz, 1H), 3.81 (s, 3H), 3.06 (q, *J* = 7.5 Hz, 2H), 2.92 (ddd, *J* = 17.2, 13.6, 5.4 Hz, 1H), 2.61 (ddd, *J* = 17.3, 4.6, 2.5 Hz, 1H), 2.40 (qd, *J* = 13.3, 4.6 Hz, 1H), 2.01 (dtd, *J* = 12.7, 5.2, 2.2 Hz, 1H), 1.20 (t, *J* = 7.4 Hz, 3H). ^13^C NMR (126 MHz, DMSO-*d_6_*) δ 172.93, 171.15, 167.88, 160.56, 157.87, 143.30, 142.50, 136.68, 126.39, 125.23, 124.15, 123.65, 121.51, 120.80, 119.37, 113.79, 110.57, 94.08, 55.25, 51.56, 47.19, 31.24, 22.56, 17.55, 15.71.

**Scheme S9. Preparation of Compound 14^a^**

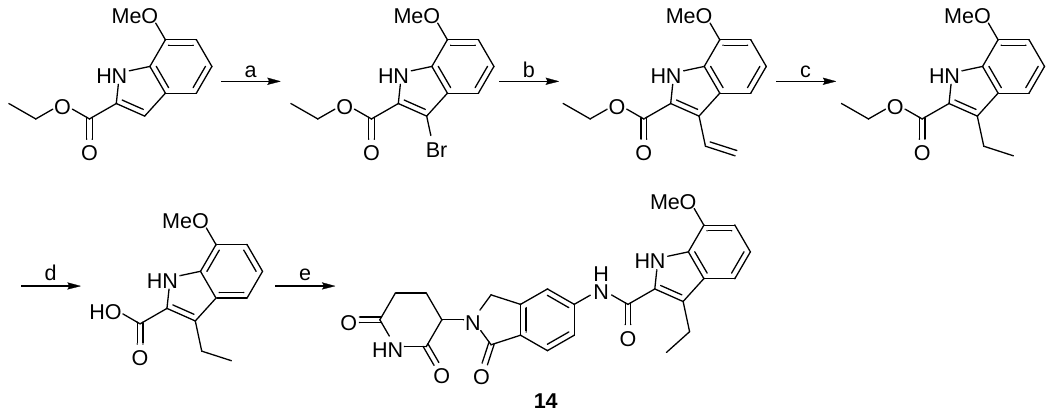

^a^Reagents and conditions: a) *N*-bromosuccinimide, DMF, rt, 16 h; b) 4,4,6-trimethyl-2-vinyl-1,3,2-dioxaborinane, K_2_CO_3_, Pd(PPh_3_)_4_, Dioxane:H_2_O (3:1), 90 °C, 16 h; c) Pd/C, triethylsilane, MeOH, rt, 1 h; d) 1M NaOH, H_2_O, 90 °C, 16 h; e) 3-(5-amino-1-oxoisoindolin-2-yl)piperidine-2,6-dione (1.0 equiv), TCFH (2.0 equiv), *N*-methylimidazole (3.0 equiv), NMP, rt, 16 h.

***(a)*** ***Synthesis of ethyl 3-bromo-7-methoxy-1H-indole-2-carboxylate*.** *N*-bromosuccinimide (170 mg, 0.958 mmol) was added portionwise to a solution of ethyl 7-methoxy-1*H*-indole-2-carboxylate (200 mg, 0.912 mmol) in DMF (3 mL) for 16 h. The reaction mixture was diluted with water (20 mL) and extracted with EtOAc (2 × 20 mL). The combined organic layers were washed with brine, dried with anhydrous Na_2_SO_4_, filtered, and concentrated. The crude product was purified using automated flash chromatography (Sfar Silica HC D 25 g column, hexanes/EtOAc gradient mobile phase). Product-containing fractions were concentrated to obtain the desired product (219 mg, 81% yield). ^1^H NMR (400 MHz, CDCl_3_) δ 9.17 (s, 1H), 7.28 – 7.21 (m, 1H), 7.13 (dd, *J* = 8.2, 7.6 Hz, 1H), 6.76 (dd, *J* = 7.7, 0.8 Hz, 1H), 4.45 (q, *J* = 7.1 Hz, 2H), 3.97 (s, 3H), 1.45 (t, *J* = 7.1 Hz, 3H).

***(b) Synthesis of ethyl 7-methoxy-3-vinyl-1H-indole-2-carboxylate*.** Pd(PPh_3_)_4_ (78 mg, 0.067 mmol) was added to a solution of ethyl 3-bromo-7-methoxy-1*H*-indole-2-carboxylate (200 mg, 0.671 mmol), 4,4,6-trimethyl-2-vinyl-1,3,2-dioxaborinane (0.116 mL, 0.671 mmol), K_2_CO_3_ (185 mg, 1.342 mmol) in degassed dioxane (3 mL) and water (1 mL). The reaction mixture was stirred for 16 h at 90 °C under N_2_. The cooled reaction mixture was diluted with water (10 mL) and extracted with EtOAc (2 × 10 mL). The combined organic layers were washed with brine, dried with anhydrous Na_2_SO_4_, filtered, and concentrated. The crude product was purified using automated flash chromatography (Sfar Silica HC D 25 g column, hexanes/EtOAc gradient mobile phase). Product-containing fractions were concentrated to obtain the desired product (92 mg, 56% yield). ^1^H NMR (400 MHz, CDCl_3_) δ 9.01 (s, 1H), 7.62 – 7.49 (m, 2H), 7.11 (dd, *J* = 8.3, 7.7 Hz, 1H), 6.75 (dd, *J* = 7.8, 0.7 Hz, 1H), 5.95 (dd, *J* = 18.1, 1.6 Hz, 1H), 5.48 (dd, *J* = 11.6, 1.6 Hz, 1H), 4.43 (q, *J* = 7.1 Hz, 2H), 3.98 (s, 3H), 1.44 (t, *J* = 7.1 Hz, 3H).

***(c) Synthesis of ethyl 3-ethyl-7-methoxy-1H-indole-2-carboxylate*.** Triethylsilane (0.156 mL, 0.978 mmol) was added dropwise to a solution of ethyl 7-methoxy-3-vinyl-1*H*-indole-2-carboxylate (80 mg, 0.326 mmol), and 10% Pd/C (34.7 mg, 0.033 mmol) stirring under N_2_ in MeOH. The reaction mixture was stirred for 1 h and then filtered through celite. The filtrate was concentrated and purified using automated flash chromatography (Sfar Silica HC D 10 g column, hexanes/EtOAc gradient mobile phase). Product-containing fractions were concentrated to obtain the desired product (74 mg, 92% yield). ^1^H NMR (400 MHz, CDCl_3_) δ 8.83 (s, 1H), 7.33 – 7.27 (m, 1H), 7.05 (dd, *J* = 8.2, 7.6 Hz, 1H), 6.72 (dd, *J* = 7.6, 0.8 Hz, 1H), 4.42 (q, *J* = 7.1 Hz, 2H), 3.97 (s, 3H), 3.11 (q, *J* = 7.5 Hz, 2H), 1.43 (t, *J* = 7.1 Hz, 3H), 1.28 (t, *J* = 7.5 Hz, 3H).

***(d) Synthesis of 3-ethyl-7-methoxy-1H-indole-2-carboxylic acid*.** A solution of ethyl 3-ethyl-7-methoxy-1*H*-indole-2-carboxylate (50 mg, 0.202 mmol) in 1M NaOH (1 mL) and water (1 mL) was stirred for 16 h at 90 °C. The reaction mixture was acidified with 1 M HCl and washed with EtOAc (3 × 10 mL). The combined organic layers were washed with brine, dried with anhydrous Na_2_SO_4_, filtered, and concentrated. The product was used in the next step without further purification. ^1^H NMR (500 MHz, DMSO-*d_6_*) δ 12.73 (s, 1H), 11.05 (s, 1H), 7.22 (d, *J* = 8.1 Hz, 1H), 6.97 (t, *J* = 7.8 Hz, 1H), 6.76 (d, *J* = 7.6 Hz, 1H), 3.89 (s, 3H), 3.02 (d, *J* = 7.5 Hz, 2H), 1.17 (t, *J* = 7.5 Hz, 3H). ^13^C NMR (126 MHz, DMSO-*d_6_*) δ 163.11, 146.65, 128.44, 126.90, 125.24, 123.56, 119.91, 112.41, 104.35, 55.31, 17.62, 15.71.

***(e) Synthesis of* *N-(2-(2,6-dioxopiperidin-3-yl)-1-oxoisoindolin-5-yl)-3-ethyl-7-methoxy-1H-indole-2-carboxamide* *14****.* A solution of 3-(5-amino-1-oxoisoindolin-2-yl)piperidine-2,6-dione (30 mg, 0.116 mmol), 3-ethyl-7-methoxy-1*H*-indole-2-carboxylic acid (25.4 mg, 0.116 mmol), TCFH (35.7 mg, 0.127 mmol), and *N*-methylimidazole (27.7 µL, 0.347 mmol) was stirred for 16 h in NMP (500 µL). The reaction mixture was purified using automated reversed-phase flash chromatography (Biotage Sfar C18 30g column, acetonitrile/water gradient mobile phase with 0.1% formic acid additive). Product-containing fractions were lyophilized to obtain a white solid (3 mg, 6% yield). HRMS (ESI) for C_25_H_25_N_4_O_5_ [M+H]^+^ calc’d 461.1825, obtained 461.1831. ^1^H NMR (500 MHz, DMSO-*d_6_*) δ 11.52 (s, 1H), 11.00 (s, 1H), 10.33 (s, 1H), 8.19 (d, *J* = 1.7 Hz, 1H), 7.78 (dd, *J* = 8.3, 1.8 Hz, 1H), 7.72 (d, *J* = 8.2 Hz, 1H), 7.26 (d, *J* = 8.1 Hz, 1H), 7.02 (t, *J* = 7.8 Hz, 1H), 6.83 (d, *J* = 7.6 Hz, 1H), 5.11 (dd, *J* = 13.3, 5.1 Hz, 1H), 4.48 (d, *J* = 17.2 Hz, 1H), 4.34 (d, *J* = 17.1 Hz, 1H), 3.97 (s, 3H), 3.11 (q, *J* = 7.4 Hz, 2H), 2.92 (ddd, *J* = 17.3, 13.7, 5.4 Hz, 1H), 2.61 (ddd, *J* = 17.2, 4.5, 2.2 Hz, 1H), 2.46 – 2.34 (m, 1H), 2.02 (ddq, *J* = 10.3, 5.3, 2.7 Hz, 1H), 1.20 (t, *J* = 7.4 Hz, 3H). ^13^C NMR (126 MHz, DMSO-*d_6_*) δ 172.93, 171.15, 167.87, 160.01, 146.41, 143.30, 142.40, 128.46, 126.49, 126.30, 125.60, 125.46, 123.62, 120.13, 119.38, 113.88, 112.24, 104.21, 55.26, 51.57, 47.20, 31.24, 22.56, 17.75, 15.54.

**Scheme S10. Preparation of Compound 15^a^**

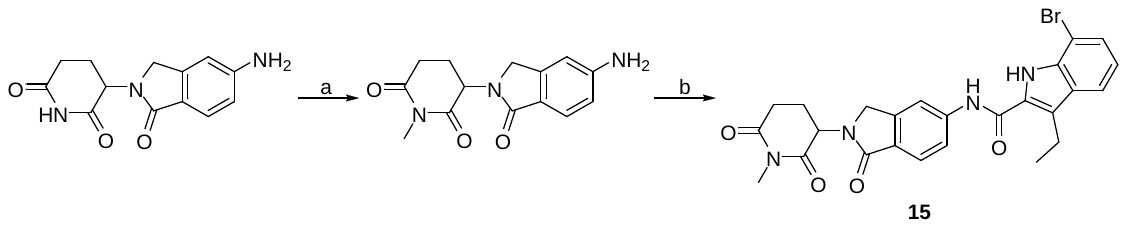

^a^Reagents and conditions: a) CH_3_I (1.5 equiv), Cs_2_CO_3_ (1.2 equiv), DMF, rt, 40 min; b) 7-bromo-3-ethyl-1*H*-indole-2-carboxylic acid (1.0 equiv), TCFH (2.0 equiv), *N*-methylimidazole (3.0 equiv), NMP, rt, 16 h.

***(a) Synthesis of* *3-(5-amino-1-oxoisoindolin-2-yl)-1-methylpiperidine-2,6-dione****.* To a solution of 3-(5-amino-1-oxoisoindolin-2-yl)piperidine-2,6-dione (100 mg, 0.386 mmol) and Cs_2_CO_3_ (151 mg, 0.463 mmol) in anhydrous DMF (2 mL) was added methyl iodide (0.036 mL, 0.579 mmol). The reaction mixture was stirred for 40 min at rt and then diluted with water (15 mL) and extracted with EtOAc (2 × 15 mL). The combined organic layers were washed with brine, dried over anhydrous Na_2_SO_4_, filtered, and concentrated. The crude product was purified using automated flash chromatography (Sfar Silica HC D 25 g column, methanol/dichloromethane gradient mobile phase). Product-containing fractions were concentrated to obtain the desired product (66 mg, 63% yield). ^1^H NMR (400 MHz, DMSO-*d_6_*) δ 7.35 (d, *J* = 8.7 Hz, 1H), 6.66 – 6.57 (m, 2H), 5.80 (s, 2H), 5.07 (dd, *J* = 13.4, 5.0 Hz, 1H), 4.24 (d, *J* = 16.5 Hz, 1H), 4.10 (d, *J* = 16.6 Hz, 1H), 3.04 – 2.92 (m, 4H), 2.73 (s, 1H), 2.33 (qd, *J* = 13.2, 4.6 Hz, 1H), 1.99 – 1.91 (m, 1H).

***(b) Synthesis of* *7-bromo-3-ethyl-N-(2-(1-methyl-2,6-dioxopiperidin-3-yl)-1-oxoisoindolin-5-yl)-1H-indole-2-carboxamide* *15****.* A solution of 3-(5-amino-1-oxoisoindolin-2-yl)-1-methylpiperidine-2,6-dione (20 mg, 0.073 mmol), 7-bromo-3-ethyl-1*H*-indole-2-carboxylic acid (19.6 mg, 0.073 mmol), TCFH (41.1 mg, 0.146 mmol), and *N*-methylimidazole (17.5 µL, 0.220 mmol) was stirred for 2.5 h in NMP (500 µL). The reaction mixture was purified using automated reversed-phase flash chromatography (Biotage Sfar C18 30 g column, acetonitrile/water gradient mobile phase with 0.1% formic acid additive). Product-containing fractions were lyophilized to obtain a white solid (12 mg, 31% yield). HRMS (ESI) for C_25_H_24_BrN_4_O_4_ [M+H]^+^ calc’d 523.0981, obtained 523.0996. ^1^H NMR (500 MHz, DMSO-*d_6_*) δ 11.52 (s, 1H), 10.61 (s, 1H), 8.18 (s, 1H), 7.81 (dd, *J* = 8.2, 1.8 Hz, 1H), 7.77 – 7.71 (m, 2H), 7.52 (dd, *J* = 7.5, 0.9 Hz, 1H), 7.05 (t, *J* = 7.7 Hz, 1H), 5.18 (dd, *J* = 13.4, 5.1 Hz, 1H), 4.49 (d, *J* = 17.2 Hz, 1H), 4.34 (d, *J* = 17.2 Hz, 1H), 3.11 (q, *J* = 7.5 Hz, 2H), 3.02 (s, 4H), 2.77 (ddd, *J* = 17.2, 4.5, 2.4 Hz, 1H), 2.41 (qd, *J* = 13.2, 4.5 Hz, 1H), 2.03 (dtd, *J* = 12.5, 5.2, 2.3 Hz, 1H), 1.21 (t, *J* = 7.4 Hz, 3H). ^13^C NMR (126 MHz, DMSO-*d_6_*) δ 171.94, 170.78, 167.85, 159.82, 143.27, 142.17, 134.21, 128.62, 127.30, 126.99, 126.72, 125.62, 123.65, 120.92, 119.76, 119.40, 114.21, 104.64, 52.10, 47.20, 31.39, 26.59, 21.80, 17.69, 15.48.

**Scheme S11. Preparation of Compound 16^a^**

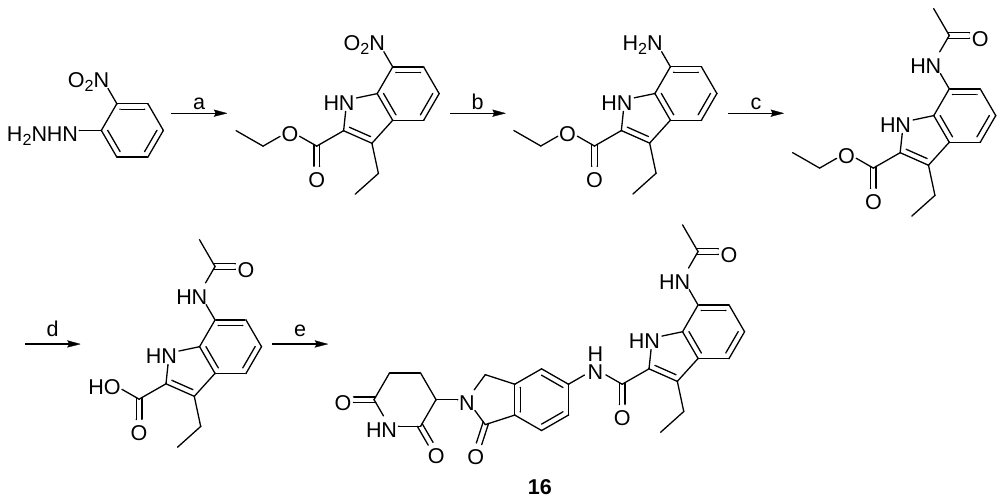

^a^Reagents and conditions: a) ethyl 2-oxopentanoate, *p*-toluenesulfonic acid monohydrate, EtOH, 90 °C, 16 h, then polyphosphoric acid, 90 °C, 1 h; b) Pd/C, triethylsilane, MeOH, rt, 1 h; c) Ac_2_O, pyridine, DCM, rt, 30 min; d) 2M KOH, EtOH, 60 °C, 1 h; e) 3-(5-amino-1-oxoisoindolin-2-yl)piperidine-2,6-dione (1.0 equiv), TCFH (2.0 equiv), *N*-methylimidazole (3.0 equiv), NMP, rt, 1 h.

***(a) Synthesis of ethyl 3-ethyl-7-nitro-1H-indole-2-carboxylate*.** To a stirring solution of (2-nitrophenyl)hydrazine hydrochloride (300 mg, 1.582 mmol) and ethyl 2-oxopentanoate (228 mg, 1.582 mmol) in EtOH (10 mL) was added *p*-toluenesulfonic acid monohydrate (903 mg, 4.75 mmol) and heated to 90 °C. After 3 h additional *p*-toluenesulfonic acid monohydrate (903 mg, 4.75 mmol) was added and the mixture stirred for 16 h. The reaction mixture was allowed to cool to rt, diluted with EtOAc (50 mL) and then washed with saturated NaHCO_3_ (50 mL). The organic layer was washed with brine, dried with anhydrous Na_2_SO_4_, filtered, and concentrated. To the crude mixture was added polyphosphoricacid (2719 mg, 11.33 mmol) and the solution was heated to 90 °C. After 1 h, the reaction mixture was allowed to cool to rt, diluted with water (20 mL) and washed with EtOAc (3 × 20 mL). The combined organic layers were washed with brine, dried with anhydrous Na_2_SO_4_, filtered, and concentrated. The crude mixture was purified by flash column chromatography (Biotage Isolera, 25 g Sfar column, 0–20% hexanes:EtOAc) to give the desired product (137 mg, 69% yield). ^1^H NMR (500 MHz, CDCl_3_) δ 10.14 (s, 1H), 8.29 (d, *J* = 8.0 Hz, 1H), 8.08 – 8.02 (m, 1H), 7.24 (d, *J* = 8.0 Hz, 1H), 4.47 (q, *J* = 7.1 Hz, 2H), 3.16 (q, *J* = 7.5 Hz, 2H), 1.46 (t, *J* = 7.1 Hz, 3H), 1.29 (t, *J* = 7.5 Hz, 3H). ^13^C NMR (126 MHz, CDCl_3_) δ 161.41, 133.42, 131.60, 129.07, 129.03, 127.72, 125.39, 122.61, 119.37, 61.40, 18.08, 15.60, 14.54.

***(b) Synthesis of ethyl 7-amino-3-ethyl-1H-indole-2-carboxylate*.** Triethylsilane (0.372 mL, 2.326 mmol) was added dropwise to a solution of ethyl 3-ethyl-7-nitro-1*H*-indole-2-carboxylate (122 mg, 0.465 mmol), and 10% Pd/C (49.5 mg, 0.047 mmol) stirring under N_2_ in MeOH (2 mL). The reaction mixture was stirred for 1 h and then filtered through celite and concentrated to give the desired product (123 mg), which was used in the next step without further purification. ^1^H NMR (500 MHz, CDCl_3_) δ 9.31 (s, 1H), 7.19 (d, *J* = 8.1 Hz, 1H), 6.98 (t, *J* = 7.7 Hz, 1H), 6.66 (d, *J* = 7.3 Hz, 1H), 4.44 (q, *J* = 7.1 Hz, 2H), 4.22 – 3.33 (m, 2H), 3.09 (q, *J* = 7.5 Hz, 2H), 1.51 – 1.40 (m, 3H), 1.27 (t, *J* = 7.4 Hz, 3H). ^13^C NMR (126 MHz, CDCl_3_) δ 163.49, 132.01, 128.86, 127.83, 127.67, 122.72, 120.98, 111.91, 110.43, 61.04, 18.59, 15.57, 14.51.

***(c) Synthesis of ethyl 7-acetamido-3-ethyl-1H-indole-2-carboxylate*.** To a stirring solution of ethyl 7-amino-3-ethyl-1*H*-indole-2-carboxylate (100 mg, 0.431 mmol) in DCM (2 mL), was added pyridine (0.174 mL, 2.153 mmol), and acetic anhydride (0.081 mL, 0.861 mmol). The reaction mixture was stirred for 30 min and then diluted with water (10 mL) and washed with EtOAc (3 × 10 mL). The combined organic layers were washed with brine, dried with anhydrous Na_2_SO_4_, filtered, and concentrated. The product (131 mg) was used in the next step without further purification. ^1^H NMR (500 MHz, CDCl_3_) δ 10.04 (s, 1H), 7.68 (s, 1H), 7.52 (d, *J* = 7.5 Hz, 1H), 7.11 – 6.97 (m, 2H), 4.41 (q, *J* = 7.1 Hz, 2H), 3.11 (q, *J* = 7.5 Hz, 2H), 2.29 (s, 3H), 1.41 (t, *J* = 7.2 Hz, 3H), 1.26 (t, *J* = 7.5 Hz, 3H). ^13^C NMR (126 MHz, CDCl_3_) δ 168.84, 162.59, 149.76, 130.13, 128.98, 127.15, 123.98, 123.42, 123.15, 119.71, 118.14, 116.62, 60.86, 24.30, 18.23, 15.55, 14.55.

***(d) Synthesis of 7-acetamido-3-ethyl-1H-indole-2-carboxylic acid*.** A solution of ethyl 7-acetamido-3-ethyl-1*H*-indole-2-carboxylate (125 mg, 0.456 mmol) in 2M KOH (1.8 mL) and EtOH (2 mL) was stirred at 60 °C. After 1 h, the reaction mixture was cooled to rt and then directly purified using automated reversed-phase flash chromatography (Biotage Sfar C18 30 g column, acetonitrile/water gradient mobile phase with 0.1% formic acid additive). Product-containing fractions were lyophilized to obtain a white solid (59 mg, 53%). ^1^H NMR (500 MHz, DMSO-*d_6_*) δ 13.17 (s, 1H), 11.21 (s, 1H), 9.90 (s, 1H), 7.93 (d, *J* = 7.6 Hz, 1H), 7.36 (d, *J* = 8.0 Hz, 1H), 6.98 (t, *J* = 7.8 Hz, 1H), 3.04 (q, *J* = 7.4 Hz, 2H), 2.16 (s, 3H), 1.18 (t, *J* = 7.4 Hz, 3H). ^13^C NMR (126 MHz, DMSO-*d_6_*) δ 168.49, 163.58, 128.33, 126.85, 124.59, 119.43, 115.28, 114.11, 23.98, 17.40, 15.74.

***(e) Synthesis of* *7-acetamido-N-(2-(2,6-dioxopiperidin-3-yl)-1-oxoisoindolin-5-yl)-3-ethyl-1H-indole-2-carboxamide* *16****.* A solution of 3-(5-amino-1-oxoisoindolin-2-yl)piperidine-2,6-dione (25 mg, 0.096 mmol), 7-acetamido-3-ethyl-1*H*-indole-2-carboxylic acid (23.8 mg, 0.096 mmol), TCFH (54.1 mg, 0.193 mmol), and *N*-methylimidazole (23.1 µL, 0.289 mmol) was stirred for 1 h in NMP (500 µL). The reaction mixture was purified using automated reversed-phase flash chromatography (Biotage Sfar C18 30 g column, acetonitrile/water gradient mobile phase with 0.1% formic acid additive). Product-containing fractions were lyophilized to obtain a white solid (23 mg, 49% yield). HRMS (ESI) for C_26_H_26_N_5_O_5_ [M+H]^+^ calc’d 488.1934, obtained 488.1930. ^1^H NMR (500 MHz, DMSO-*d_6_*) δ 11.12 (s, 1H), 11.00 (s, 1H), 10.47 (s, 1H), 9.84 (s, 1H), 8.13 (s, 1H), 7.88 – 7.76 (m, 1H), 7.73 (d, *J* = 8.2 Hz, 1H), 7.68 (d, *J* = 7.6 Hz, 1H), 7.45 (d, *J* = 8.0 Hz, 1H), 7.04 (t, *J* = 7.8 Hz, 1H), 5.11 (dd, *J* = 13.3, 5.1 Hz, 1H), 4.48 (d, *J* = 17.2 Hz, 1H), 4.34 (d, *J* = 17.2 Hz, 1H), 3.07 (q, *J* = 7.5 Hz, 2H), 2.92 (ddd, *J* = 18.0, 13.6, 5.4 Hz, 1H), 2.68 – 2.57 (m, 1H), 2.40 (qd, *J* = 13.1, 4.3 Hz, 1H), 2.17 (s, 3H), 2.02 (ddd, *J* = 11.1, 6.3, 4.1 Hz, 1H), 1.21 (t, *J* = 7.5 Hz, 3H). ^13^C NMR (126 MHz, DMSO-*d_6_*) δ 172.93, 171.14, 168.47, 167.84, 160.89, 143.27, 142.22, 128.37, 127.95, 126.72, 126.66, 124.18, 123.68, 122.91, 119.73, 119.70, 115.83, 114.23, 51.58, 47.20, 31.24, 23.83, 22.55, 17.56, 15.71.

**Scheme S12. Preparation of Compound 17–18^a^**

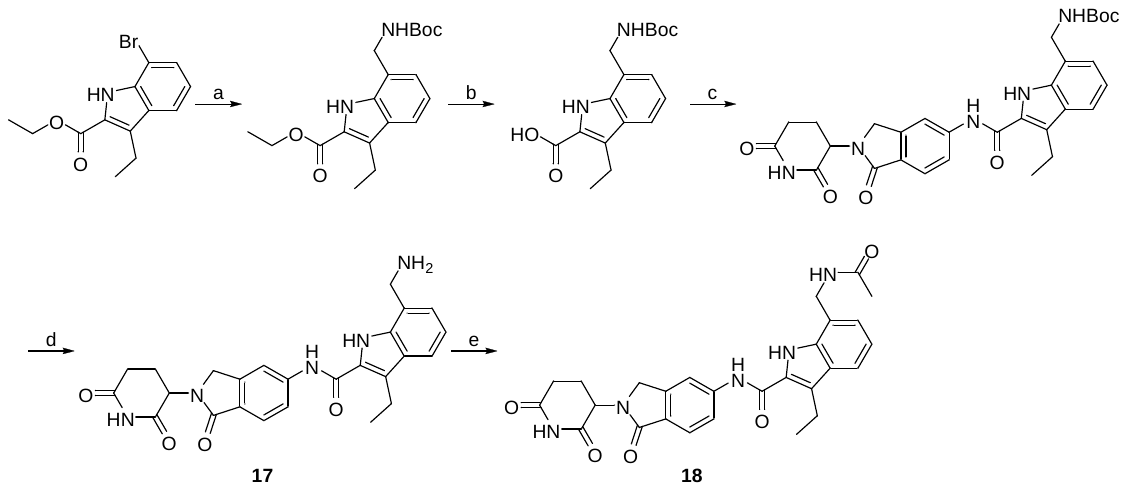

^a^Reagents and conditions: a) potassium *N*-Boc-aminomethyltrifluoroborate, Pd(OAc)_2_, XPhos, K_2_CO_3_, toluene:H_2_O (4:1), 90 °C, 16 h; b) 2M KOH, EtOH, 60 °C, 1 h; c) 3-(5-amino-1-oxoisoindolin-2-yl)piperidine-2,6-dione (1.0 equiv), TCFH (2.0 equiv), *N*-methylimidazole (3.0 equiv), NMP, rt, 90 min; d) 4M HCl in dioxane (8.0 equiv), rt, 5 h; e) Ac_2_O (2.0 equiv), pyridine (5.0 equiv), DCM, rt, 30 min.

***(a)*** ***Synthesis of ethyl 7-(((tert-butoxycarbonyl)amino)methyl)-3-ethyl-1H-indole-2-carboxylate*.** To a round-bottom flask was added ethyl 7-bromo-3-ethyl-1*H*-indole-2-carboxylate (100 mg, 0.338 mmol), potassium *N*-Boc-aminomethyltrifluoroborate (84 mg, 0.355 mmol), Pd(OAc)_2_ (3.79 mg, 0.017 mmol), XPhos (16.10 mg, 0.034 mmol), and K_2_CO_3_ (140 mg, 1.013 mmol). The flask was flushed with N_2_ and then toluene (1 mL) and water (0.250 mL) were added. The reaction mixture was stirred at 90 °C for 16 h. The solution was allowed to cool to rt and diluted with water (15 mL) and washed with EtOAc (2 × 15 mL). The combined organic layers were washed with brine, dried with anhydrous Na_2_SO_4_, filtered, and concentrated. The crude mixture was purified by flash column chromatography (Biotage Isolera, 10 g Sfar column, 0–25% hexanes:EtOAc) to give the desired product (55 mg, 47% yield). ^1^H NMR (500 MHz, CDCl_3_) δ 10.33 (s, 1H), 7.65 (d, *J* = 7.9 Hz, 1H), 7.08 (d, *J* = 6.9 Hz, 1H), 7.03 (t, *J* = 7.5 Hz, 1H), 5.06 (t, *J* = 6.8 Hz, 1H), 4.54 (d, *J* = 6.8 Hz, 2H), 4.41 (q, *J* = 7.1 Hz, 2H), 3.15 (q, *J* = 7.5 Hz, 2H), 1.48 (s, 9H), 1.44 (t, *J* = 7.1 Hz, 3H), 1.28 (t, *J* = 7.5 Hz, 3H). ^13^C NMR (126 MHz, CDCl_3_) δ 162.14, 157.53, 134.73, 127.99, 127.22, 125.09, 123.41, 123.18, 120.80, 119.44, 80.55, 60.52, 42.39, 28.50, 18.19, 15.59, 14.57.

***(b)*** ***Synthesis of 7-(((tert-butoxycarbonyl)amino)methyl)-3-ethyl-1H-indole-2-carboxylic acid*.** A solution of ethyl 7-(((*tert*-butoxycarbonyl)amino)methyl)-3-ethyl-1*H*-indole-2-carboxylate (50 mg, 0.144 mmol) in 2M KOH (0.6 mL) and EtOH (1 mL) was stirred at 90 °C. After 4 h, the reaction mixture was cooled to rt and then directly purified using automated reversed-phase flash chromatography (Biotage Sfar C18 30 g column, acetonitrile/water gradient mobile phase with 0.1% formic acid additive). Product-containing fractions were lyophilized to obtain a white solid (33 mg, 72% yield). ^1^H NMR (500 MHz, DMSO-*d_6_*) δ 12.97 (s, 1H), 10.87 (s, 1H), 7.60 – 7.49 (m, 2H), 7.10 (d, *J* = 7.0 Hz, 1H), 7.01 (t, *J* = 7.5 Hz, 1H), 4.36 (d, *J* = 6.3 Hz, 2H), 3.05 (d, *J* = 7.5 Hz, 2H), 1.39 (s, 9H), 1.18 (t, *J* = 7.4 Hz, 3H). ^13^C NMR (126 MHz, DMSO-*d_6_*) δ 163.28, 156.35, 134.12, 127.26, 124.07, 124.01, 123.57, 119.28, 119.24, 78.25, 40.67, 28.20, 17.45, 15.77.

***(c) Synthesis of* *tert-butyl ((2-((2-(2,6-dioxopiperidin-3-yl)-1-oxoisoindolin-5-yl)carbamoyl)-3-ethyl-1H-indol-7-yl)methyl)carbamate****.* A solution of 3-(5-amino-1-oxoisoindolin-2-yl)piperidine-2,6-dione (26 mg, 0.100 mmol), 7-(((*tert*-butoxycarbonyl)amino)methyl)-3-ethyl-1*H*-indole-2-carboxylic acid (31.9 mg, 0.100 mmol), TCFH (56.3 mg, 0.201 mmol), and *N*-methylimidazole (24.7 mg, 0.301 mmol) was stirred for 90 min in NMP (500 µL). The reaction mixture was purified using automated reversed-phase flash chromatography (Biotage Sfar C18 30 g column, acetonitrile/water gradient mobile phase with 0.1% formic acid additive). Product-containing fractions were lyophilized to obtain a white solid (33 mg, 59% yield). ^1^H NMR (500 MHz, DMSO-*d_6_*) δ 11.17 (s, 1H), 11.00 (s, 1H), 10.39 (s, 1H), 8.15 (s, 1H), 7.82 – 7.71 (m, 2H), 7.58 (d, *J* = 7.9 Hz, 1H), 7.48 (t, *J* = 6.2 Hz, 1H), 7.13 (d, *J* = 7.1 Hz, 1H), 7.06 (t, *J* = 7.5 Hz, 1H), 5.11 (dd, *J* = 13.3, 5.1 Hz, 1H), 4.52 – 4.44 (m, 3H), 4.34 (d, *J* = 17.2 Hz, 1H), 3.08 (q, *J* = 7.4 Hz, 2H), 2.92 (ddd, *J* = 18.0, 13.7, 5.4 Hz, 1H), 2.65 – 2.58 (m, 1H), 2.39 (td, *J* = 13.1, 4.3 Hz, 1H), 2.06 – 1.98 (m, 1H), 1.40 (s, 9H), 1.21 (t, *J* = 7.5 Hz, 3H). ^13^C NMR (126 MHz, DMSO-*d_6_*) δ 172.93, 171.15, 167.85, 160.69, 156.04, 143.30, 142.29, 133.73, 127.13, 126.63, 123.83, 123.66, 122.41, 119.61, 119.52, 118.57, 114.06, 78.06, 51.58, 47.20, 31.24, 28.24, 22.55, 17.52, 15.74.

***(d) Synthesis of* *7-(aminomethyl)-N-(2-(2,6-dioxopiperidin-3-yl)-1-oxoisoindolin-5-yl)-3-ethyl-1H-indole-2-carboxamide*** ***17****.* HCl (4M in dioxane, 0.09 mL, 0.357 mmol) was added to a stirring solution of *tert*-butyl ((2-((2-(2,6-dioxopiperidin-3-yl)-1-oxoisoindolin-5-yl)carbamoyl)-3-ethyl-1*H*-indol-7-yl)methyl)carbamate (25 mg, 0.045 mmol) in Dioxane (1 mL). After 5 h, the solution was concentrated to give the desired product. HRMS (ESI) for C_25_H_26_N_5_O_4_ [M+H]^+^ calc’d 460.1985, obtained 460.1979. ^1^H NMR (500 MHz, DMSO-*d_6_*) δ 12.41 (s, 1H), 11.00 (s, 1H), 10.95 (s, 1H), 8.35 (t, *J* = 4.7 Hz, 4H), 8.03 (dd, *J* = 8.4, 1.8 Hz, 1H), 7.73 (dd, *J* = 8.2, 3.0 Hz, 2H), 7.37 (d, *J* = 7.2 Hz, 1H), 7.14 (t, *J* = 7.6 Hz, 1H), 5.11 (dd, *J* = 13.3, 5.1 Hz, 1H), 4.55 – 4.45 (m, 3H), 4.34 (d, *J* = 17.2 Hz, 1H), 3.16 (q, *J* = 7.4 Hz, 2H), 2.93 (ddd, *J* = 18.0, 13.6, 5.4 Hz, 1H), 2.65 – 2.58 (m, 1H), 2.41 (qd, *J* = 13.5, 4.7 Hz, 1H), 2.06 – 1.97 (m, 1H), 1.21 (t, *J* = 7.5 Hz, 3H). ^13^C NMR (126 MHz, DMSO-*d_6_*) δ 172.94, 171.15, 167.89, 160.30, 143.22, 142.57, 134.21, 127.50, 126.58, 126.34, 125.59, 125.03, 123.55, 120.58, 119.53, 119.47, 117.88, 113.90, 66.36, 51.58, 47.23, 31.24, 22.55, 17.50, 15.66.

***(e) Synthesis of* *7-(acetamidomethyl)-N-(2-(2,6-dioxopiperidin-3-yl)-1-oxoisoindolin-5-yl)-3-ethyl-1H-indole-2-carboxamide*** ***18****.* To a stirring solution of **17** (17 mg, 0.034 mmol) in DCM (1 mL) was added pyridine (13.9 µL, 0.171 mmol), and acetic anhydride (6.5 µL, 0.069 mmol). After 30 min, the reaction mixture was purified using automated reversed-phase flash chromatography (Biotage Sfar C18 30 g column, acetonitrile/water gradient mobile phase with 0.1% formic acid additive). Product-containing fractions were lyophilized to obtain the desired product (7 mg, 41%). HRMS (ESI) for C_27_H_28_N_5_O_5_ [M+H]^+^ calc’d 502.2090, obtained 502.2087. ^1^H NMR (500 MHz, DMSO-*d_6_*) δ 11.20 (s, 1H), 11.00 (s, 1H), 10.37 (s, 1H), 8.47 (t, *J* = 5.8 Hz, 1H), 8.16 (s, 1H), 7.80 – 7.71 (m, 2H), 7.59 (d, *J* = 8.0 Hz, 1H), 7.15 (d, *J* = 7.1 Hz, 1H), 7.06 (t, *J* = 7.5 Hz, 1H), 5.11 (dd, *J* = 13.3, 5.1 Hz, 1H), 4.58 (d, *J* = 5.8 Hz, 2H), 4.48 (d, *J* = 17.2 Hz, 1H), 4.35 (d, *J* = 17.2 Hz, 1H), 3.09 (q, *J* = 7.4 Hz, 2H), 2.92 (ddd, *J* = 18.1, 13.7, 5.4 Hz, 1H), 2.65 – 2.58 (m, 1H), 2.40 (qd, *J* = 13.2, 4.4 Hz, 1H), 2.02 (ddd, *J* = 13.9, 6.7, 4.5 Hz, 1H), 1.92 (s, 3H), 1.21 (t, *J* = 7.4 Hz, 3H). ^13^C NMR (126 MHz, DMSO-*d_6_*) δ 172.93, 171.15, 169.81, 167.85, 160.63, 143.31, 142.27, 133.94, 127.15, 126.64, 126.58, 123.85, 123.67, 123.18, 122.85, 119.62, 119.54, 118.74, 114.09, 51.58, 47.21, 31.24, 22.61, 22.55, 17.53, 15.72.

**Scheme S13. Preparation of Compound 19^a^**

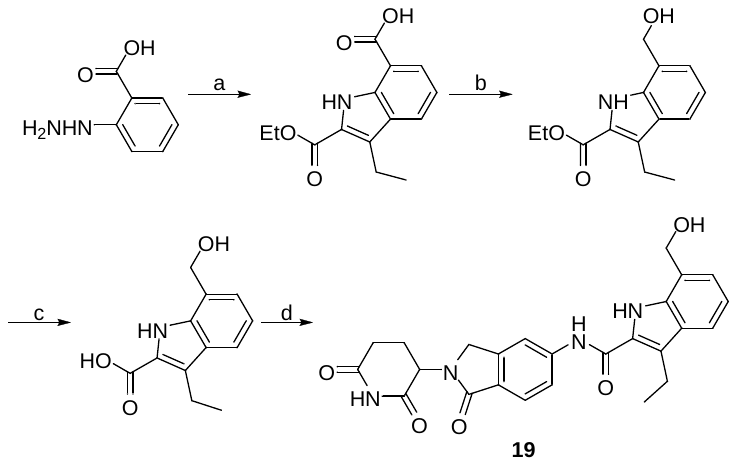

^a^Reagents and conditions: a) ethyl 2-oxopentanoate, H_2_SO_4_, AcOH, 95 °C, 12 h; b) BH_3_/Me_2_S, THF, 0–20 °C, 2 h; c) LiOH, MeOH, H_2_O, 50 °C, 12 h; d) 3-(5-amino-1-oxoisoindolin-2-yl)piperidine-2,6-dione (1.0 equiv), TCFH (2.0 equiv), *N*-methylimidazole (3.0 equiv), NMP, rt, 1 h.

***(a)*** ***Synthesis of 2-(ethoxycarbonyl)-3-ethyl-1H-indole-7-carboxylic acid*.** To a solution of 2-hydrazineylbenzoic acid (5 g, 26.51 mmol, 1 eq, HCl) in AcOH (100 mL) was added H_2_SO_4_ (3.90 g, 39.76 mmol, 2.12 mL, 1.5 eq) and ethyl 2-oxopentanoate (4.59 g, 31.81 mmol, 1.2 eq). The mixture was stirred at 95 °C for 12 h. LC-MS showed the reaction was completed and the desired MS was detected. The reaction was filtered and concentrated under reduced pressure to give a residue. The residue was diluted with ice cold water, adjusted to pH = 9 with saturated NaHCO_3_ (100 mL) and extracted with EtOAc 150 mL (50 mL × 3). The combined organic layers were dried over sodium sulfate, filtered, concentrated to give the crude product (6 g, brown solid), which was used into the next step without further purification.

***(b)*** ***Synthesis of ethyl 3-ethyl-7-(hydroxymethyl)-1H-indole-2-carboxylate*.** 2-(Ethoxycarbonyl)-3-ethyl-1*H*-indole-7-carboxylic acid (1.8 g, 6.89 mmol, 1 eq) was placed in a 250 mL round bottom flask and dissolved in THF (10 mL) under nitrogen atmosphere. The reaction flask was placed in an ice bath to reach 0 °C and BH_3_-Me_2_S (10 M, 3.44 mL, 5 eq) was added dropwise. After 15 min at 0 °C, the ice bath was removed, and the reaction was stirred at 20 °C for 2 h. After reaction completion, ice water (50 ml) was carefully added to the reaction, and the mixture extracted with diethyl ether (50 mL × 3). The reaction mixture was concentrated under reduced pressure to give the crude product (1.5 g, yellow oil) which was used in the next step without further purification.

***(c)*** ***Synthesis of 3-ethyl-7-(hydroxymethyl)-1H-indole-2-carboxylic acid*.** To a solution of ethyl 3-ethyl-7-(hydroxymethyl)-1*H*-indole-2-carboxylate (1 g, 4.04 mmol, 1 eq) in MeOH (5 mL) and H_2_O (5 mL) was added LiOH (4 M, 5.05 mL, 5 eq). The mixture was stirred at 50 °C for 1 h. The reaction was concentrated under reduced pressure to give a residue. The residue was added to HCl (1M, 150 mL) and the resulting precipitate was collected by filtration. The crude product was triturated with MTBE and MeCN at 25 °C for 30 min to give the desired product (310 mg, 1.17 mmol, 28.8% yield, 96.1% purity, HCl) as a white solid. ^1^H NMR (400 MHz, DMSO-*d_6_*): δ 12.86 (br s, 1H), 10.85 (s, 1H), 7.55 (d, J = 8.0 Hz, 1H), 7.22 (d, J = 7.0 Hz, 1H), 7.03 (t, J = 7.6 Hz, 1H), 5.22 (t, J = 5.9 Hz, 1H), 4.79 (d, J = 5.8 Hz, 2H), 3.05 (q, J = 7.5 Hz, 2H), 1.18 (t, J = 7.4 Hz, 1H).

***(d) Synthesis of* *N-(2-(2,6-dioxopiperidin-3-yl)-1-oxoisoindolin-5-yl)-3-ethyl-7-(hydroxymethyl)-1H-indole-2-carboxamide* *19****.* A solution of 3-(5-amino-1-oxoisoindolin-2-yl)piperidine-2,6-dione (25 mg, 0.096 mmol), 3-ethyl-7-(hydroxymethyl)-1*H*-indole-2-carboxylic acid (21.1 mg, 0.096 mmol), TCFH (54.1 mg, 0.193 mmol), and *N*-methylimidazole (23.8 mg, 0.289 mmol) was stirred for 1 h in NMP (500 µL). The reaction mixture was purified using automated reversed-phase flash chromatography (Biotage Sfar C18 30 g column, acetonitrile/water gradient mobile phase with 0.1% formic acid additive). Product-containing fractions were lyophilized to obtain a white solid (23 mg, 52% yield). HRMS (ESI) for C_25_H_25_N_4_O_5_ [M+H]^+^ calc’d 461.1825, obtained 461.1827. ^1^H NMR (500 MHz, DMSO-*d_6_*) δ 11.12 (s, 1H), 11.00 (s, 1H), 10.37 (s, 1H), 8.16 (s, 1H), 7.84 – 7.70 (m, 2H), 7.58 (d, *J* = 8.0 Hz, 1H), 7.27 (d, *J* = 7.0 Hz, 1H), 7.07 (t, *J* = 7.5 Hz, 1H), 5.28 (t, *J* = 5.8 Hz, 1H), 5.11 (dd, *J* = 13.3, 5.1 Hz, 1H), 4.84 (d, *J* = 5.6 Hz, 2H), 4.48 (d, *J* = 17.2 Hz, 1H), 4.35 (d, *J* = 17.1 Hz, 1H), 3.11 (q, *J* = 7.4 Hz, 2H), 2.92 (ddd, *J* = 18.0, 13.6, 5.4 Hz, 1H), 2.61 (dd, *J* = 17.6, 4.0 Hz, 1H), 2.40 (qd, *J* = 13.2, 4.4 Hz, 1H), 2.02 (ddd, *J* = 10.6, 5.5, 3.3 Hz, 1H), 1.21 (t, *J* = 7.4 Hz, 3H). ^13^C NMR (126 MHz, DMSO-*d_6_*) δ 172.94, 171.16, 167.87, 160.42, 143.30, 142.30, 133.63, 127.12, 126.61, 126.35, 126.05, 124.55, 123.64, 122.50, 119.63, 119.44, 118.62, 114.12, 60.07, 51.59, 47.21, 31.24, 22.56, 17.60, 15.68.

**Scheme S14. Preparation of Compound 20^a^**

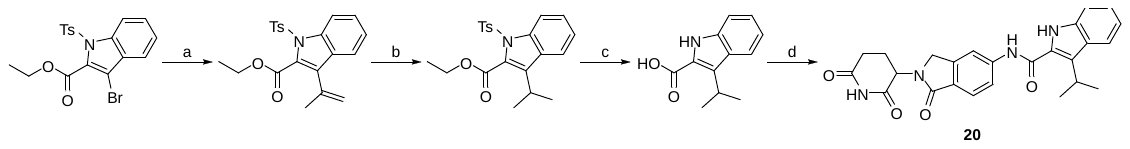

^a^Reagents and conditions: a) 4,4,5,5-tetramethyl-2-(prop-1-en-2-yl)-1,3,2-dioxaborolane (1.05 equiv), Ph(PPh_3_)_4_ (0.1 equiv), K_2_CO_3_ (2 equiv), dioxane:water (3:1), 90 °C, 1 h; b) 10% Pd/C (0.1 equiv), triethylsilane (3 equiv), EtOAc, rt, 16 h; c) 2M KOH, EtOH, 80 °C, 1 h; d) 3-(5-amino-1-oxoisoindolin-2-yl)piperidine-2,6-dione (1 equiv), TCFH (1.1 equiv), *N*-methylimidazole (3 equiv), NMP, rt, 3 h.

**(a)** ***Synthesis of ethyl 3-(prop-1-en-2-yl)-1-tosyl-1H-indole-2-carboxylate.*** A mixture of ethyl 3-bromo-1-tosyl-1*H*-indole-2-carboxylate (0.22 g, 0.52 mmol), 2-isopropenylboronic ester (0.10 mL, 0.55 mmol), potassium carbonate (0.14 g, 1.0 mmol), and Pd(Ph_3_)_4_ (0.060 g, 0.052 mmol) in 3:1 dioxane/water (4 mL) was degassed then stirred at 90 °C under nitrogen atmosphere for 1 h, then cooled to room temperature. The reaction mixture was diluted with ethyl acetate (15 mL) and washed with brine (3 × 15 mL). The organic phase was dried over sodium sulfate, filtered, and concentrated *in vacuo*. The crude was purified using automated flash chromatography (Biotage Sfar Silica HC D 50g column, ethyl acetate/hexanes gradient mobile phase). Product-containing fractions were evaporated *in vacuo* to obtain a colorless oil (0.14 g, 70% yield). ^1^H NMR (500 MHz, CDCl_3_) δ 8.02 (d, *J* = 8.4 Hz, 1H), 7.87 (d, *J* = 7.4 Hz, 2H), 7.52 (d, *J* = 7.9 Hz, 1H), 7.37 (t, *J* = 7.8 Hz, 1H), 7.26 – 7.19 (m, 3H), 5.33 (s, 1H), 5.12 (s, 1H), 4.45 (q, *J* = 7.1 Hz, 2H), 2.34 (s, 3H), 2.09 (s, 3H), 1.41 (t, *J* = 7.1 Hz, 3H). ^13^C NMR (126 MHz, CDCl_3_) δ 162.86, 145.02, 135.96, 135.55, 134.69, 129.61, 128.89, 128.34, 127.41, 126.35, 124.02, 121.31, 117.86, 115.00, 62.32, 23.17, 21.64, 13.98.

**(b) *Synthesis of ethyl 3-isopropyl-1-tosyl-1H-indole-2-carboxylate.*** To a microwave vial fitted with a magnetic stir bar was added 10% Pd/C (0.007 g, 6.52 µmol). The vial was capped and purged with nitrogen. Ethyl 3-(prop-1-en-2-yl)-1-tosyl-1*H*-indole-2-carboxylate (0.025 g, 0.065 mmol) was diluted in ethyl acetate (1 mL) and added to the microwave vial via syringe. The microwave vial was then degassed and purged with nitrogen. Triethylsilane (0.031 mL, 0.20 mmol) was added to the microwave vial dropwise at room temperature. After stirring at room temperature for 2 h an additional molar equivalent of triethylsilane (0.010 mL, 0.065 mmol) was added. After stirring at room temperature for a total of 18 h the reaction mixture was filtered through celite and the filtrate was concentrated *in vacuo*. The crude was purified using automated flash chromatography (Biotage Sfar HC D 10g column, ethyl acetate/hexanes gradient mobile phase). Product-containing fractions were evaporated *in vacuo* to obtain a colorless oil (0.024 g, 95% yield). ^1^H NMR (500 MHz, CDCl_3_) δ 8.01 (d, *J* = 8.4 Hz, 1H), 7.75 (d, *J* = 8.3 Hz, 2H), 7.60 (d, *J* = 8.0 Hz, 1H), 7.34 (t, *J* = 7.8 Hz, 1H), 7.21 (t, *J* = 7.6 Hz, 1H), 7.16 (d, *J* = 8.2 Hz, 2H), 4.47 (q, *J* = 7.2 Hz, 2H), 3.29 (p, *J* = 7.1 Hz, 1H), 2.31 (s, 3H), 1.43 (t, *J* = 7.2 Hz, 3H), 1.34 (d, *J* = 7.1 Hz, 6H). ^13^C NMR (126 MHz, CDCl_3_) δ 163.40, 144.79, 136.90, 134.15, 133.06, 129.39, 128.98, 127.25, 126.08, 123.70, 121.60, 115.64, 62.25, 26.02, 21.91, 21.59, 14.07.

**(c) *Synthesis of 3-isopropyl-1H-indole-2-carboxylic acid.*** To a solution of ethyl 3-isopropyl-1-tosyl-1*H*-indole-2-carboxylate (0.035 g, 0.016 mmol) in ethanol (1.5 mL) was added 2M potassium hydroxide (0.36 mL, 0.73 mmol). The reaction mixture was stirred at 80 °C for 1 h, then cooled to room temperature. The crude was concentrated *in vacuo* to remove volatiles, then was purified using automated reversed-phase flash chromatography (Biotage Sfar C18 12g column, acetonitrile/water gradient mobile phase with 0.1% formic acid additive). Product-containing fractions were lyophilized to obtain a white solid (8 mg, 43% yield).

**(d) *Synthesis of N-(2-(2,6-dioxopiperidin-3-yl)-1-oxoisoindolin-5-yl)-3-isopropyl-1H-indole-2-carboxamide 20.*** A mixture of 3-(5-amino-1-oxoisoindolin-2-yl)piperidine-2,6-dione (0.006 g, 0.025 mmol), 3-isopropyl-1*H*-indole-2-carboxylic acid (0.005 g, 0.025 mmol), TCFH (0.008 g, 0.027 mmol) and NMI (0.006 mL, 0.074 mmol) was stirred at room temperature in NMP (0.5 mL) for 3 h. The reaction mixture was diluted with ethyl acetate (3 mL) and washed with brine (3 × 3 mL). The organic phase was dried over sodium sulfate, filtered and concentrated *in vacuo*. The crude was purified using automated flash chromatography (Biotage Sfar Amino D 11g column, methanol/dichloromethane gradient mobile phase). Product-containing fractions were evaporated *in vacuo* to obtain a white solid (2.4 mg, 22% yield). LCMS (ESI+): 445.51. ^1^H NMR (500 MHz, DMSO-*d_6_*) δ 11.44 (s, 1H), 10.99 (s, 1H), 10.39 (s, 1H), 8.15 (s, 1H), 7.82 (d, *J* = 8.1 Hz, 1H), 7.73 (q, *J* = 8.4 Hz, 2H), 7.45 (d, *J* = 8.2 Hz, 1H), 7.23 (t, *J* = 7.6 Hz, 1H), 7.04 (t, *J* = 7.6 Hz, 1H), 5.11 (dd, *J* = 13.3, 5.1 Hz, 1H), 4.53 – 4.28 (m, 2H), 3.91 (p, *J* = 7.1 Hz, 1H), 2.95-2.88 (m, 1H), 2.66 – 2.57 (m, 1H), 2.43 – 2.34 (m, 1H), 2.07 – 1.94 (m, 1H), 1.41 (d, *J* = 7.1 Hz, 6H). ^13^C NMR (126 MHz, DMSO-*d_6_*) δ 173.40, 171.63, 168.33, 161.76, 143.74, 142.87, 136.43, 127.03, 126.94, 126.86, 126.22, 124.15, 124.10, 121.95, 119.96, 119.62, 114.34, 112.88, 52.04, 47.68, 31.71, 25.39, 23.37, 23.02.

**^1^H and ^13^C NMR spectra of Compounds 1–19.**

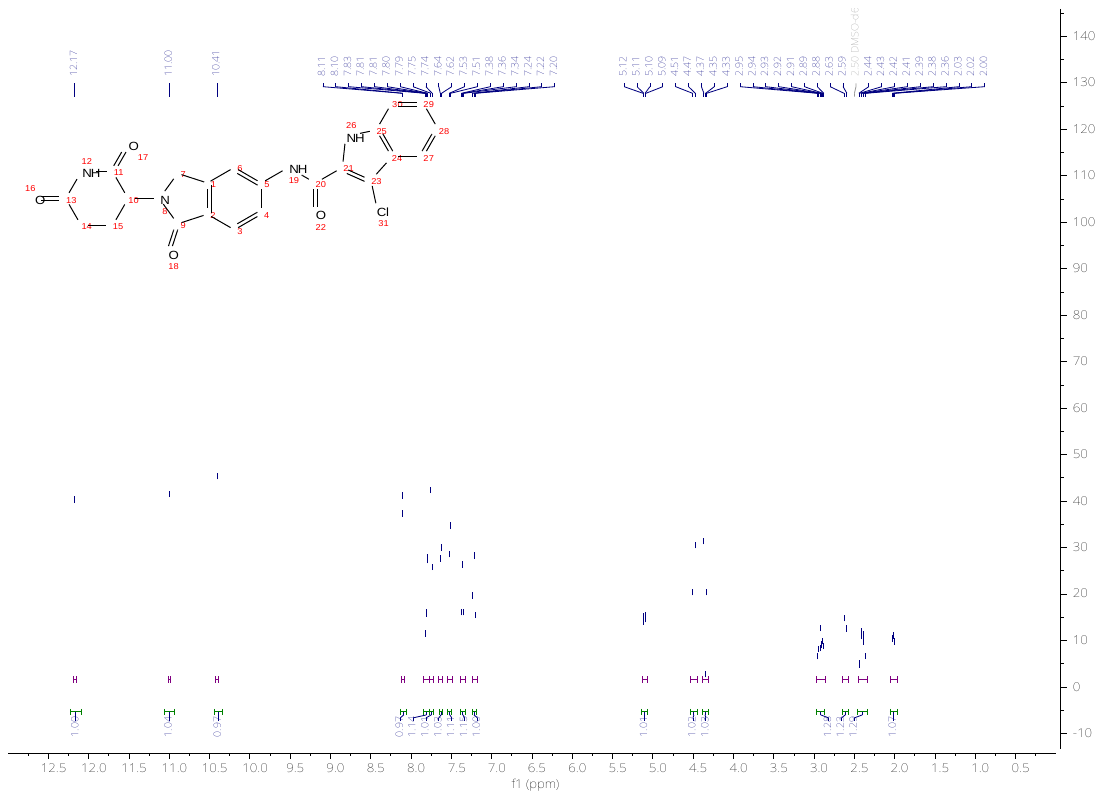

**Figure S4.** ^1^H NMR spectrum of **1** in DMSO-*d_6_*.

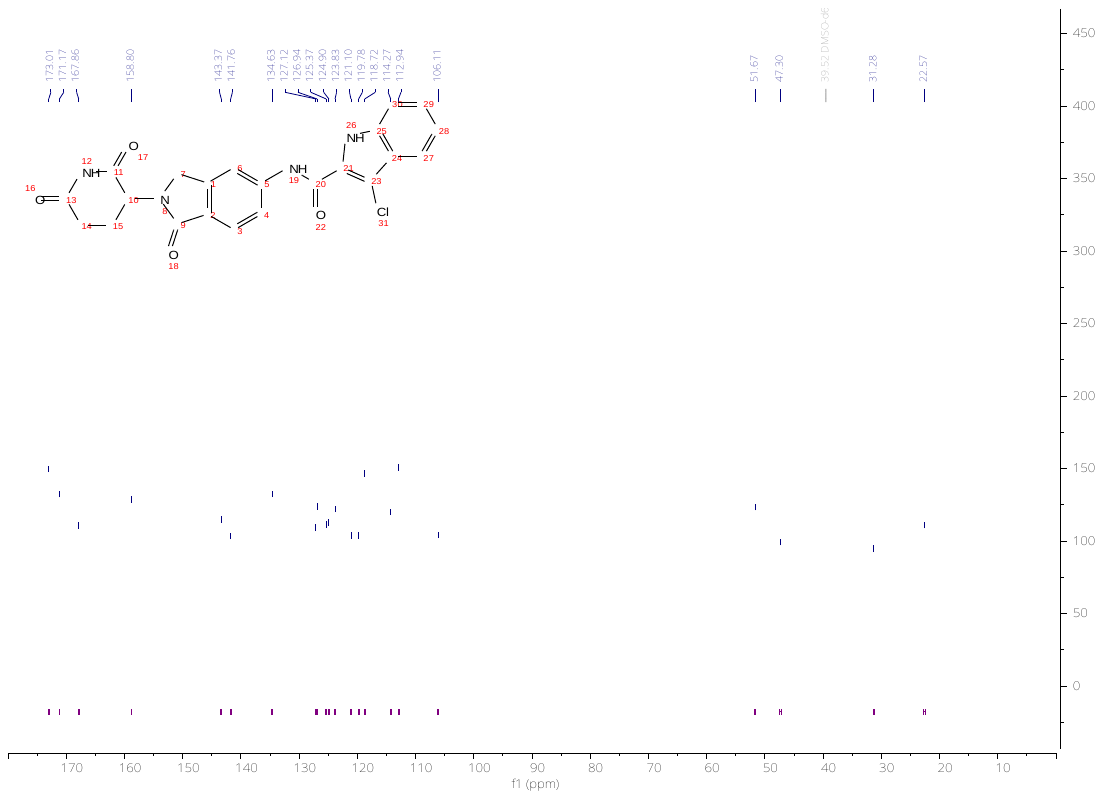

**Figure S5.** ^13^C NMR spectrum of **1** in DMSO-*d_6_*.

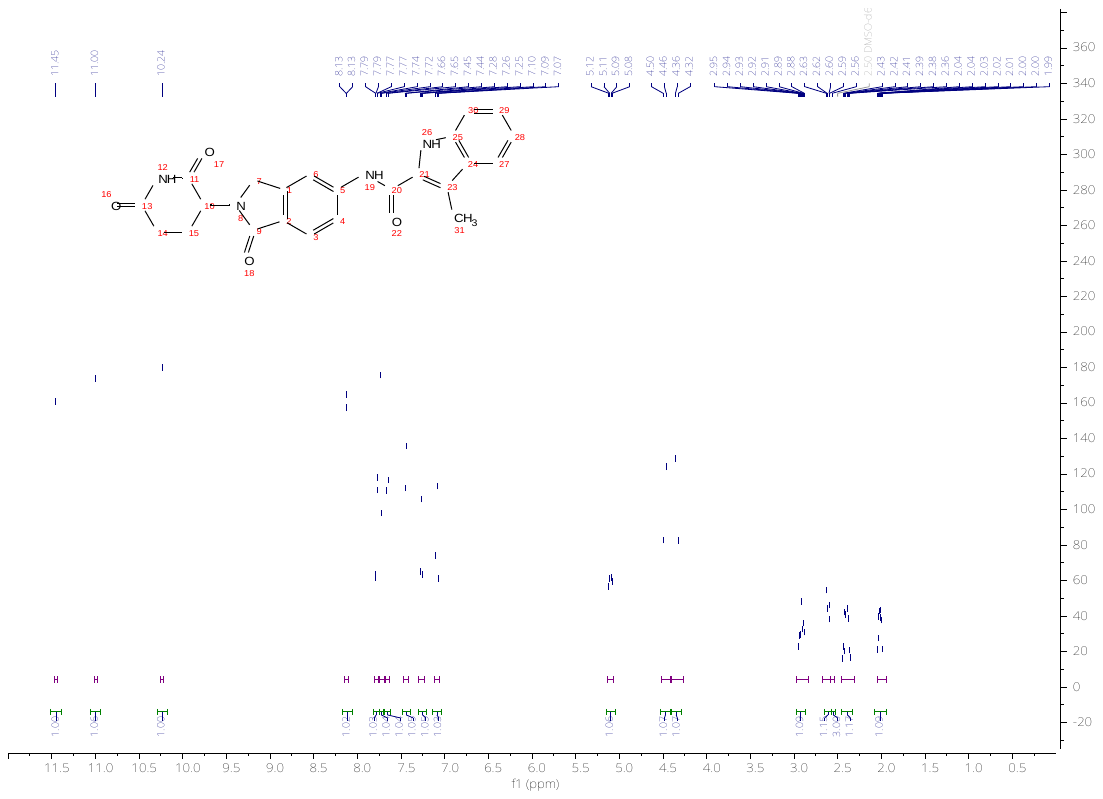

**Figure S6.** ^1^H NMR spectrum of **2** in DMSO-*d_6_*.

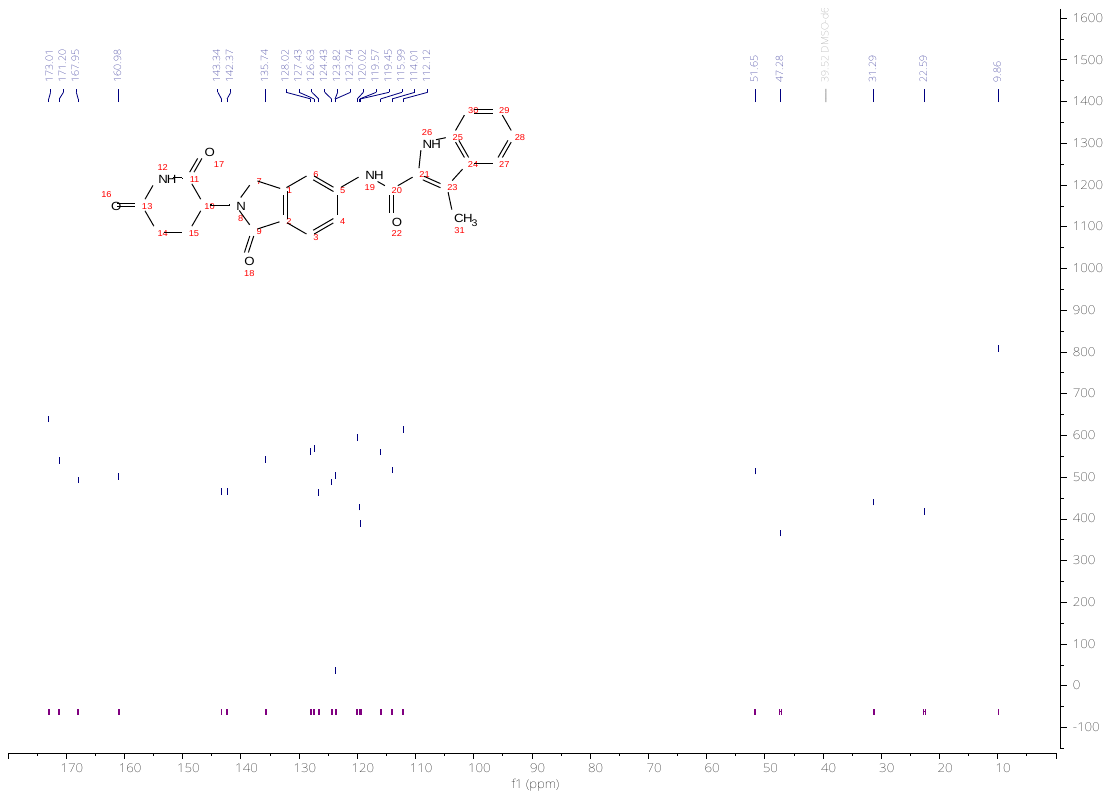

**Figure S7.** ^13^C NMR spectrum of **2** in DMSO-*d_6_*.

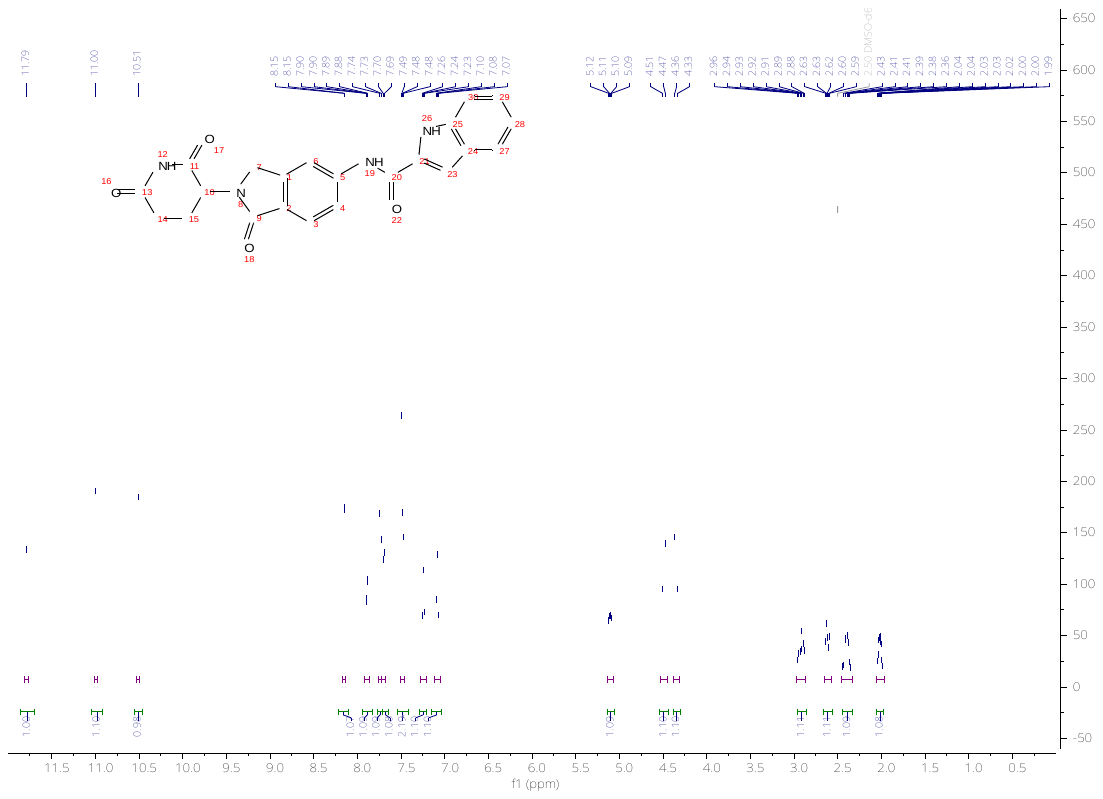

**Figure S8.** ^1^H NMR spectrum of **3** in DMSO-*d_6_*.

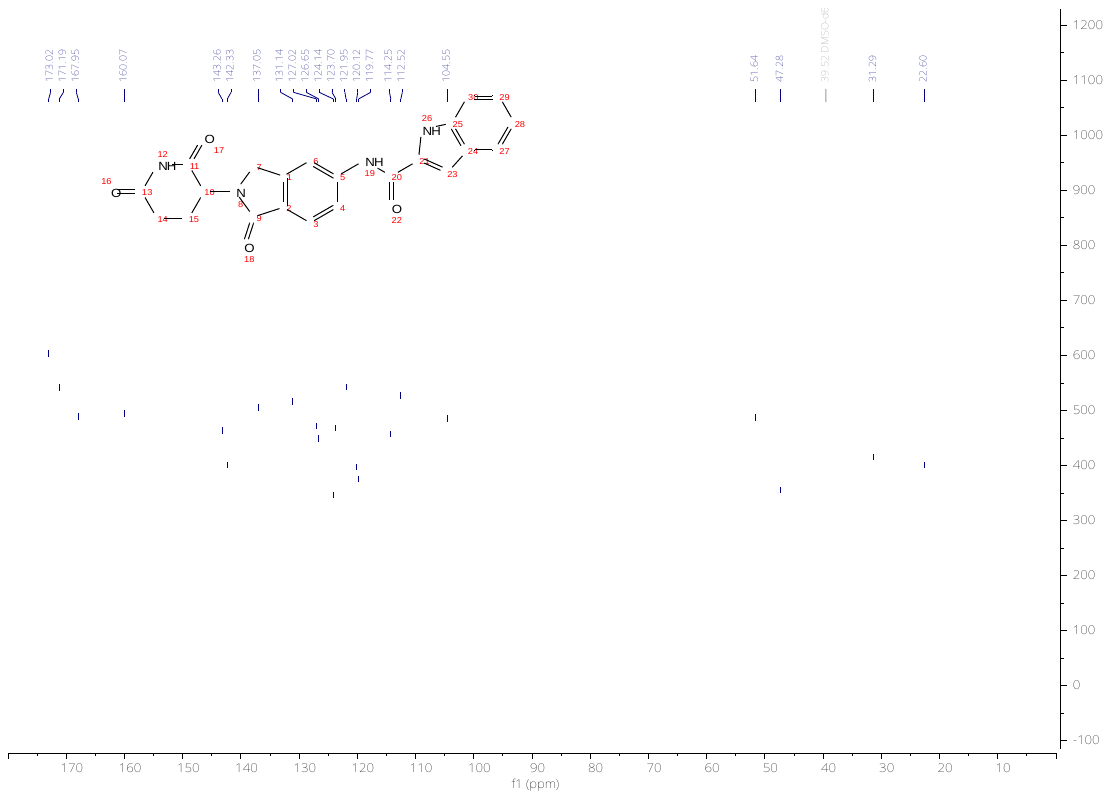

**Figure S9.** ^13^C NMR spectrum of **3** in DMSO-*d_6_*.

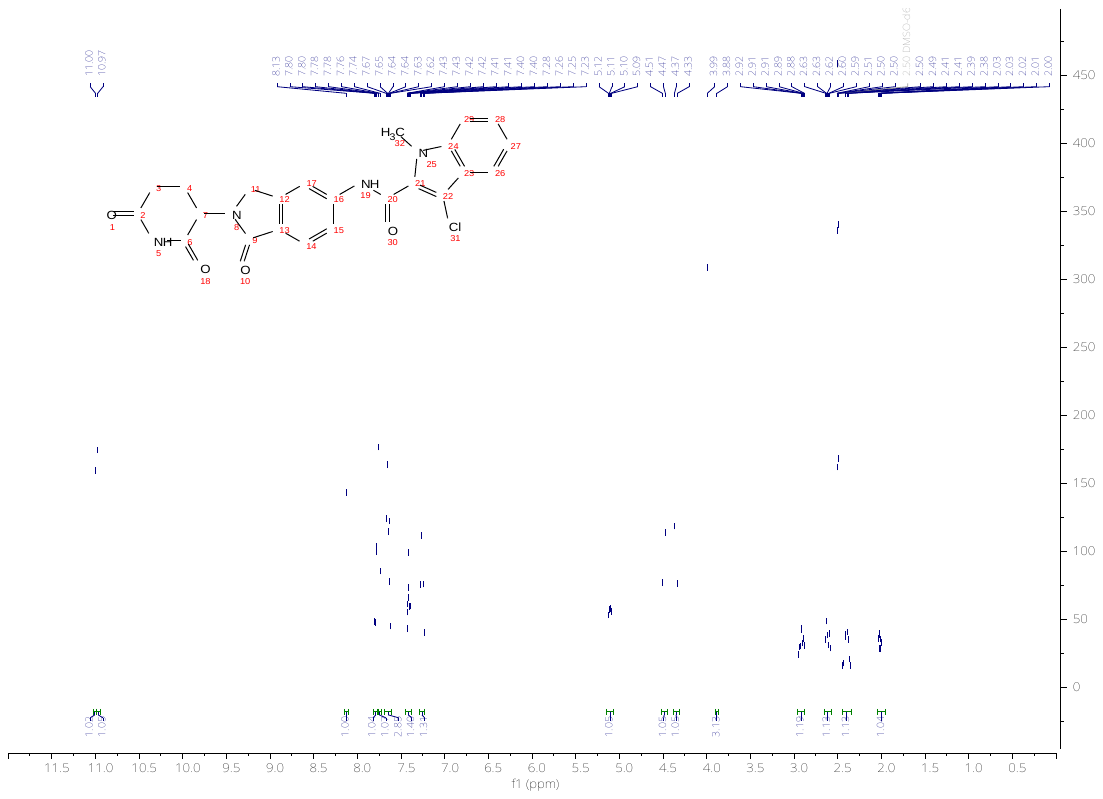

**Figure S10.** ^1^H NMR spectrum of **4** in DMSO-*d_6_*.

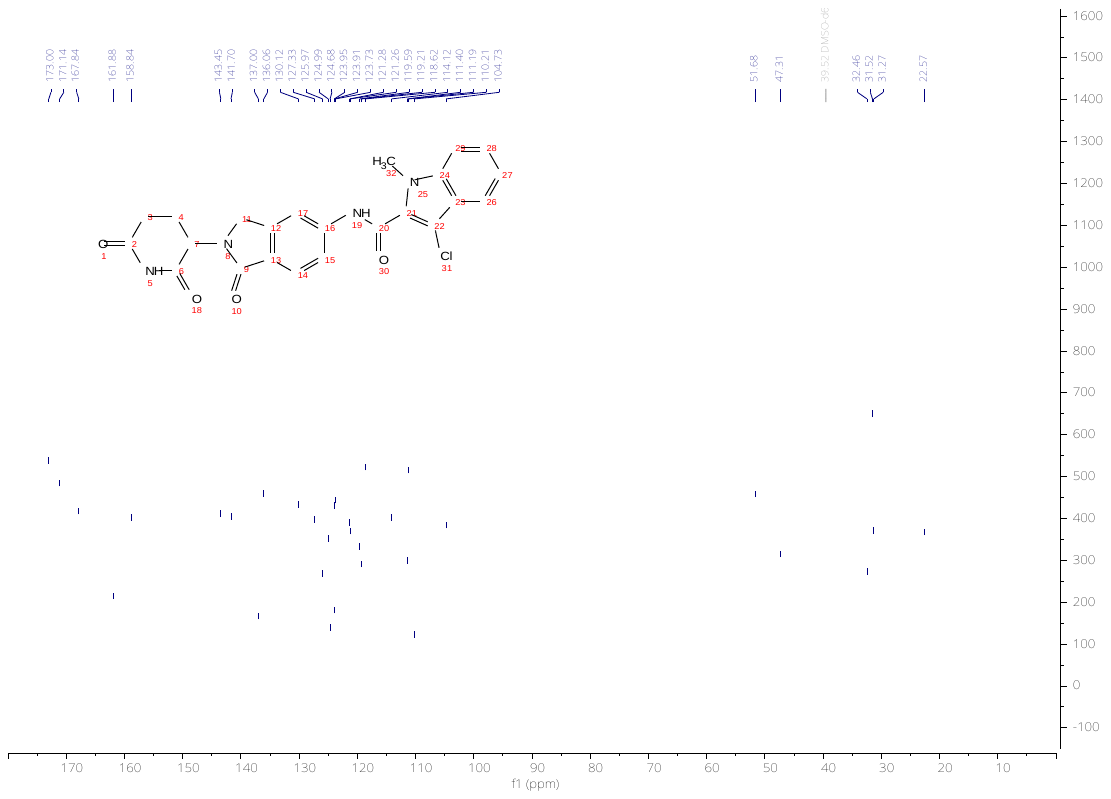

**Figure S11.** ^13^C NMR spectrum of **4** in DMSO-*d_6_*.

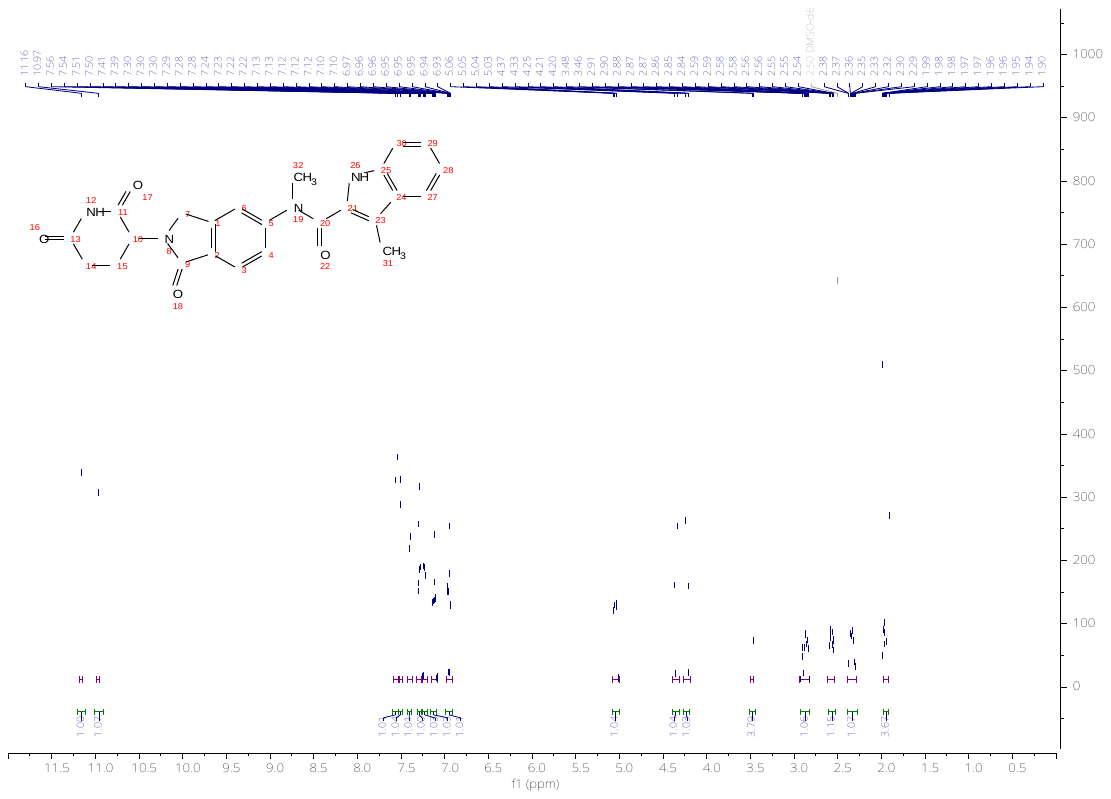

**Figure S12.** ^1^H NMR spectrum of **5** in DMSO-*d_6_*.

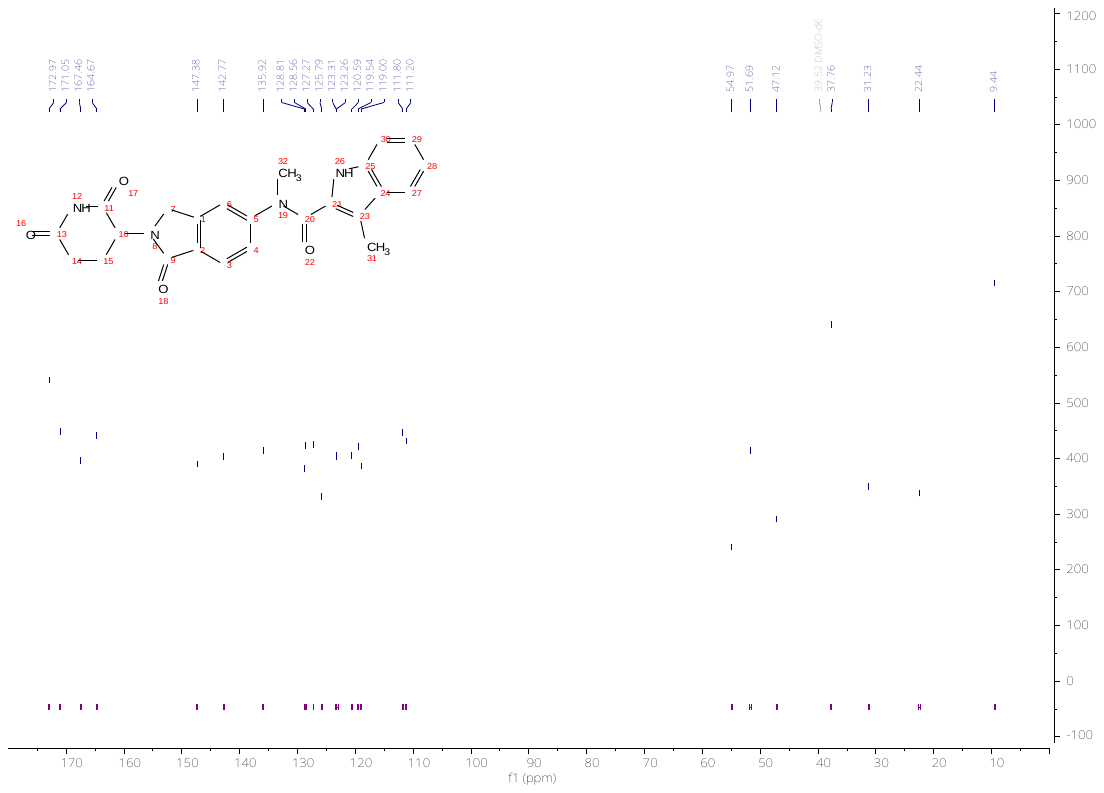

**Figure S13.** ^13^C NMR spectrum of **5** in DMSO-*d_6_*.

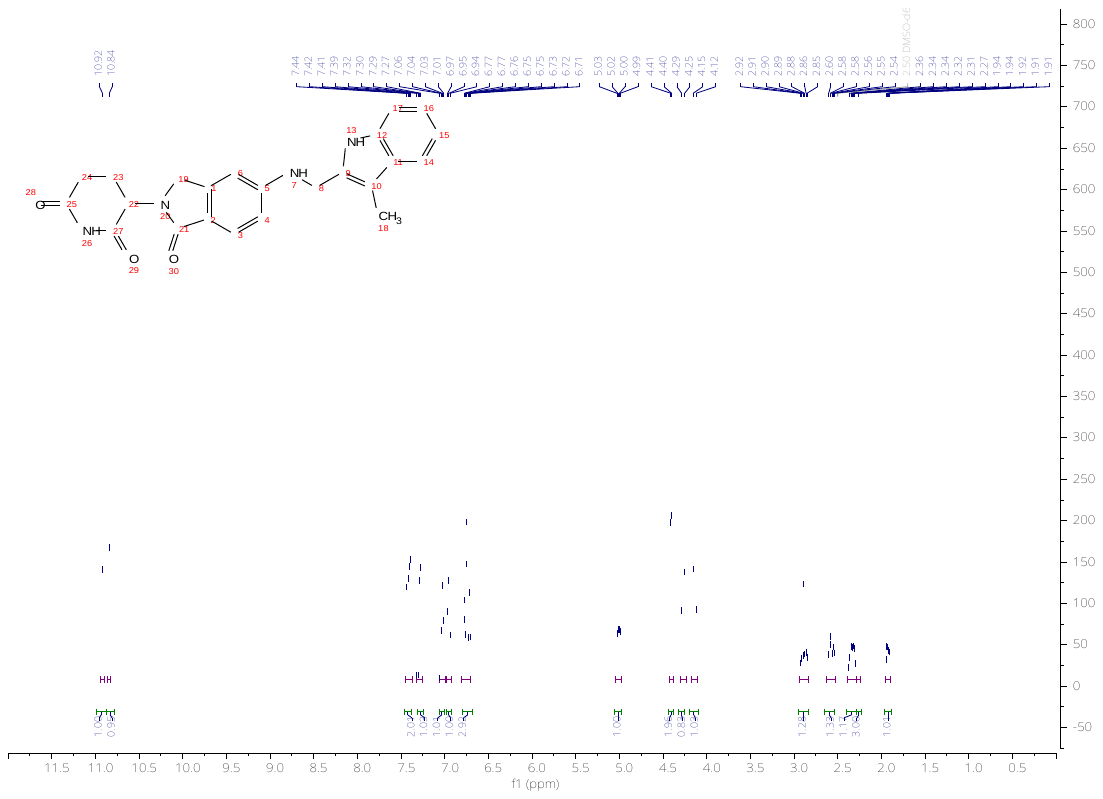

**Figure S14.** ^1^H NMR spectrum of **6** in DMSO-*d_6_*.

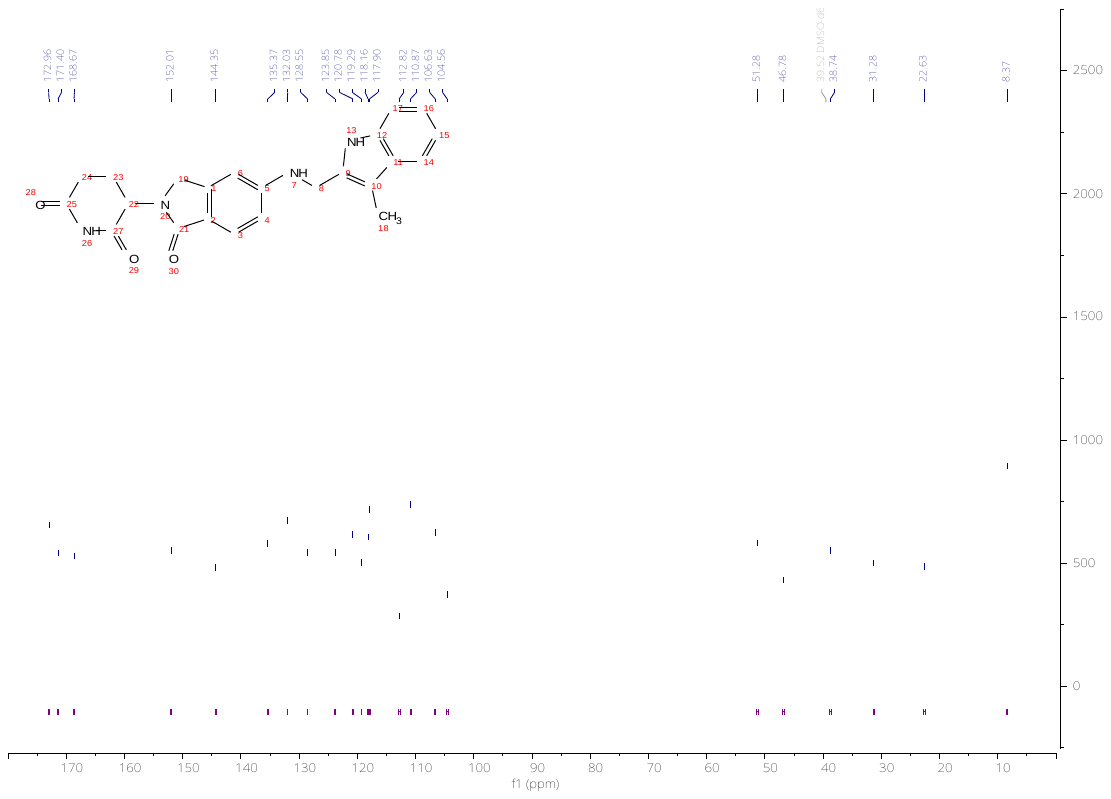

**Figure S15.** ^13^C NMR spectrum of **6** in DMSO-*d_6_*.

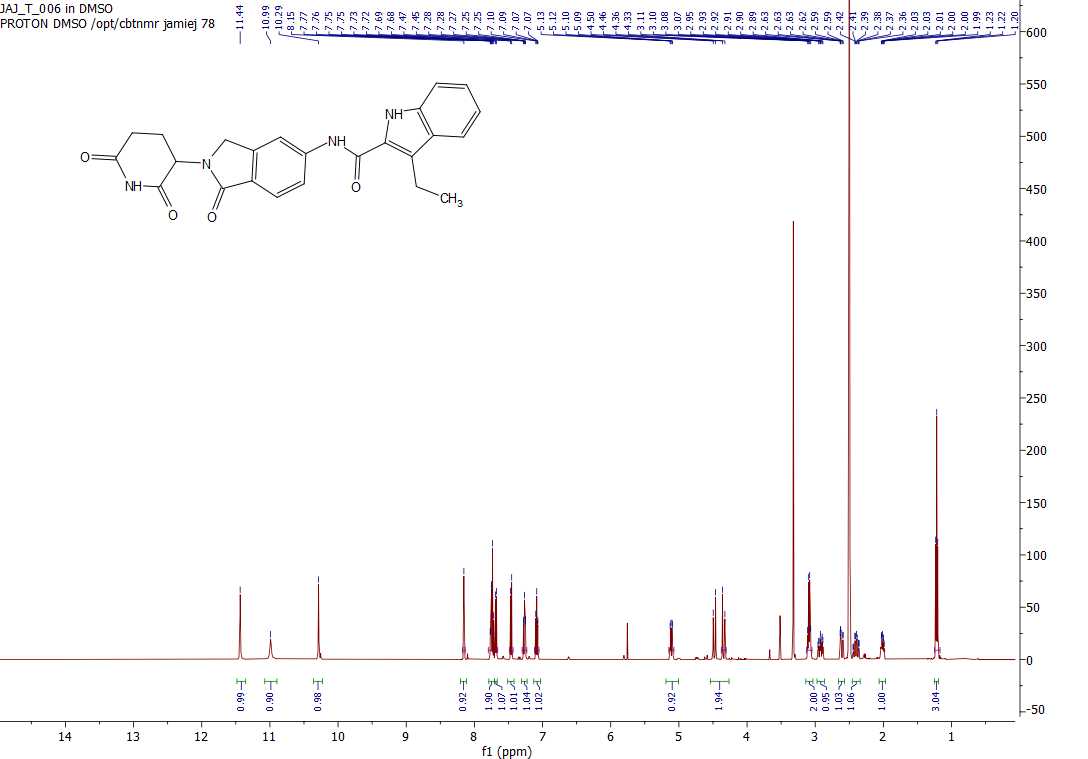

**Figure S16.** ^1^H NMR spectrum of **7** in DMSO-*d_6_*.

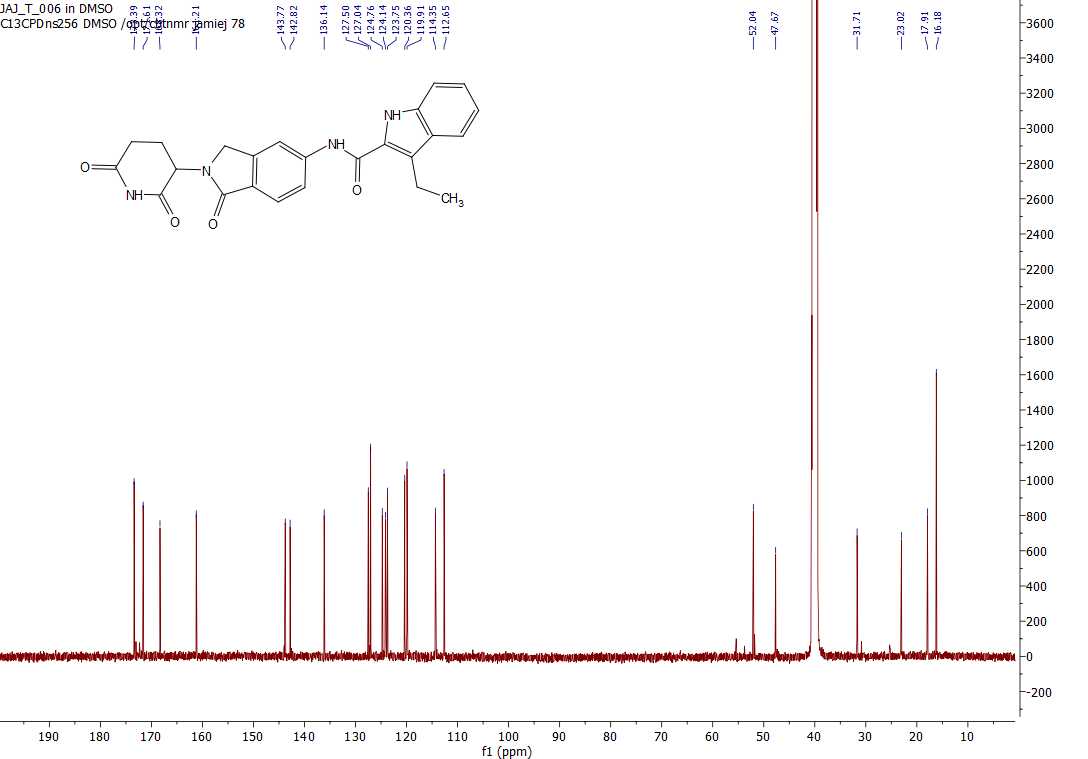

**Figure S17.** ^13^C NMR spectrum of **7** in DMSO-*d_6_*.

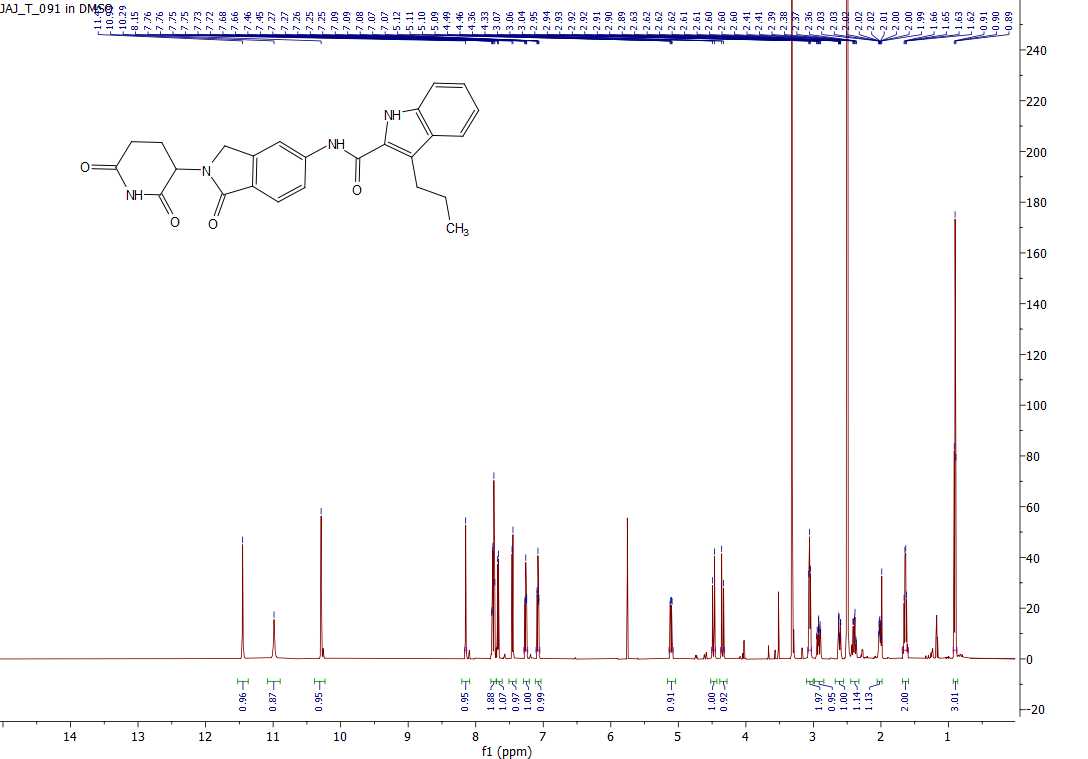

**Figure S18.** ^1^H NMR spectrum of **8** in DMSO-*d_6_*.

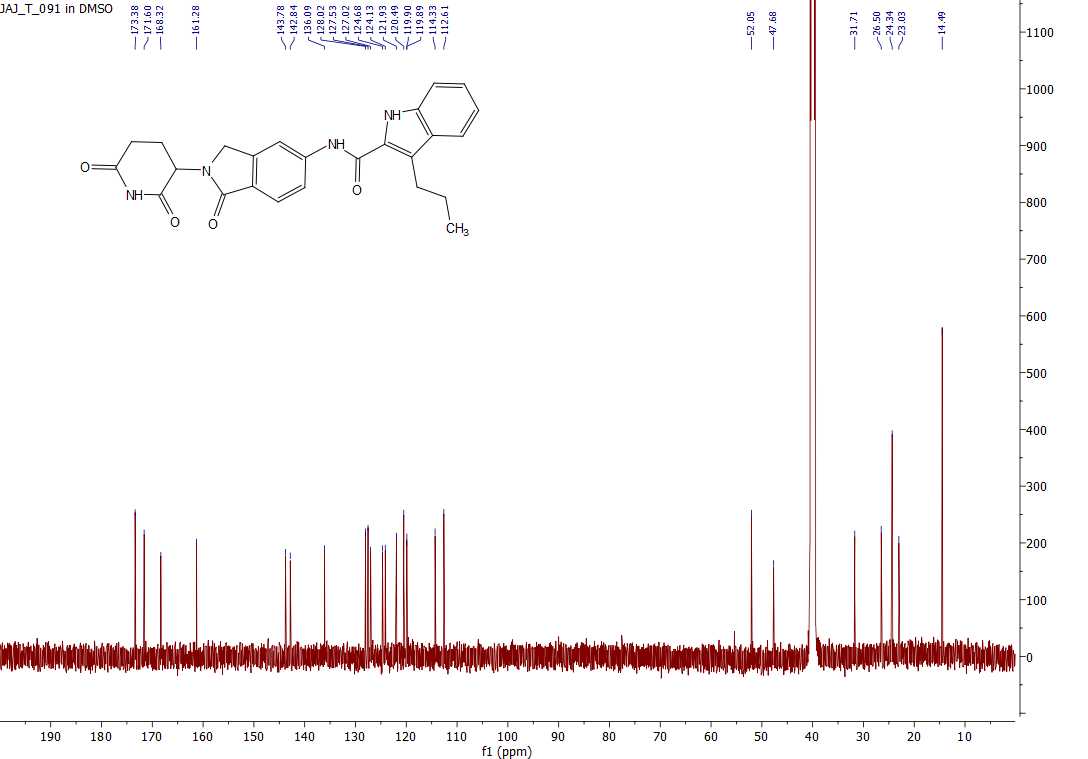

**Figure S19.** ^13^C NMR spectrum of **8** in DMSO-*d_6_*.

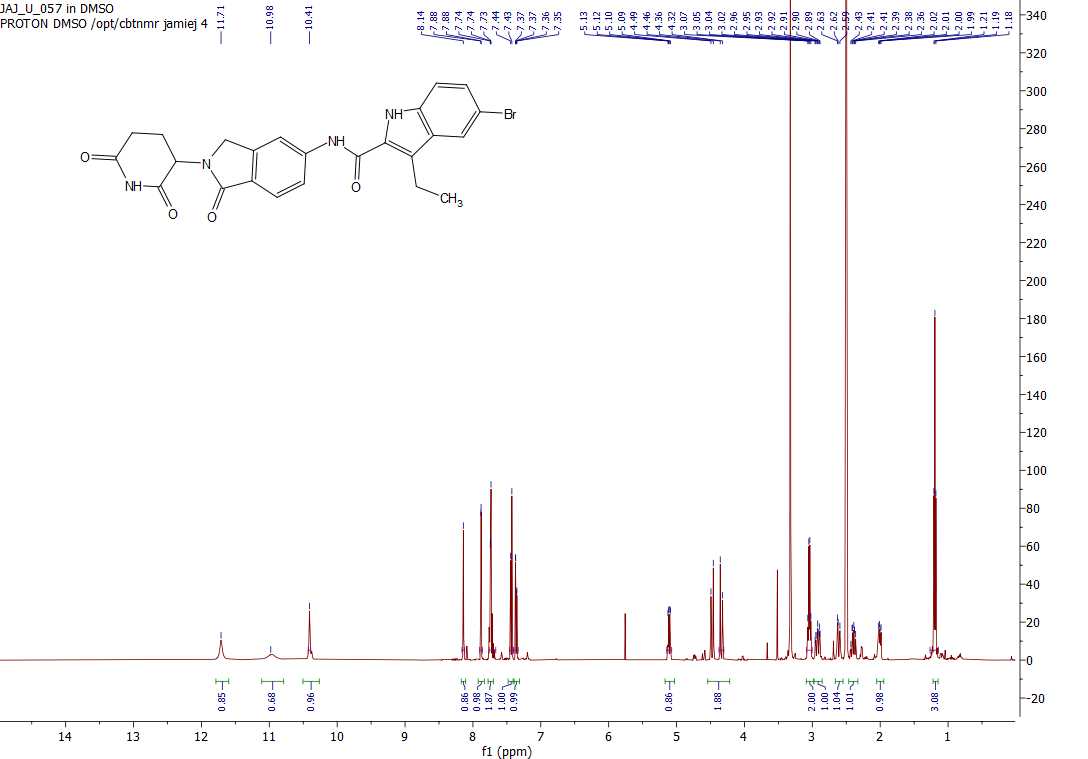

**Figure S20.** ^1^H NMR spectrum of **9** in DMSO-*d_6_*.

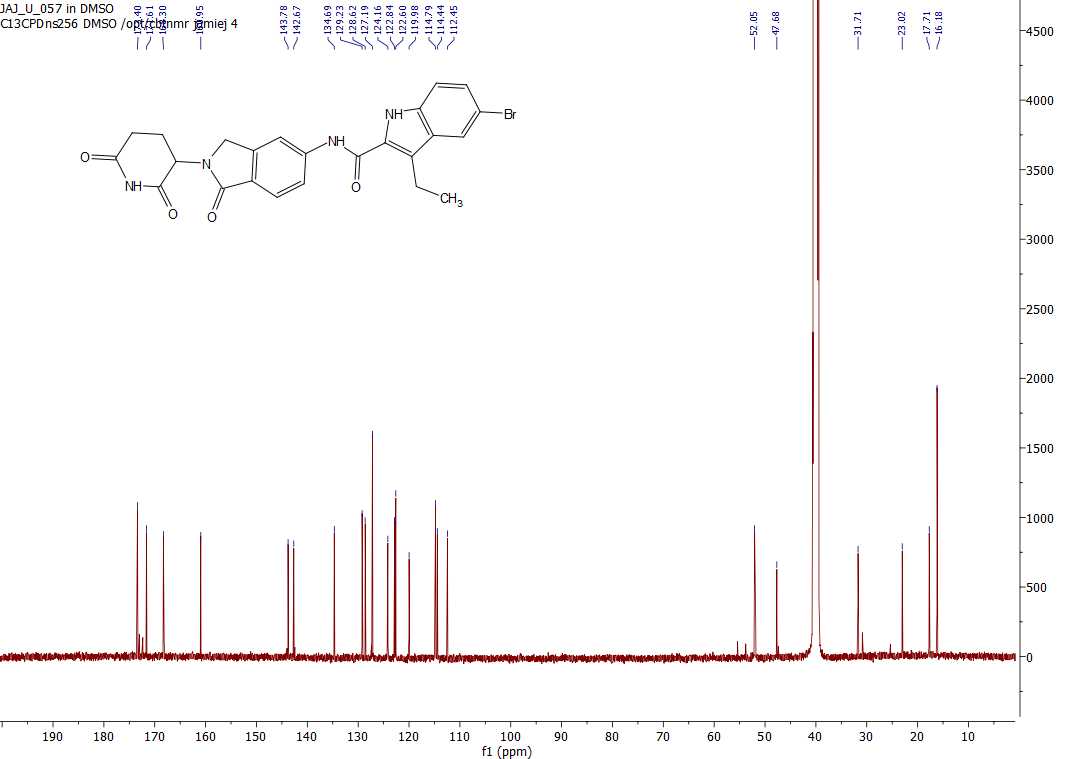

**Figure S21.** ^13^C NMR spectrum of **9** in DMSO-*d_6_*.

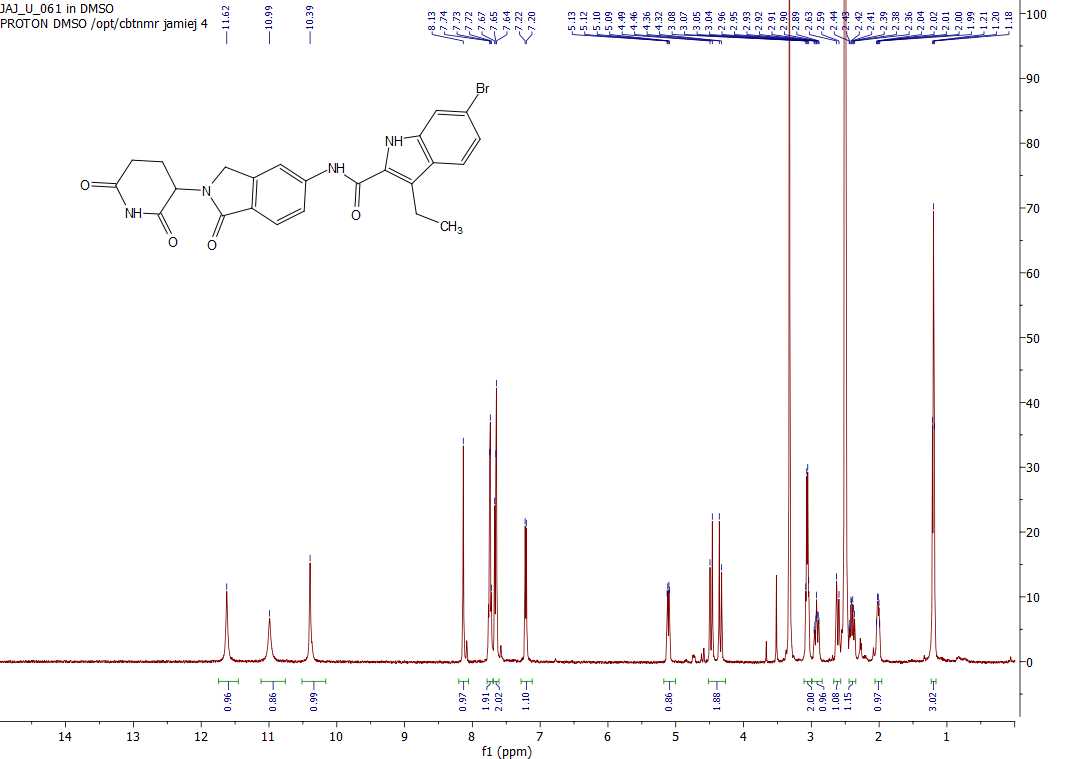

**Figure S22.** ^1^H NMR spectrum of **10** in DMSO-*d_6_*.

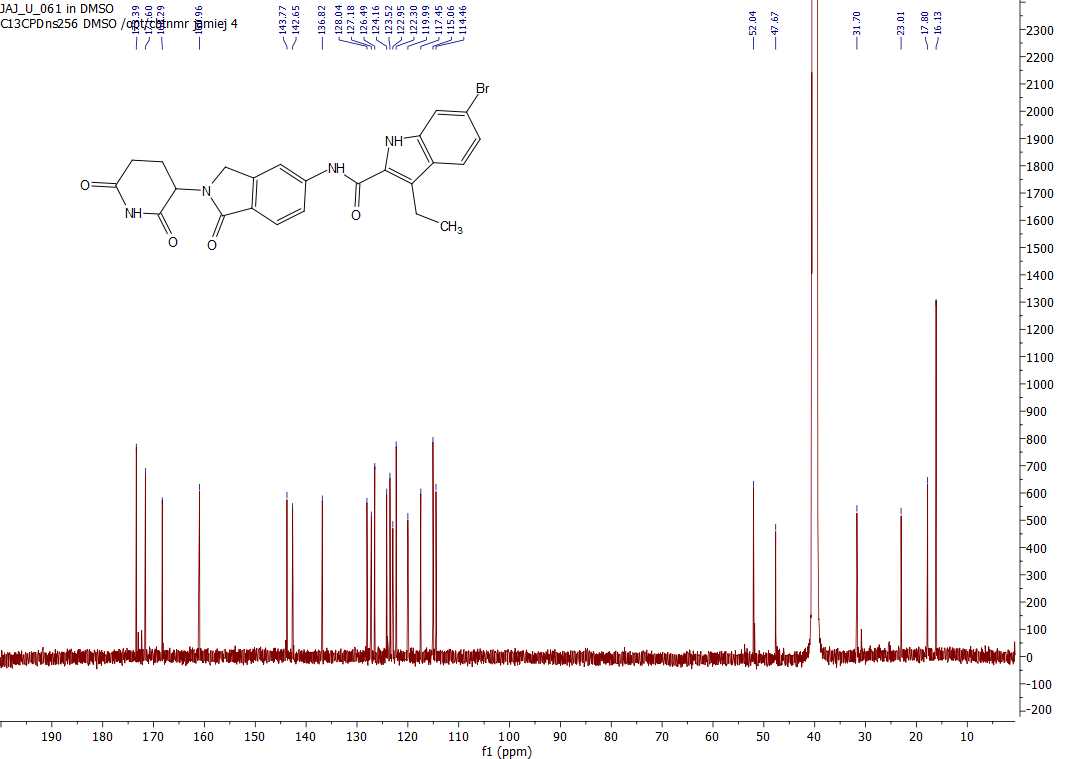

**Figure S23.** ^13^C NMR spectrum of **10** in DMSO-*d_6_*.

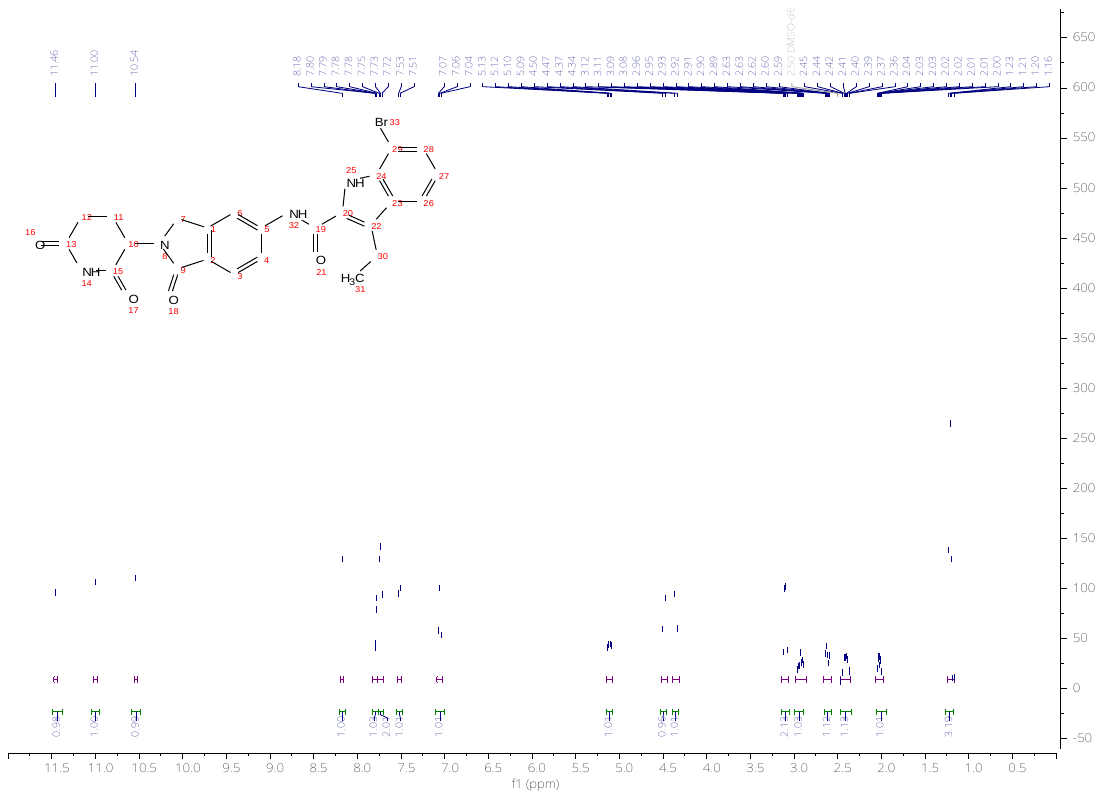

**Figure S24.** ^1^H NMR spectrum of **11** in DMSO-*d_6_*.

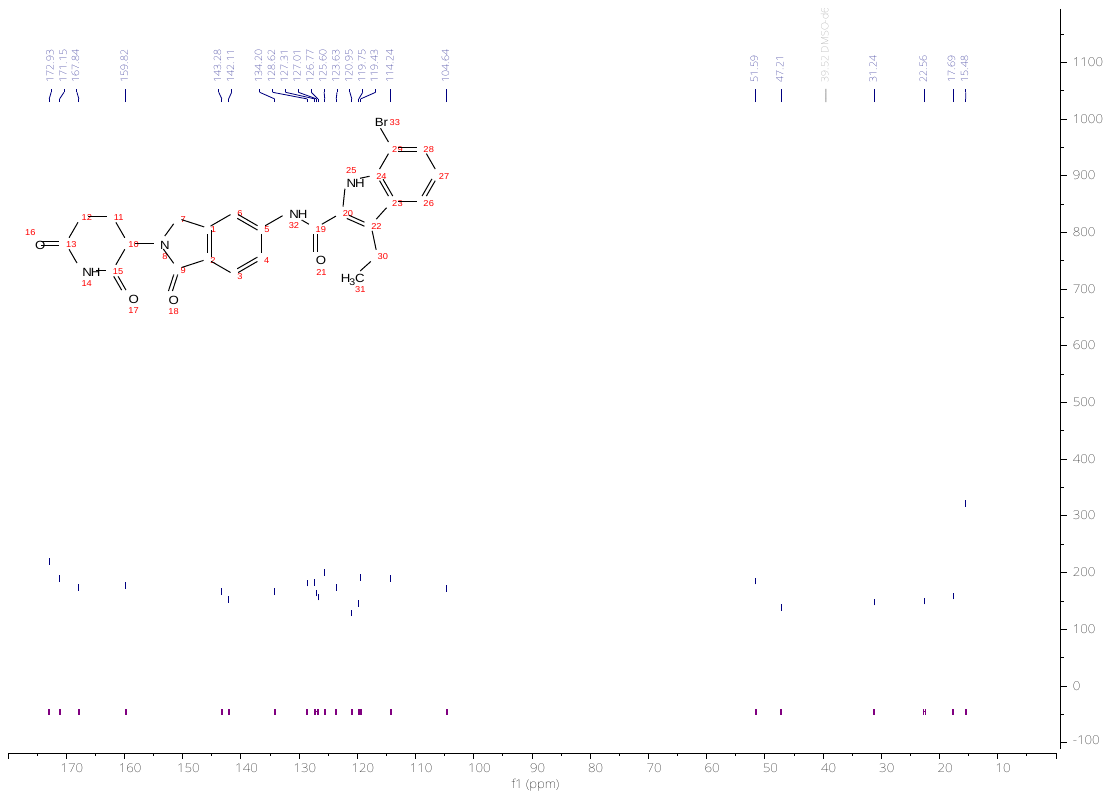

**Figure S25.** ^13^C NMR spectrum of **11** in DMSO-*d_6_*.

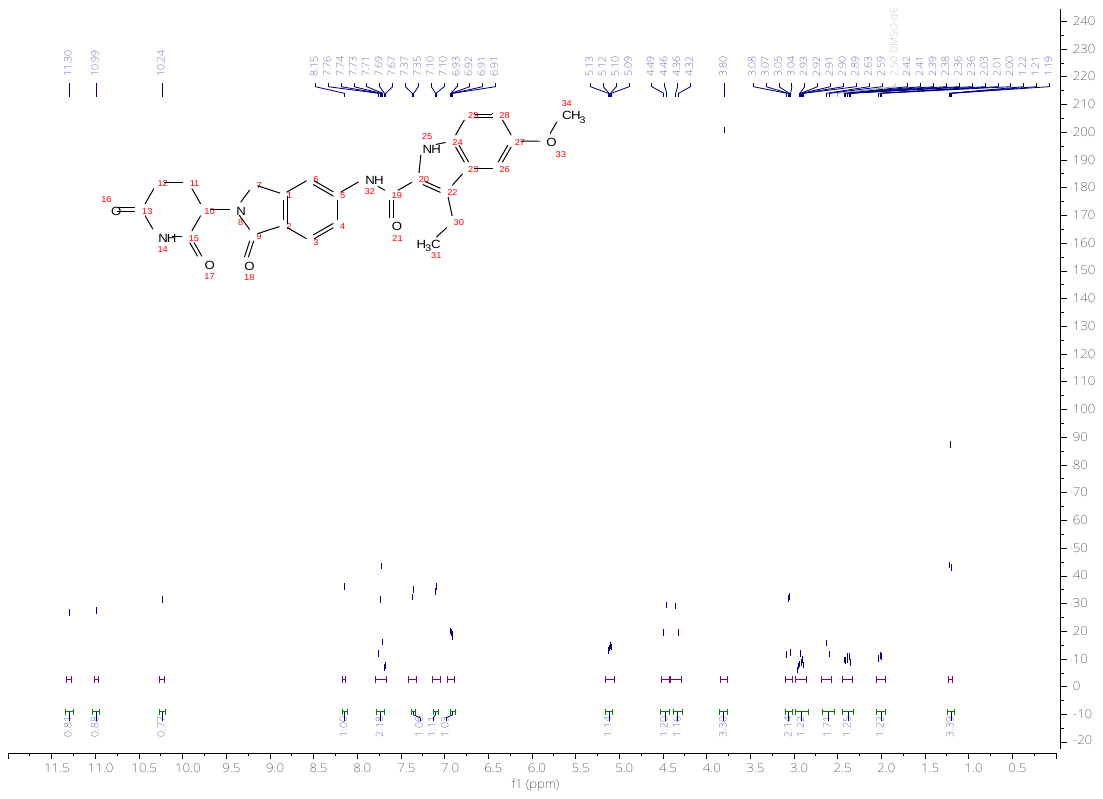

**Figure S26.** ^1^H NMR spectrum of **12** in DMSO-*d_6_*.

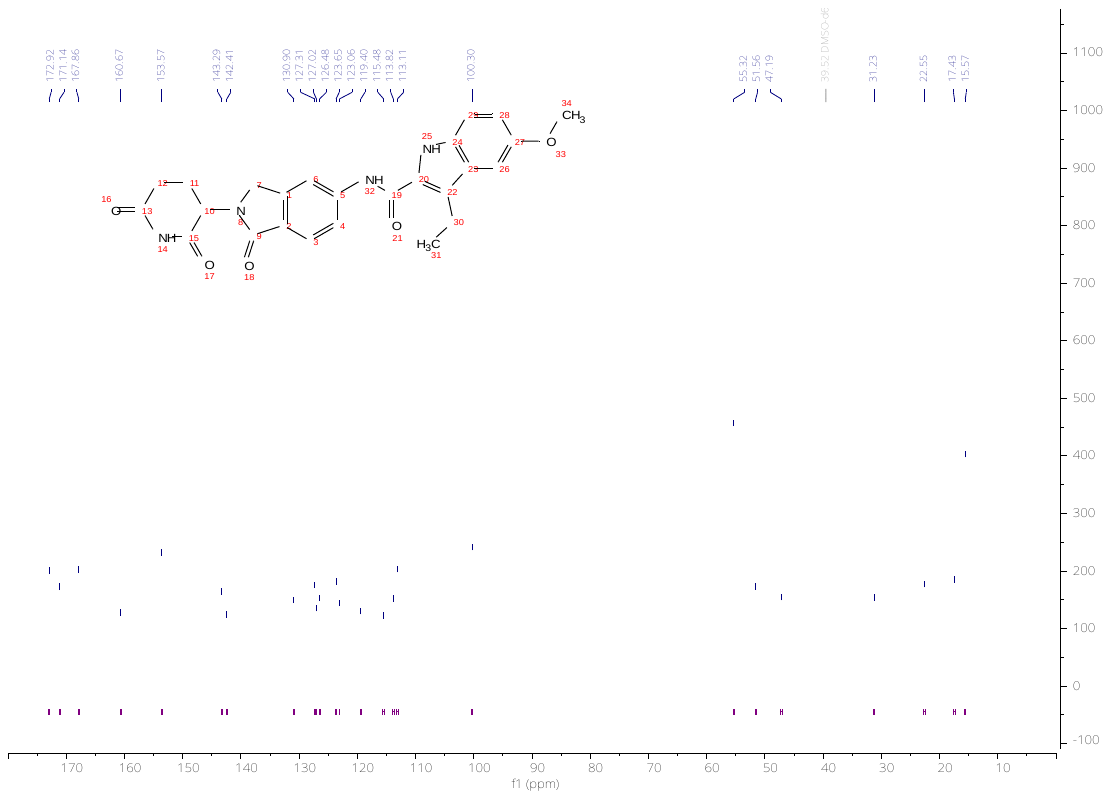

**Figure S27.** ^13^C NMR spectrum of **12** in DMSO-*d_6_*.

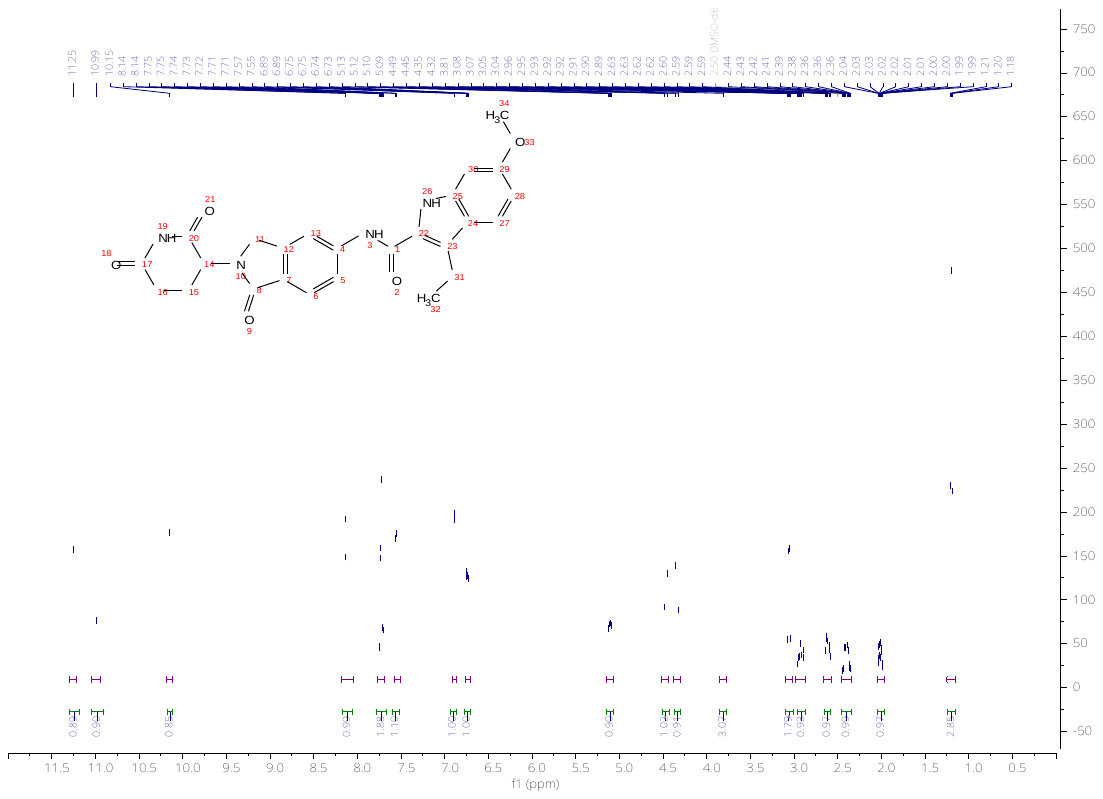

**Figure S28.** ^1^H NMR spectrum of **13** in DMSO-*d_6_*.

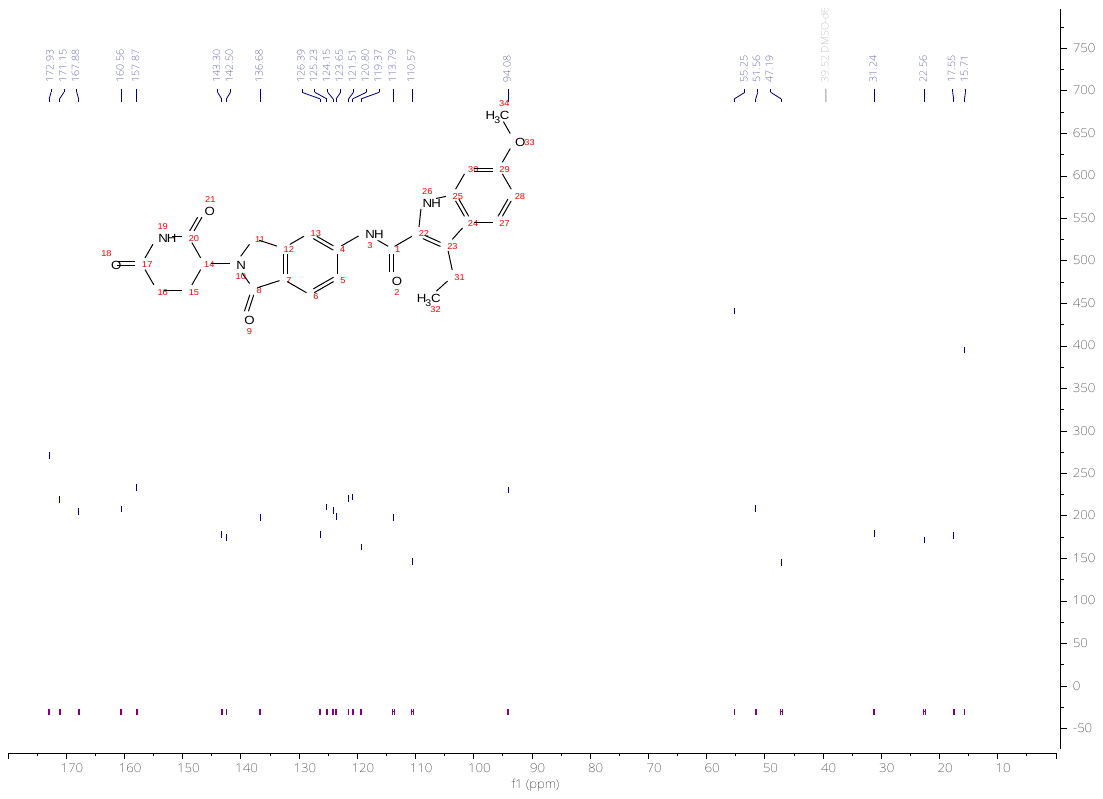

**Figure S29.** ^13^C NMR spectrum of **13** in DMSO-*d_6_*.

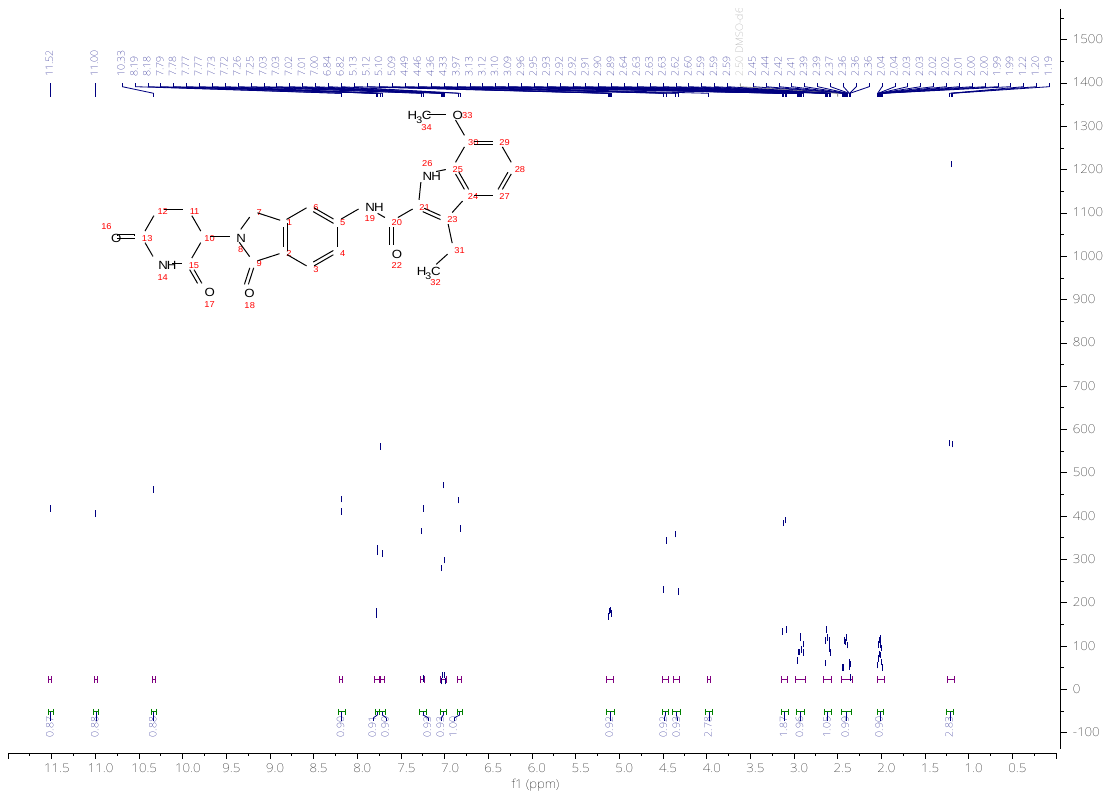

**Figure S30.** ^1^H NMR spectrum of **14** in DMSO-*d_6_*.

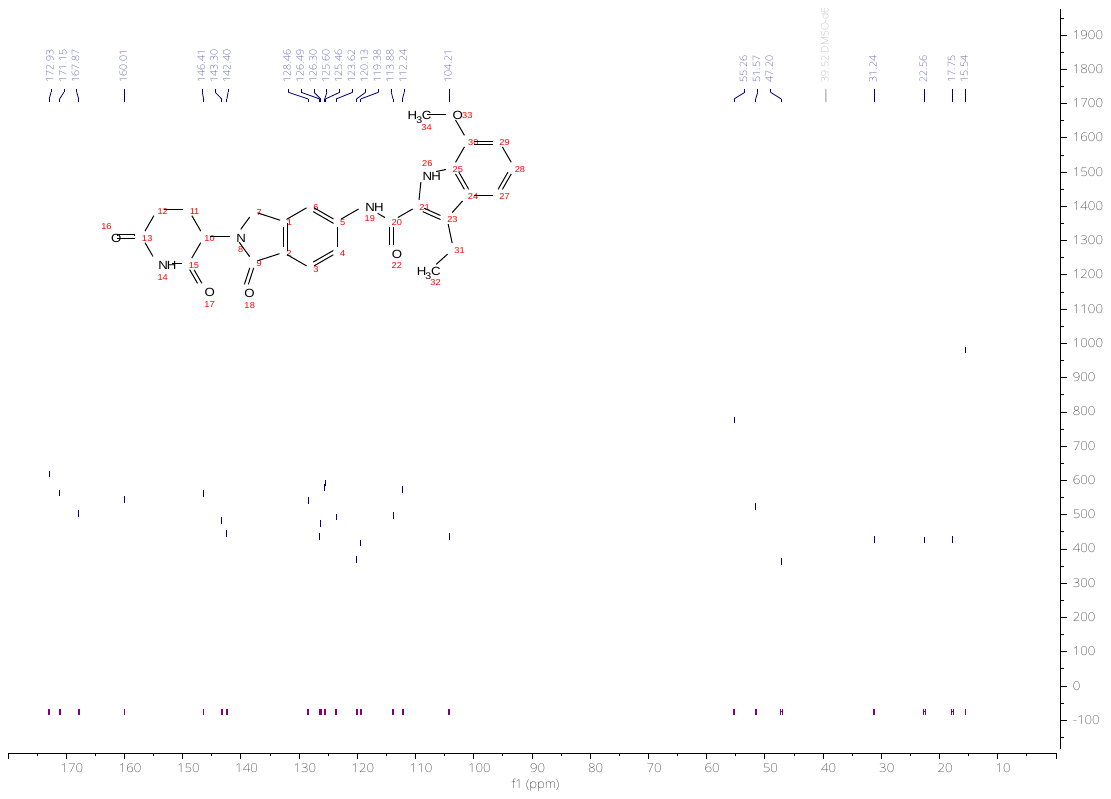

**Figure S31.** ^13^C NMR spectrum of **14** in DMSO-*d_6_*.

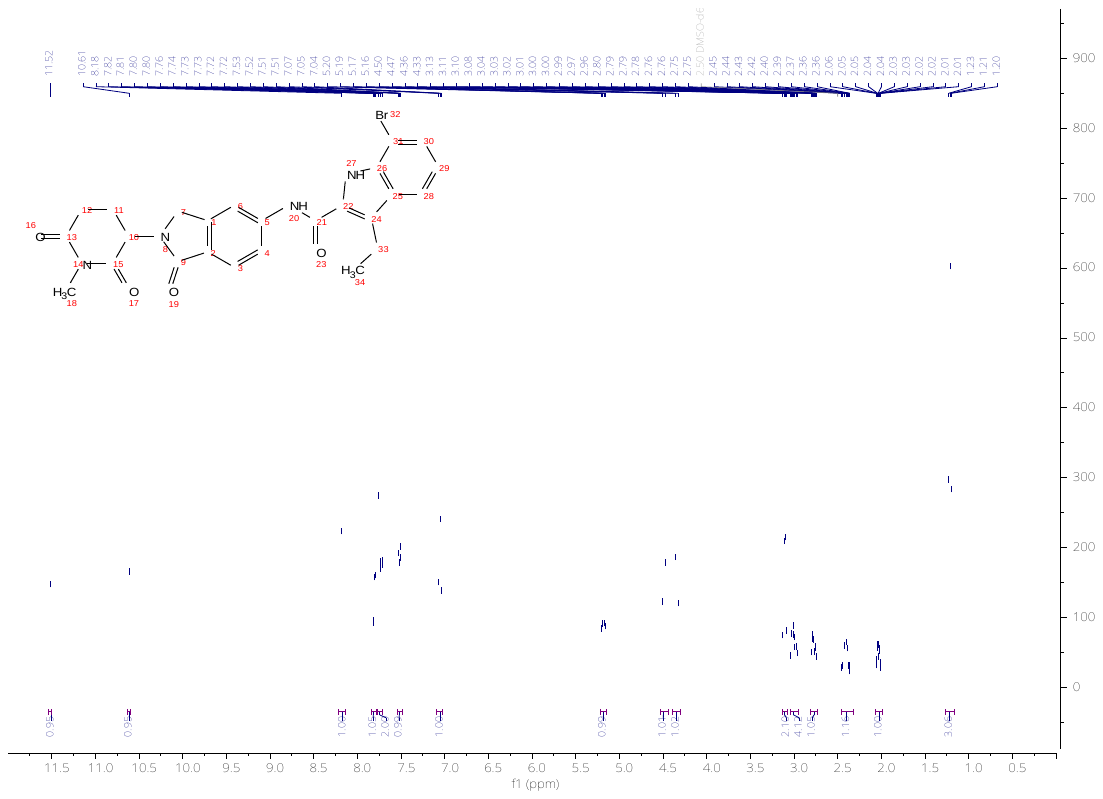

**Figure S32.** ^1^H NMR spectrum of **15** in DMSO-*d_6_*.

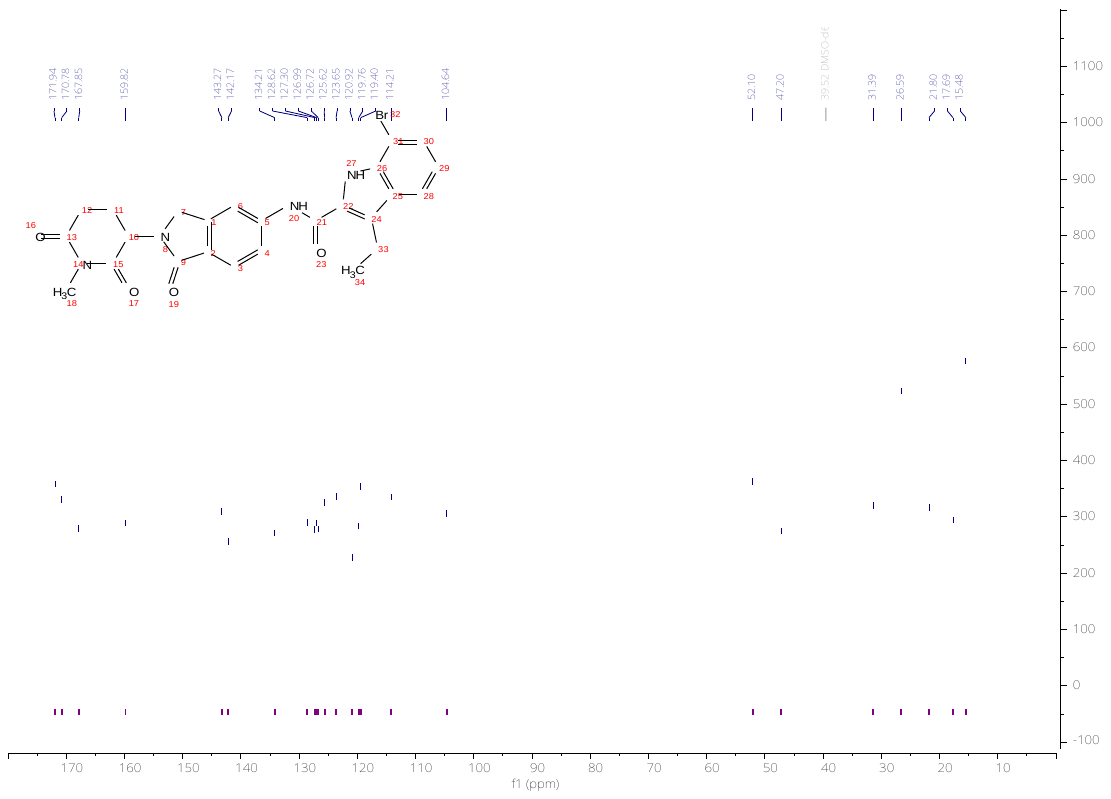

**Figure S33.** ^13^C NMR spectrum of **15** in DMSO-*d_6_*.

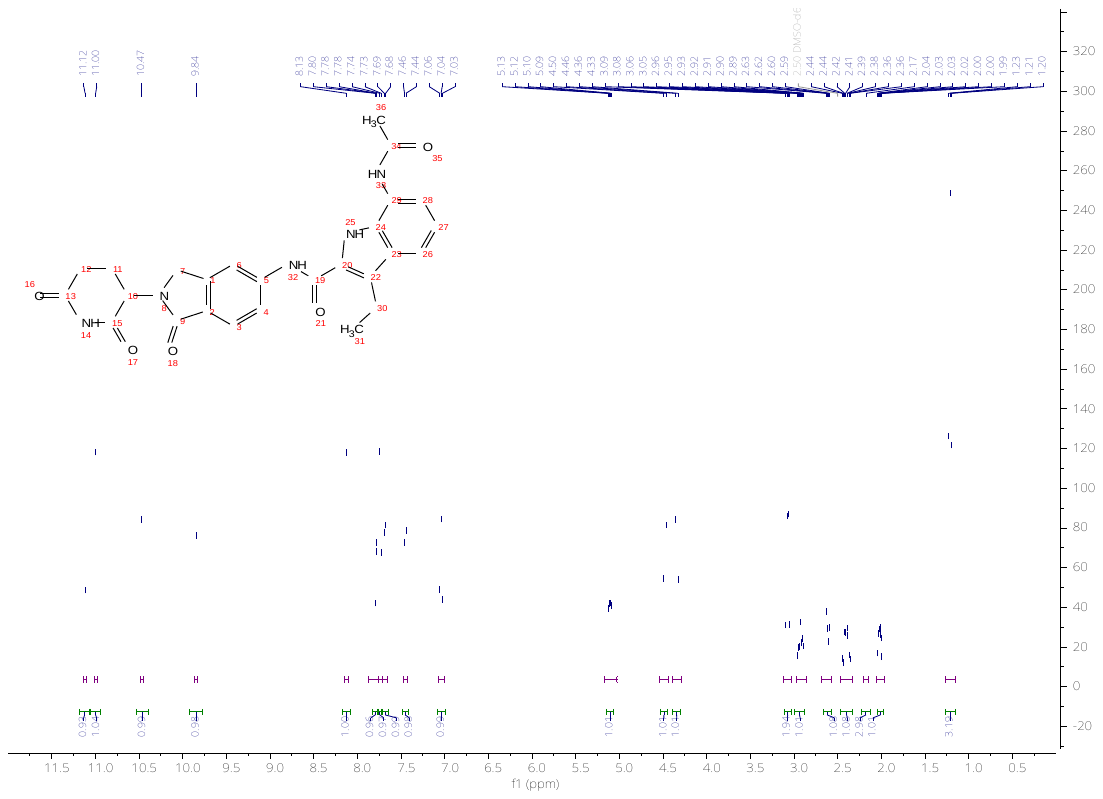

**Figure S34.** ^1^H NMR spectrum of **16** in DMSO-*d_6_*.

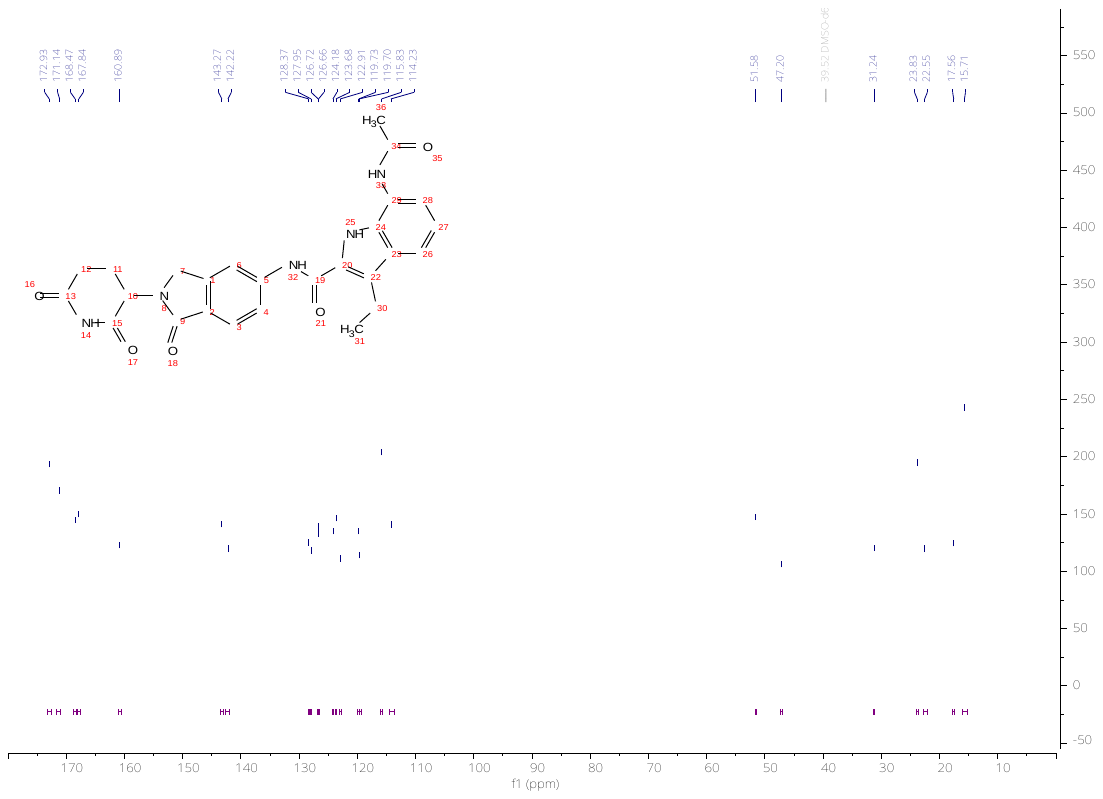

**Figure S35.** ^13^C NMR spectrum of **16** in DMSO-*d_6_*.

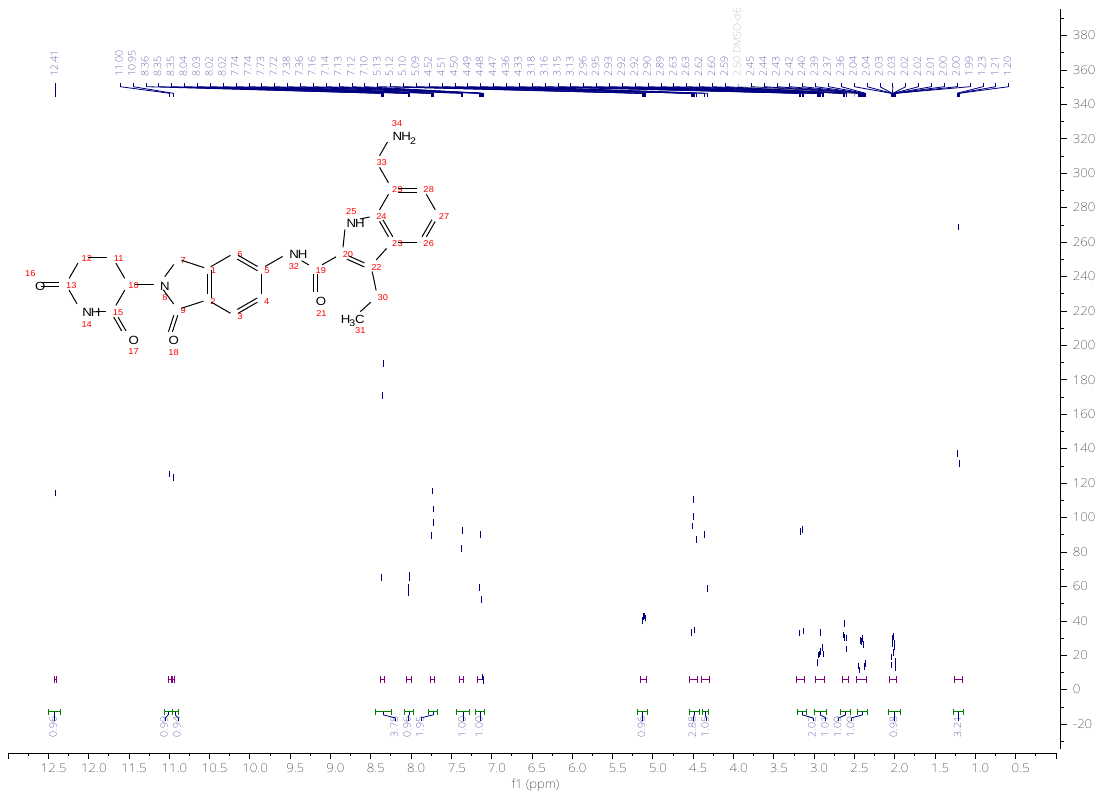

**Figure S36.** ^1^H NMR spectrum of **17** in DMSO-*d_6_*.

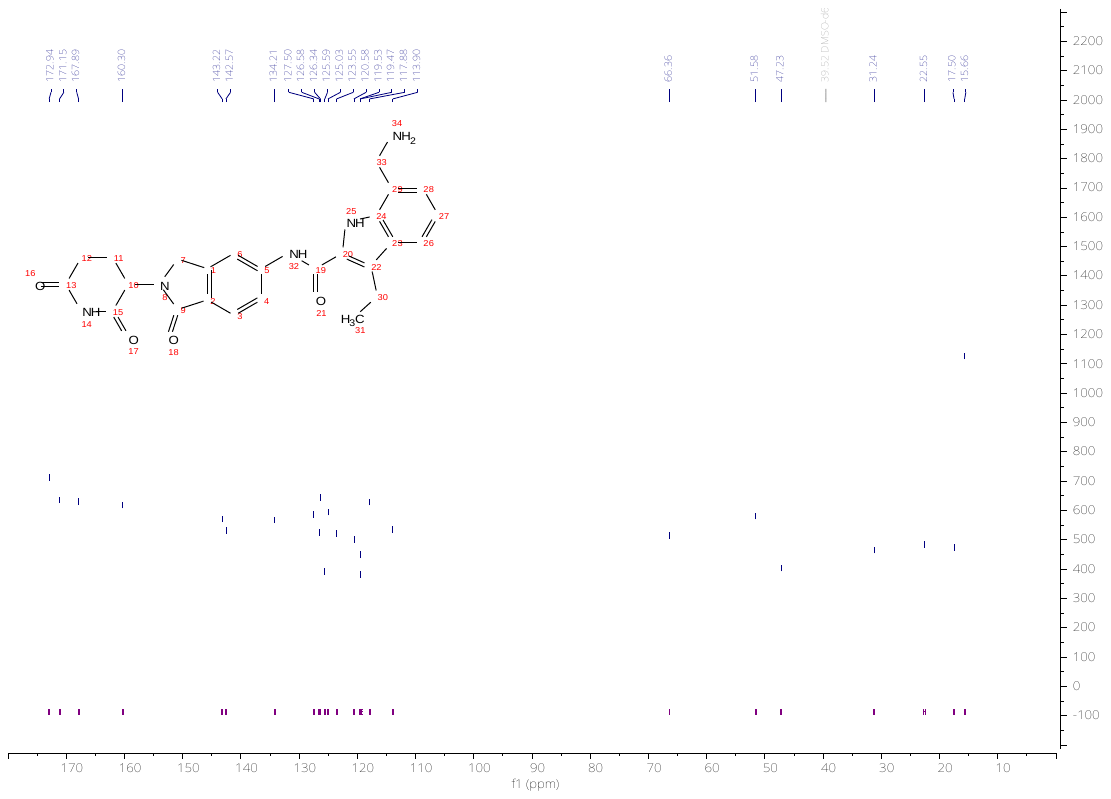

**Figure S37.** ^13^C NMR spectrum of **17** in DMSO-*d_6_*.

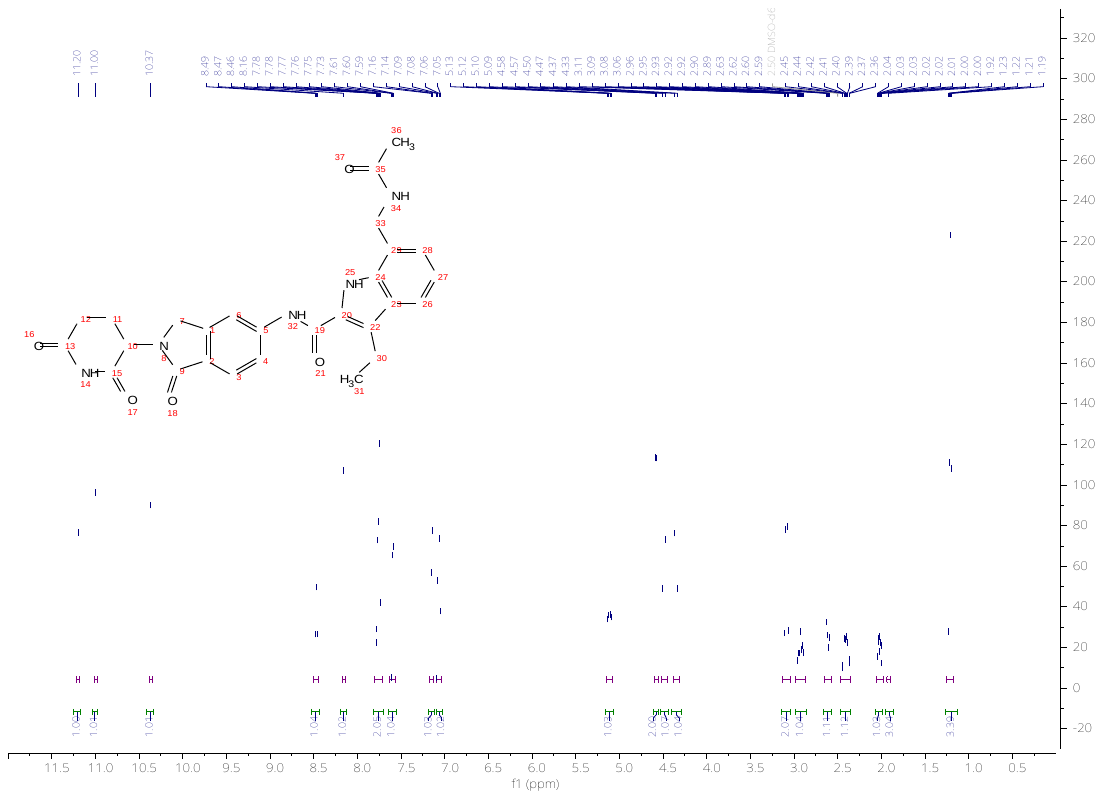

**Figure S38.** ^1^H NMR spectrum of **18** in DMSO-*d_6_*.

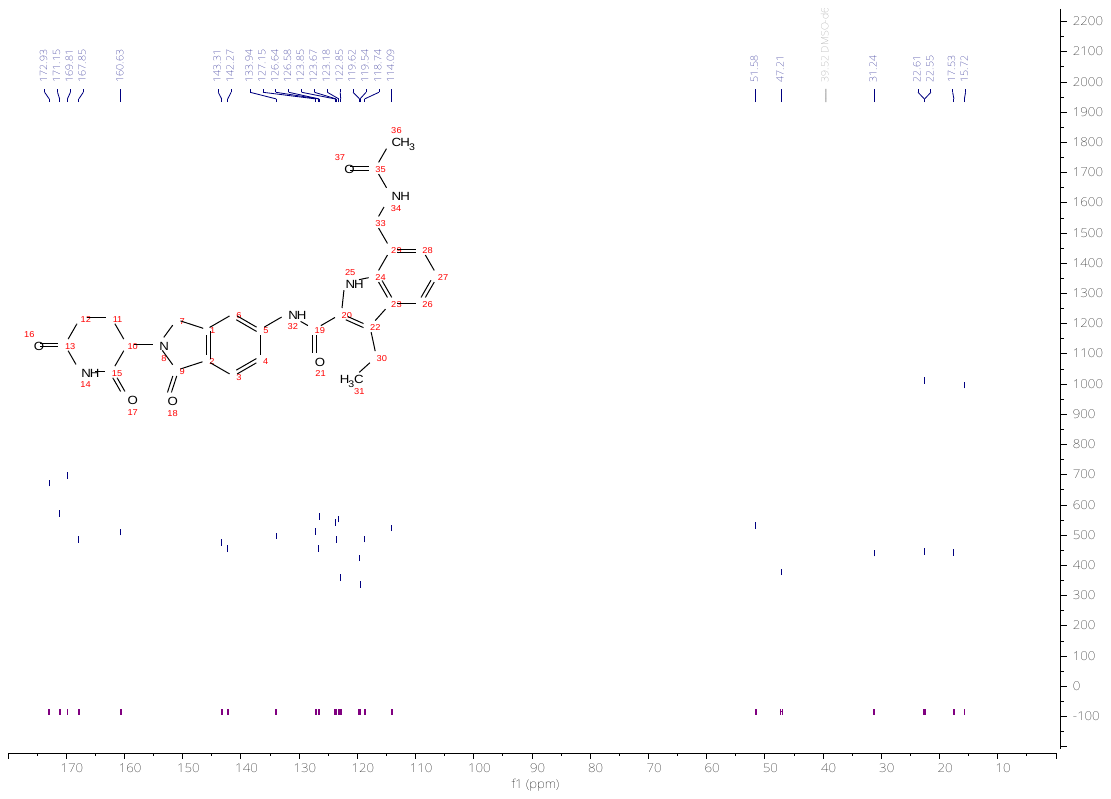

**Figure S39.** ^13^C NMR spectrum of **18** in DMSO-*d_6_*.

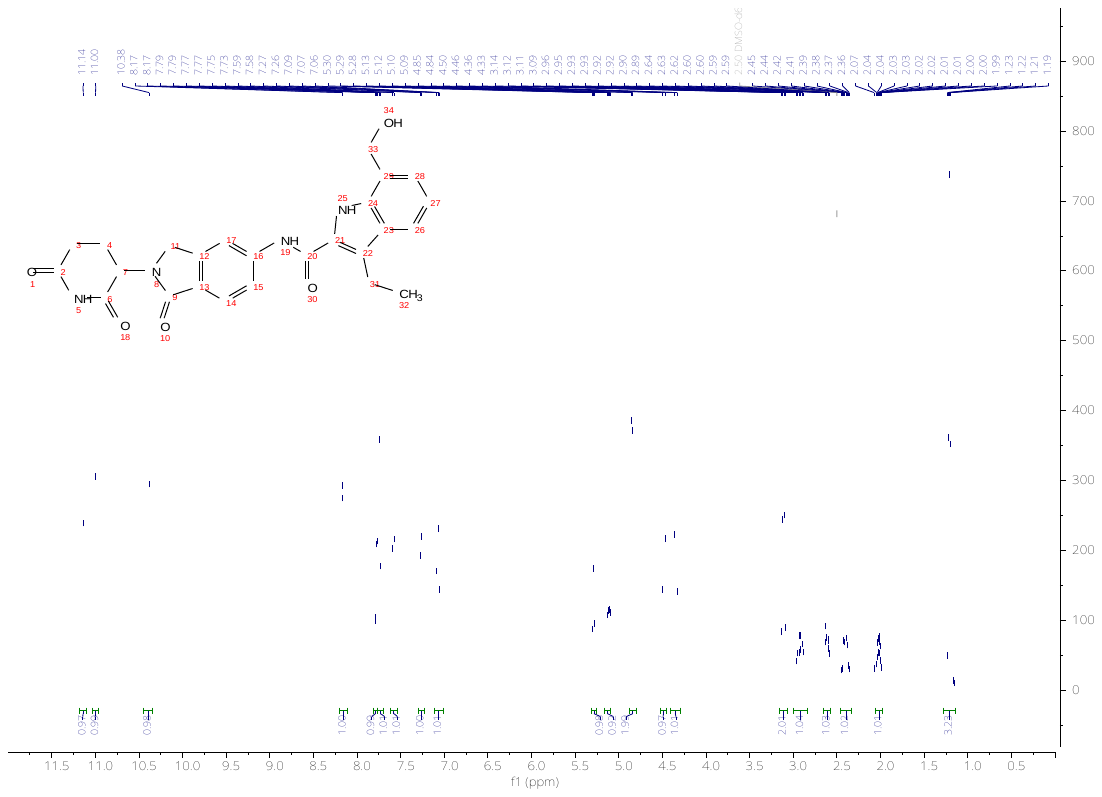

**Figure S40.** ^1^H NMR spectrum of **19** in DMSO-*d_6_*.

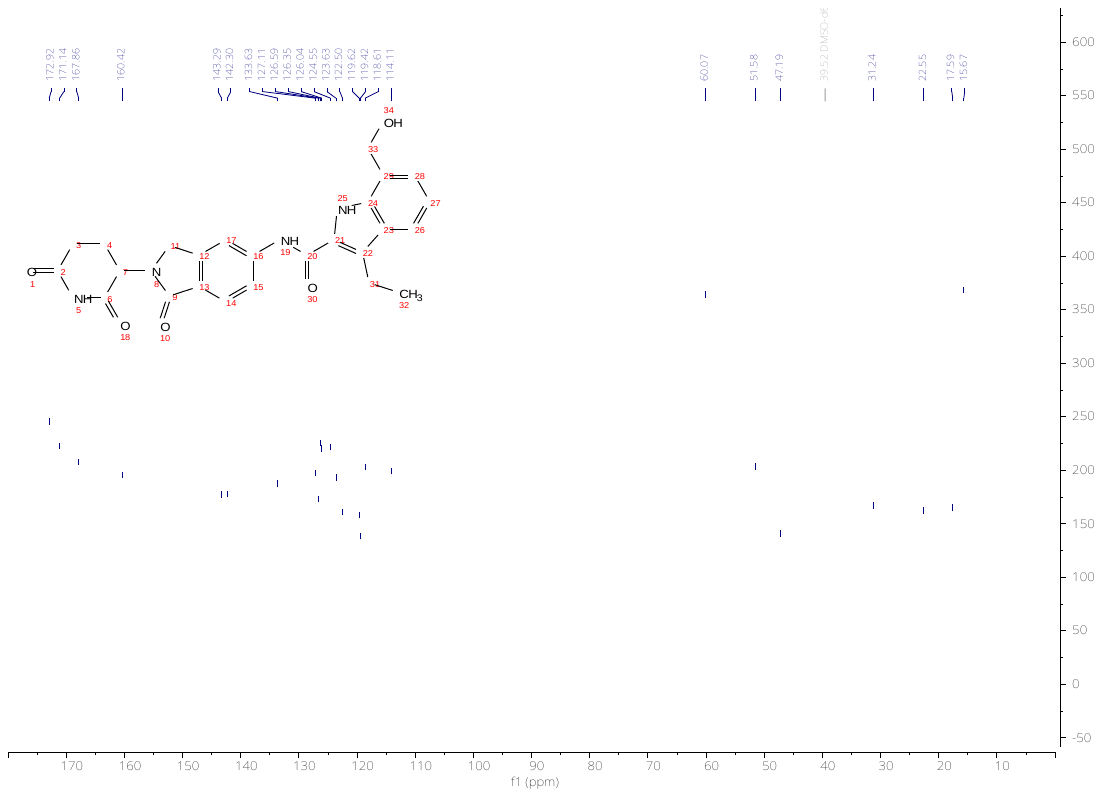

**Figure S41.** ^13^C NMR spectrum of **19** in DMSO-*d_6_*.

**LC Chromatograms of Compounds 1–19.**

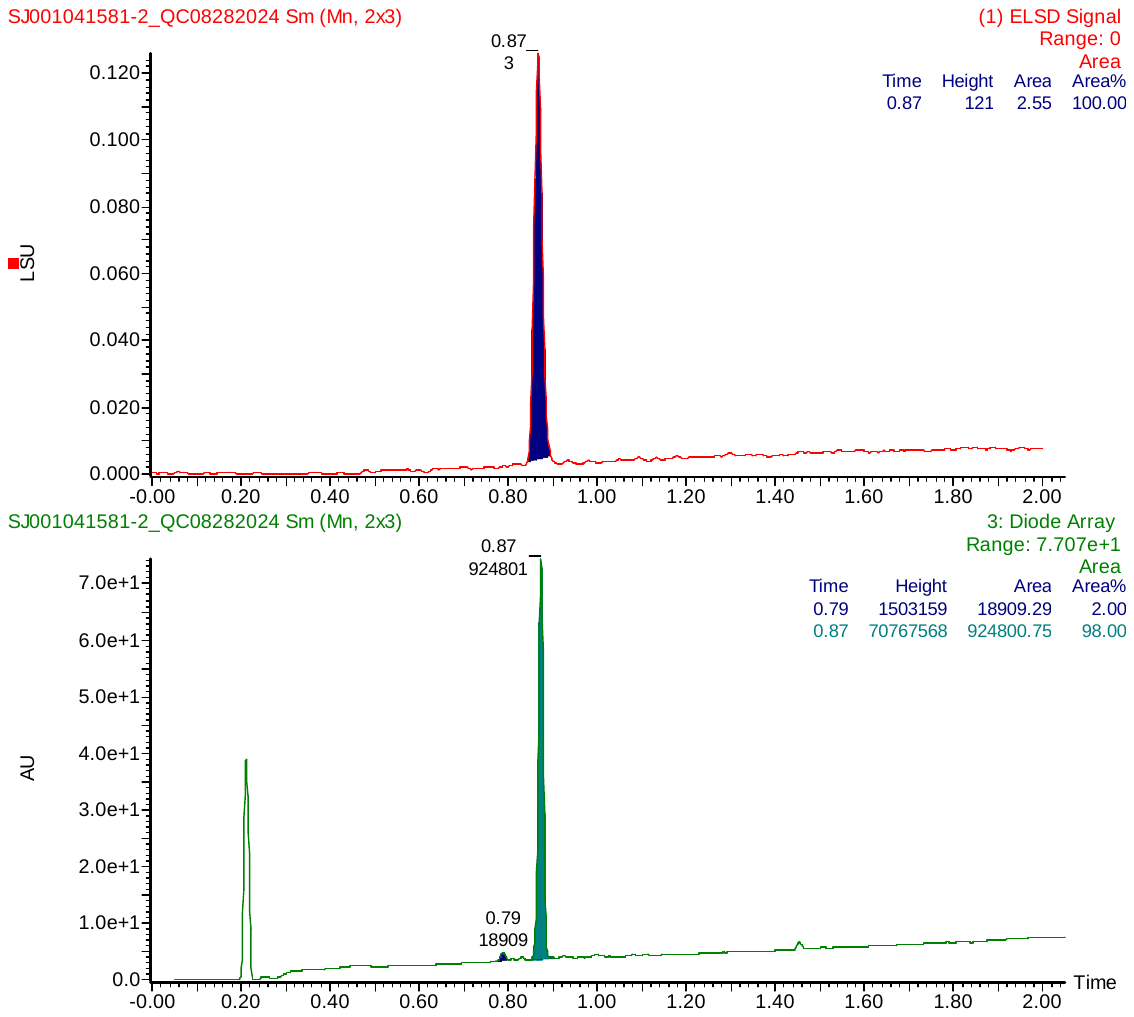

**Figure S42.** LC Chromatogram of **1** detected by ELSD and UV diode array.

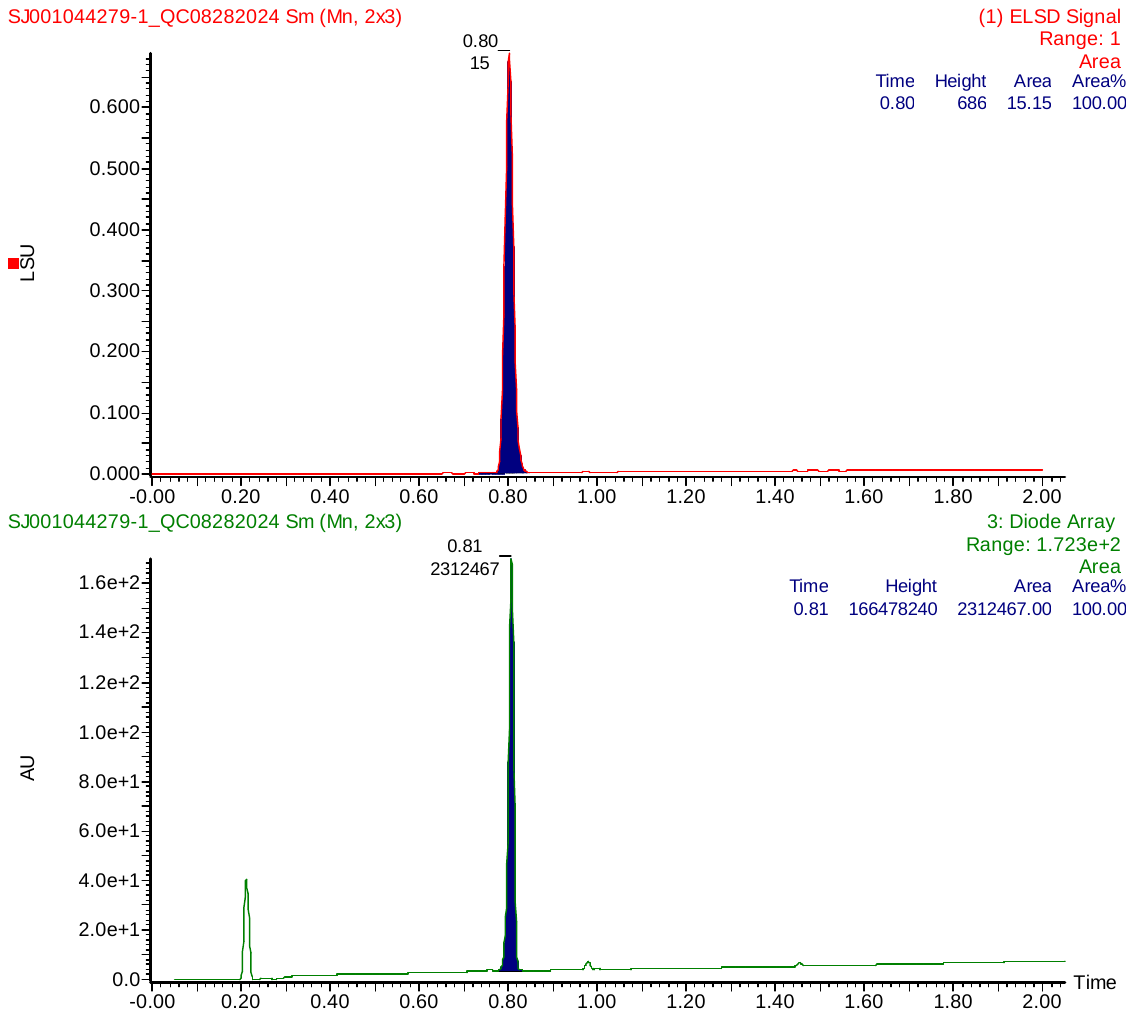

**Figure S43.** LC Chromatogram of **2** detected by ELSD and UV diode array.

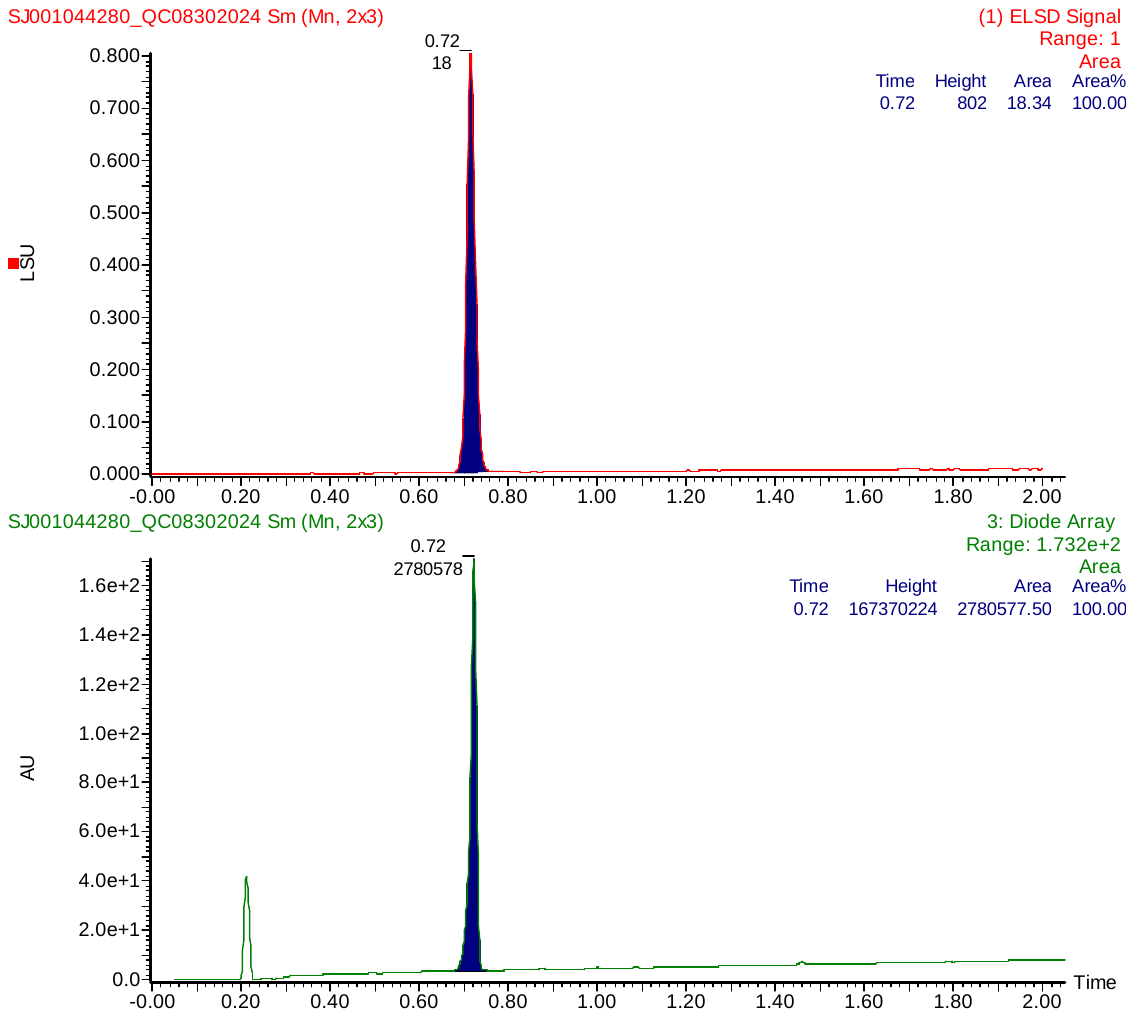

**Figure S44.** LC Chromatogram of **3** detected by ELSD and UV diode array.

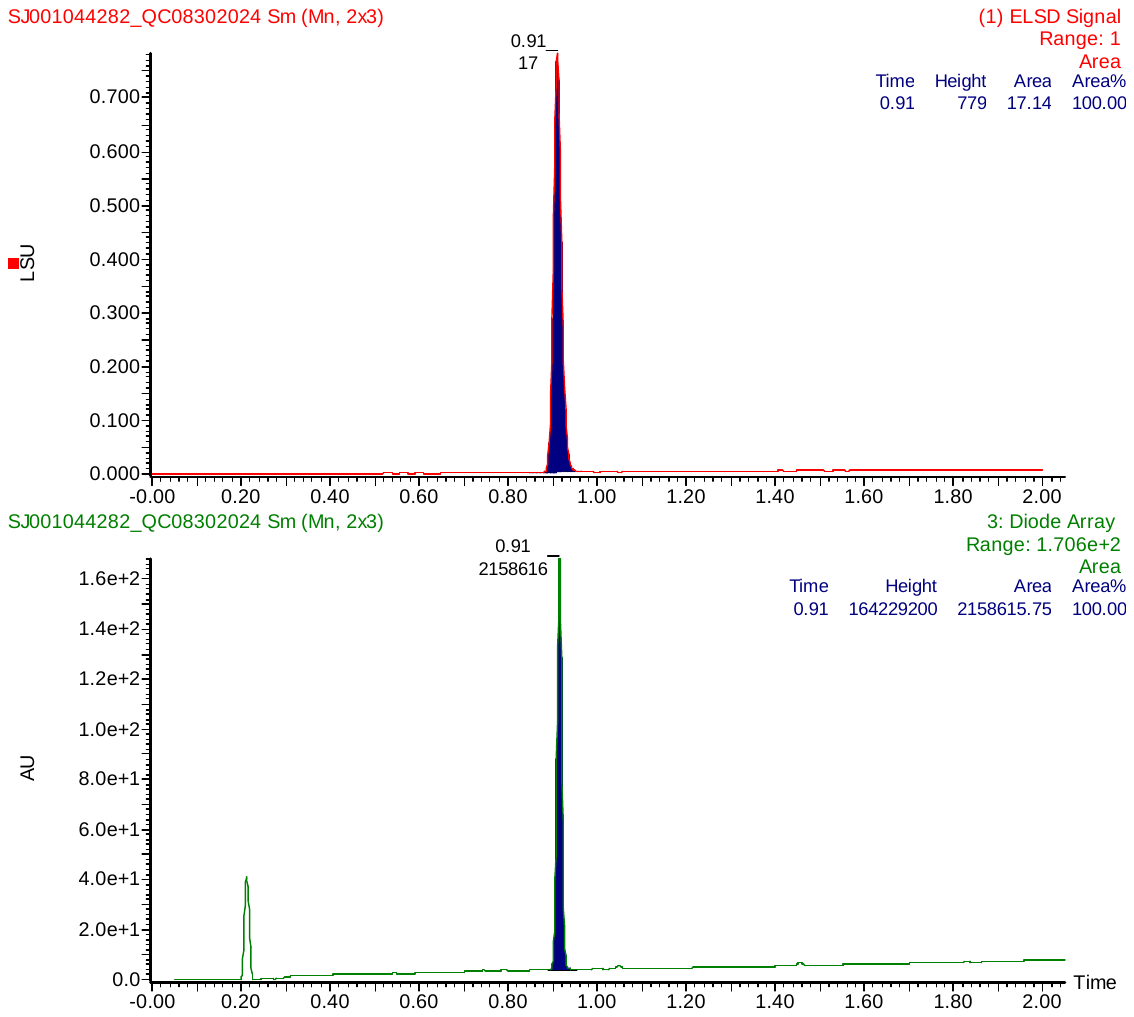

**Figure S45.** LC Chromatogram of **4** detected by ELSD and UV diode array.

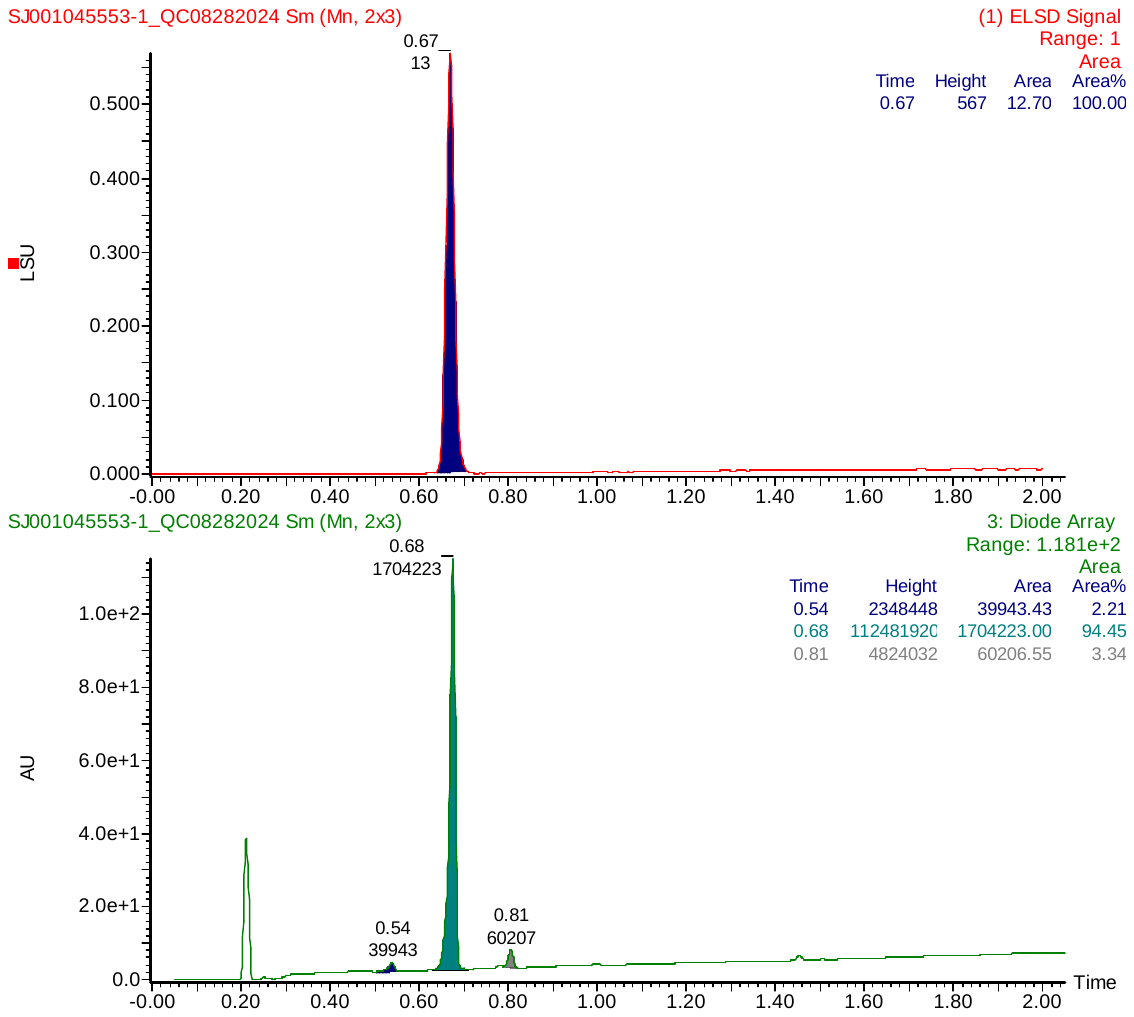

**Figure S46.** LC Chromatogram of **5** detected by ELSD and UV diode array.

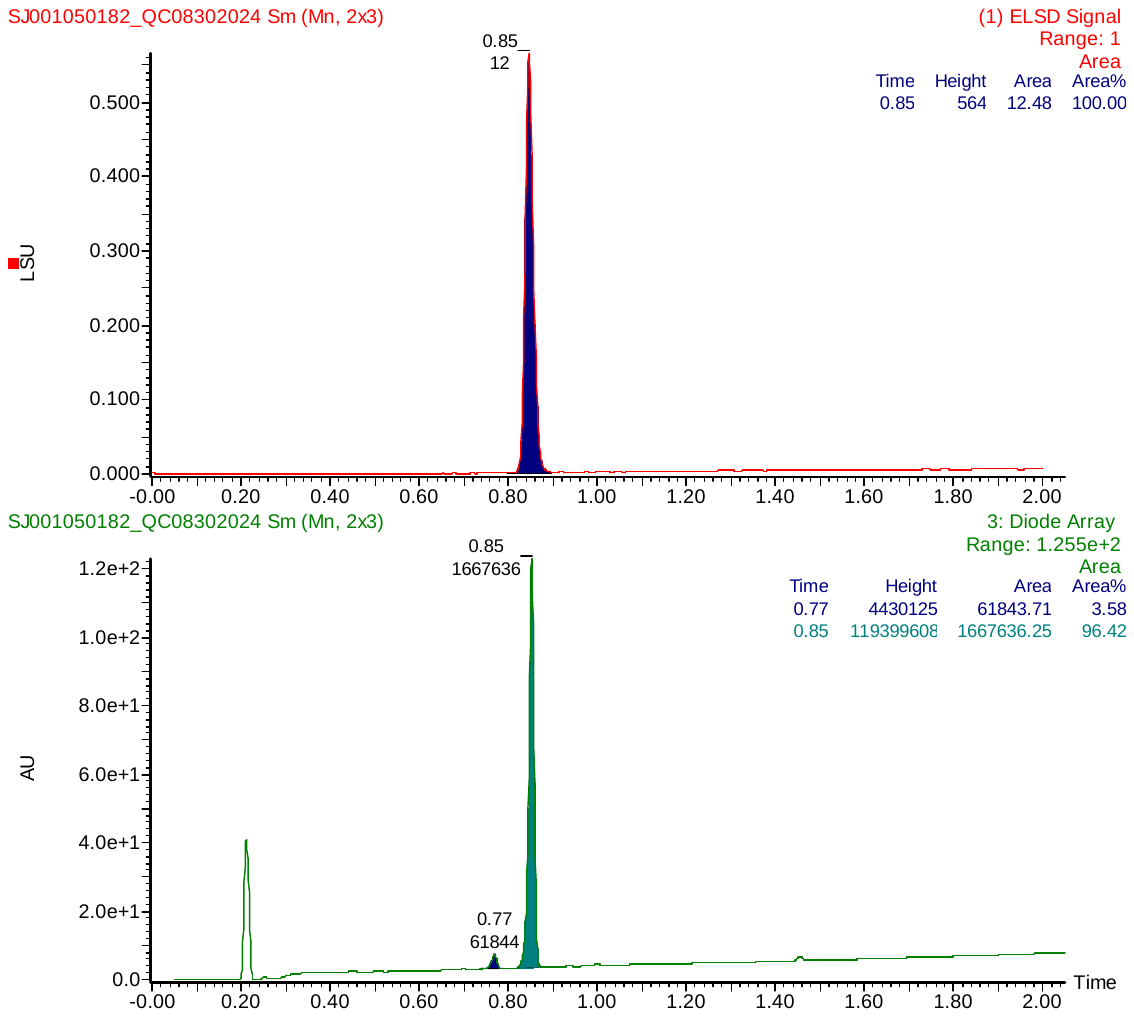

**Figure S47.** LC Chromatogram of **6** detected by ELSD and UV diode array.

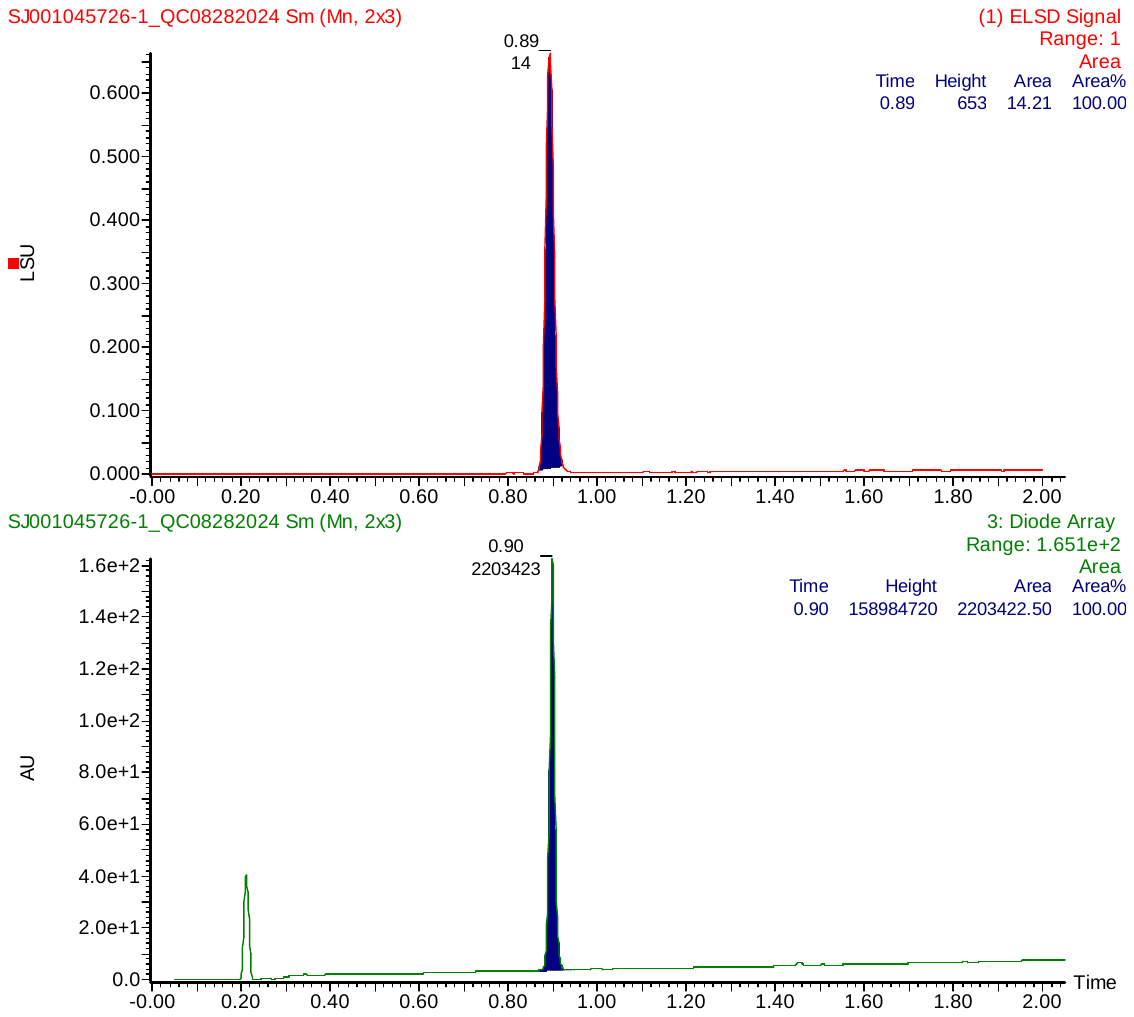

**Figure S48.** LC Chromatogram of **7** detected by ELSD and UV diode array.

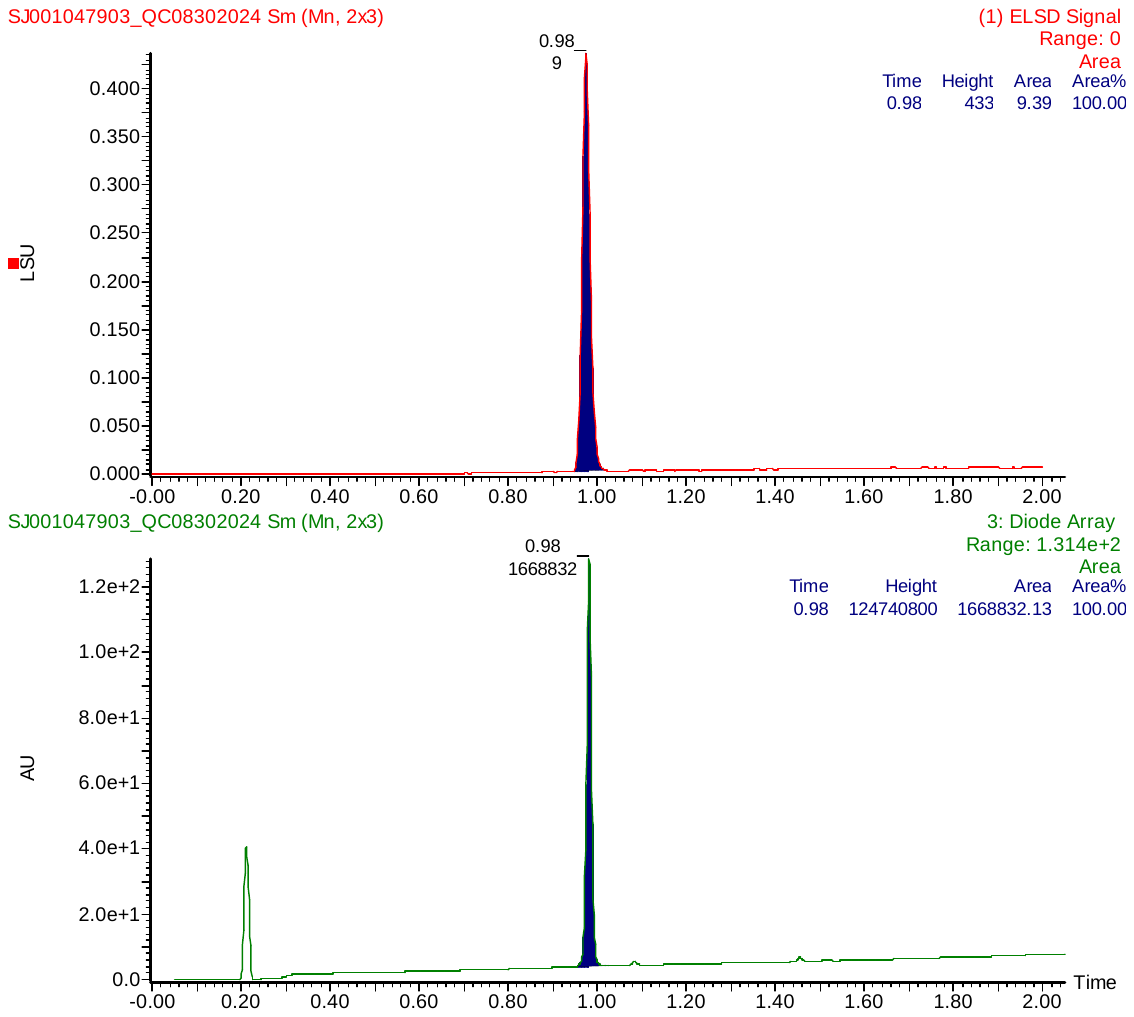

**Figure S49.** LC Chromatogram of **8** detected by ELSD and UV diode array.

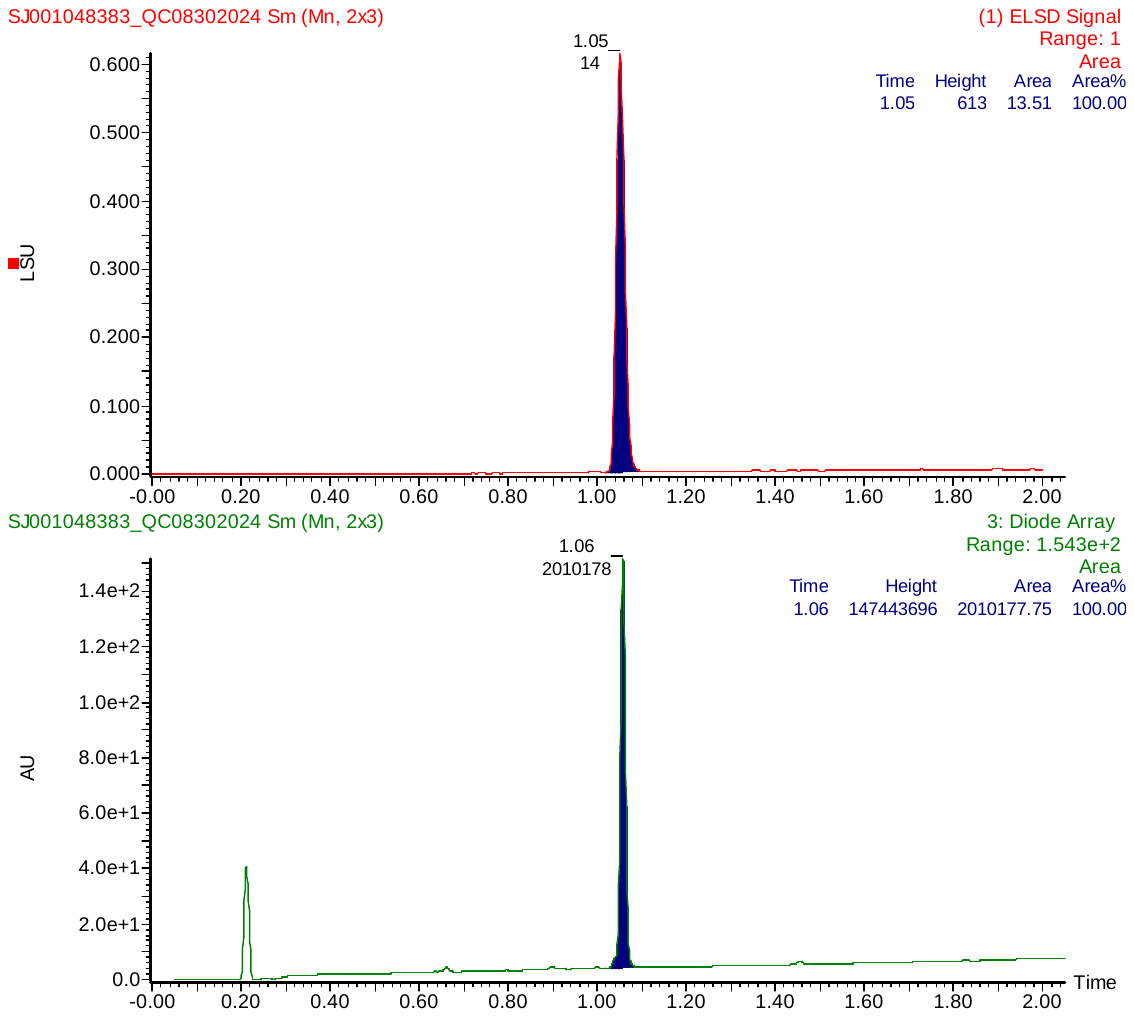

**Figure S50.** LC Chromatogram of **9** detected by ELSD and UV diode array.

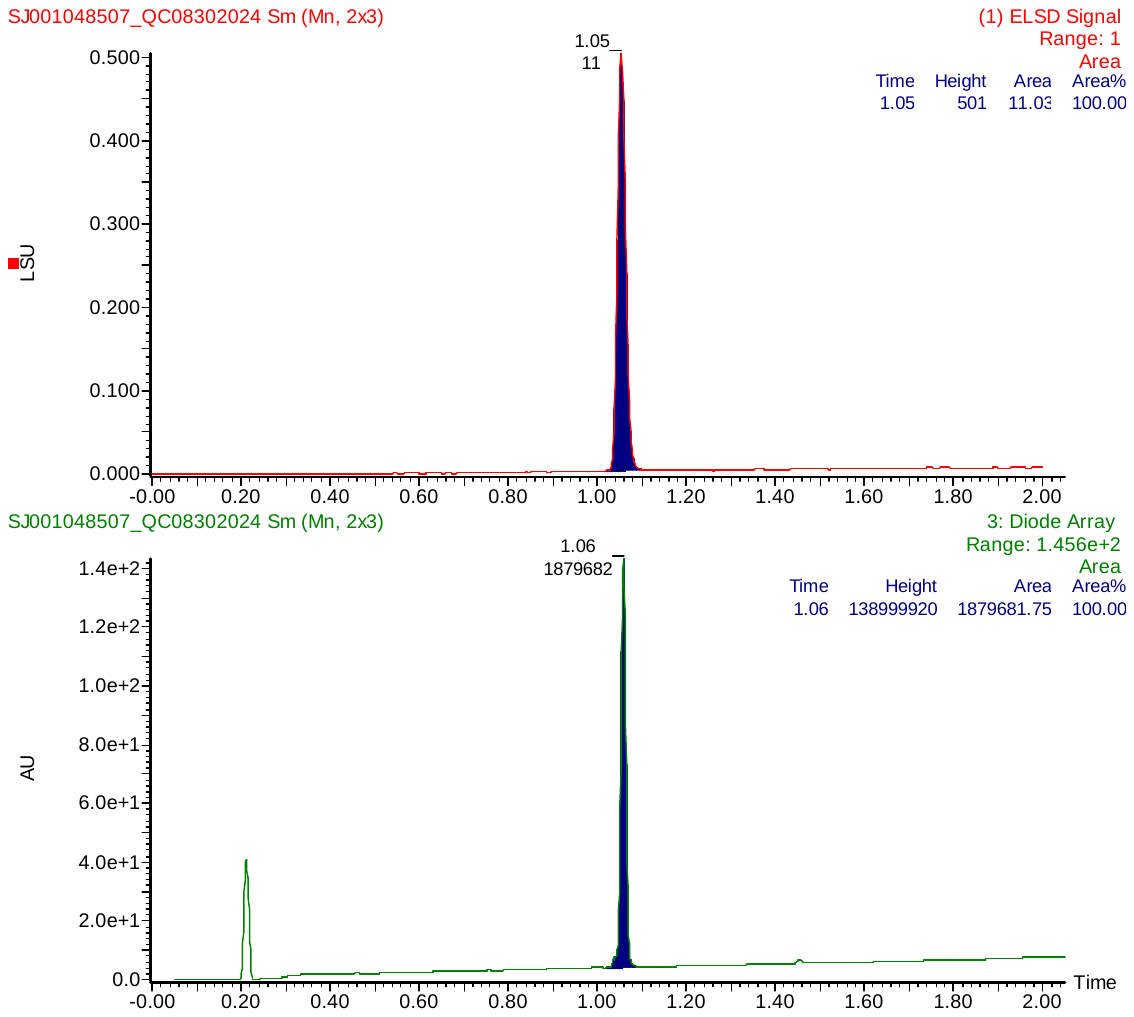

**Figure S51.** LC Chromatogram of **10** detected by ELSD and UV diode array.

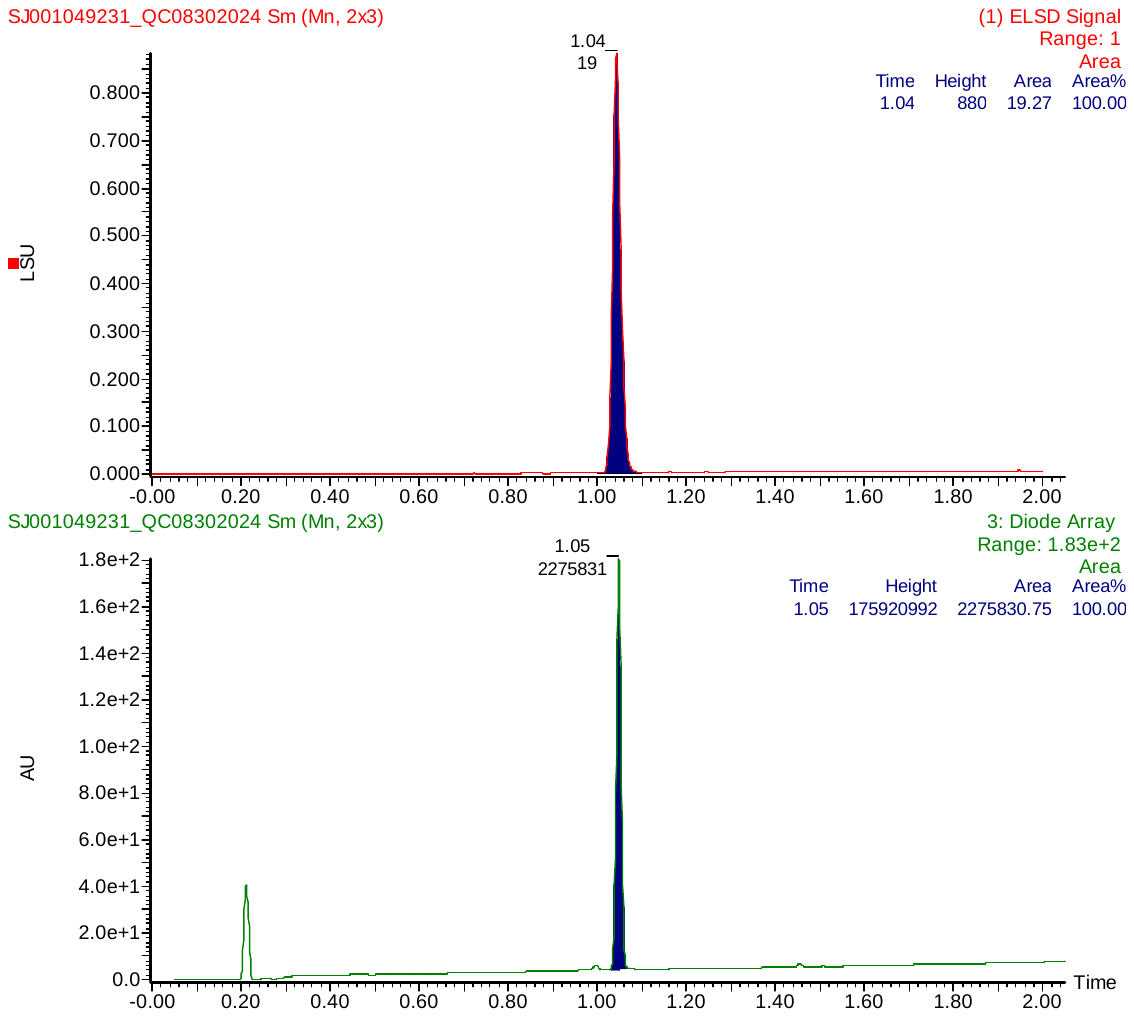

**Figure S52.** LC Chromatogram of **11** detected by ELSD and UV diode array.

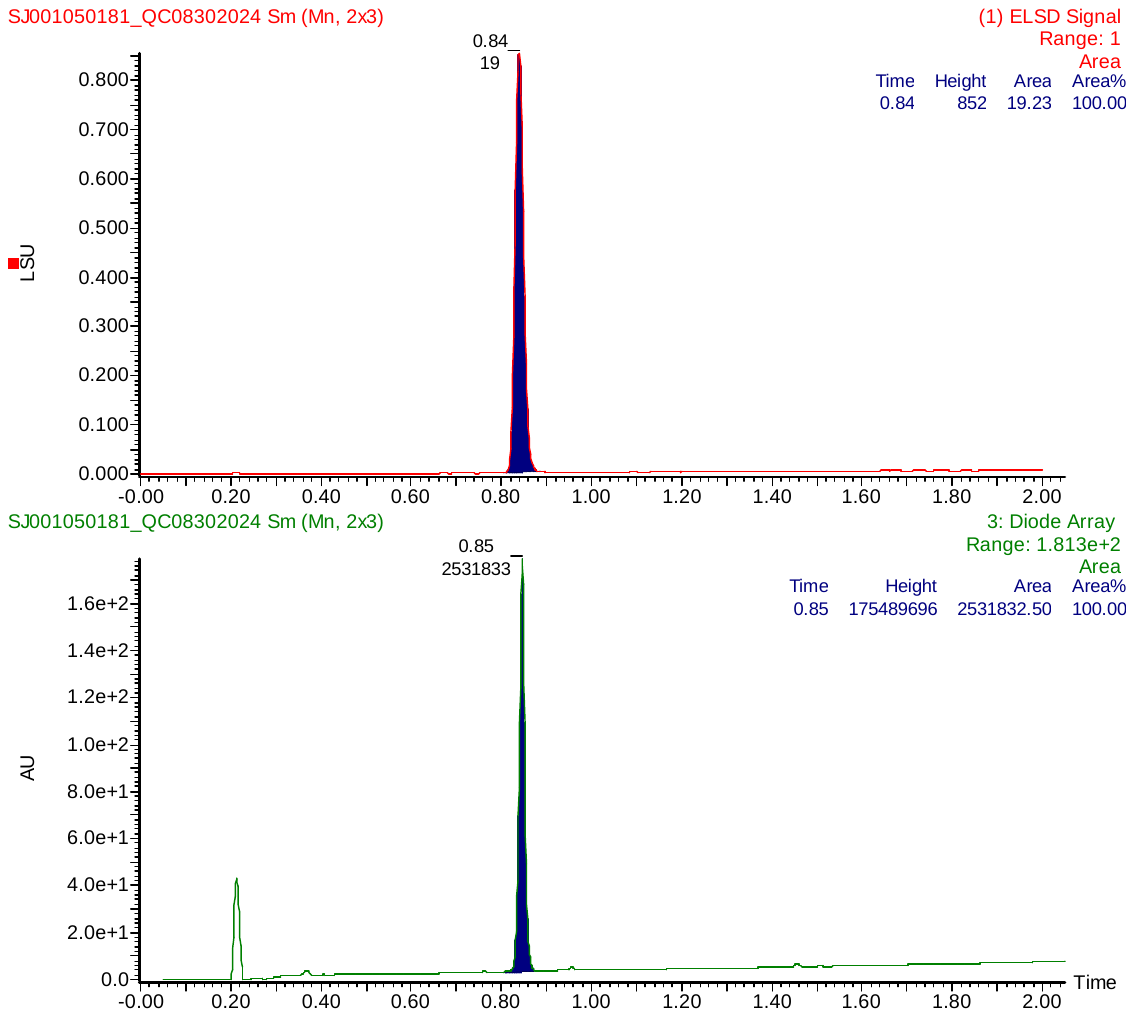

**Figure S53.** LC Chromatogram of **12** detected by ELSD and UV diode array.

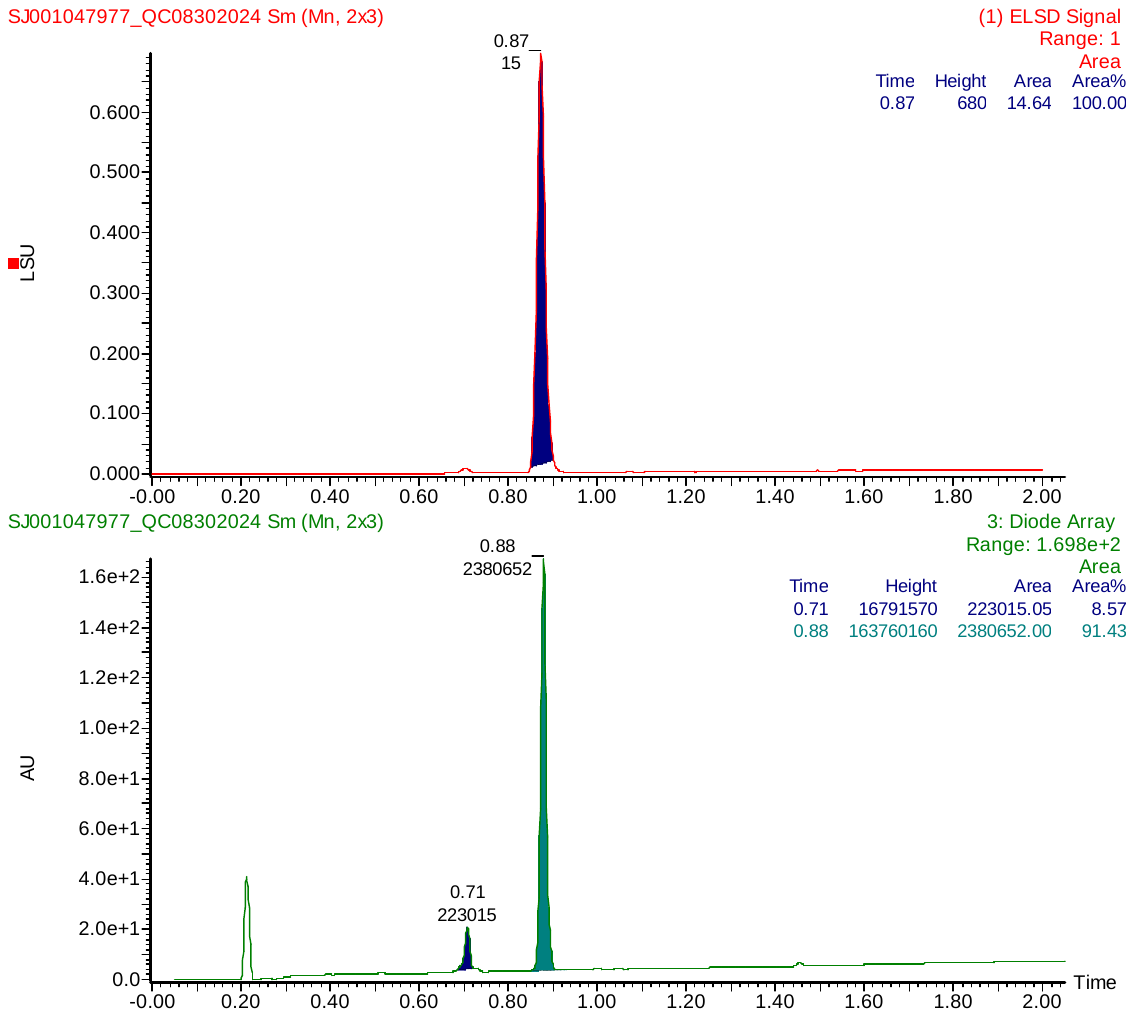

**Figure S54.** LC Chromatogram of **13** detected by ELSD and UV diode array.

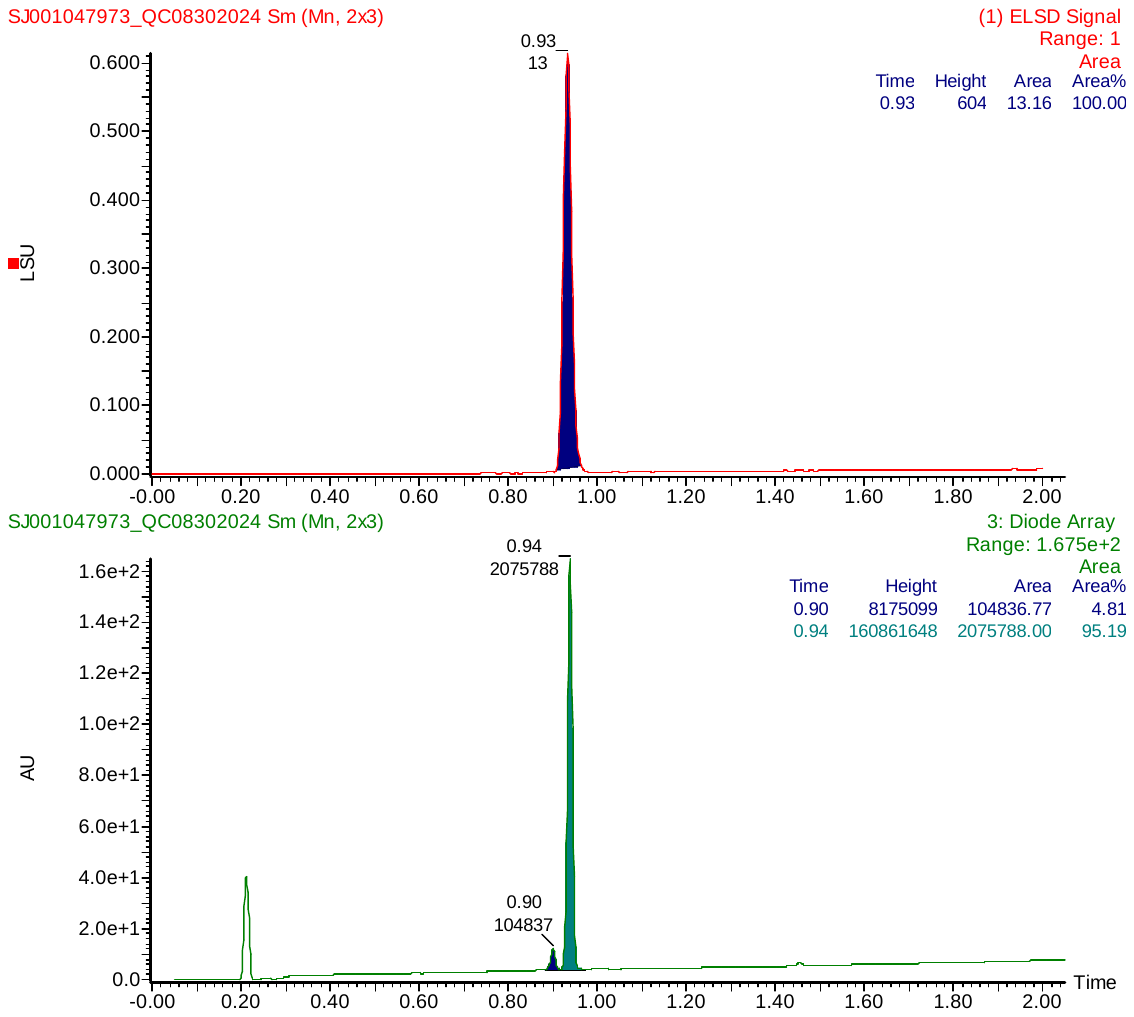

**Figure S55.** LC Chromatogram of **14** detected by ELSD and UV diode array.

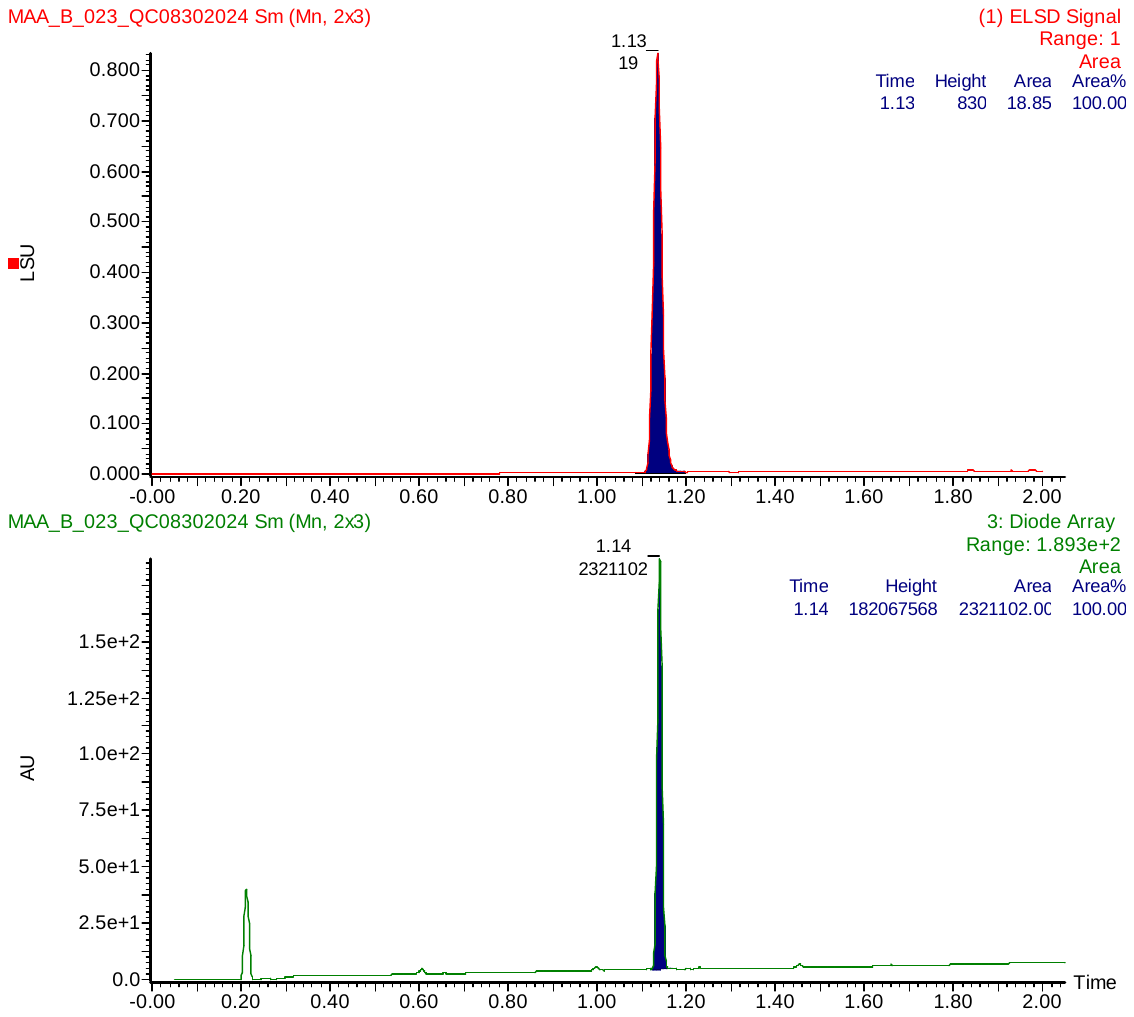

**Figure S56.** LC Chromatogram of **15** detected by ELSD and UV diode array.

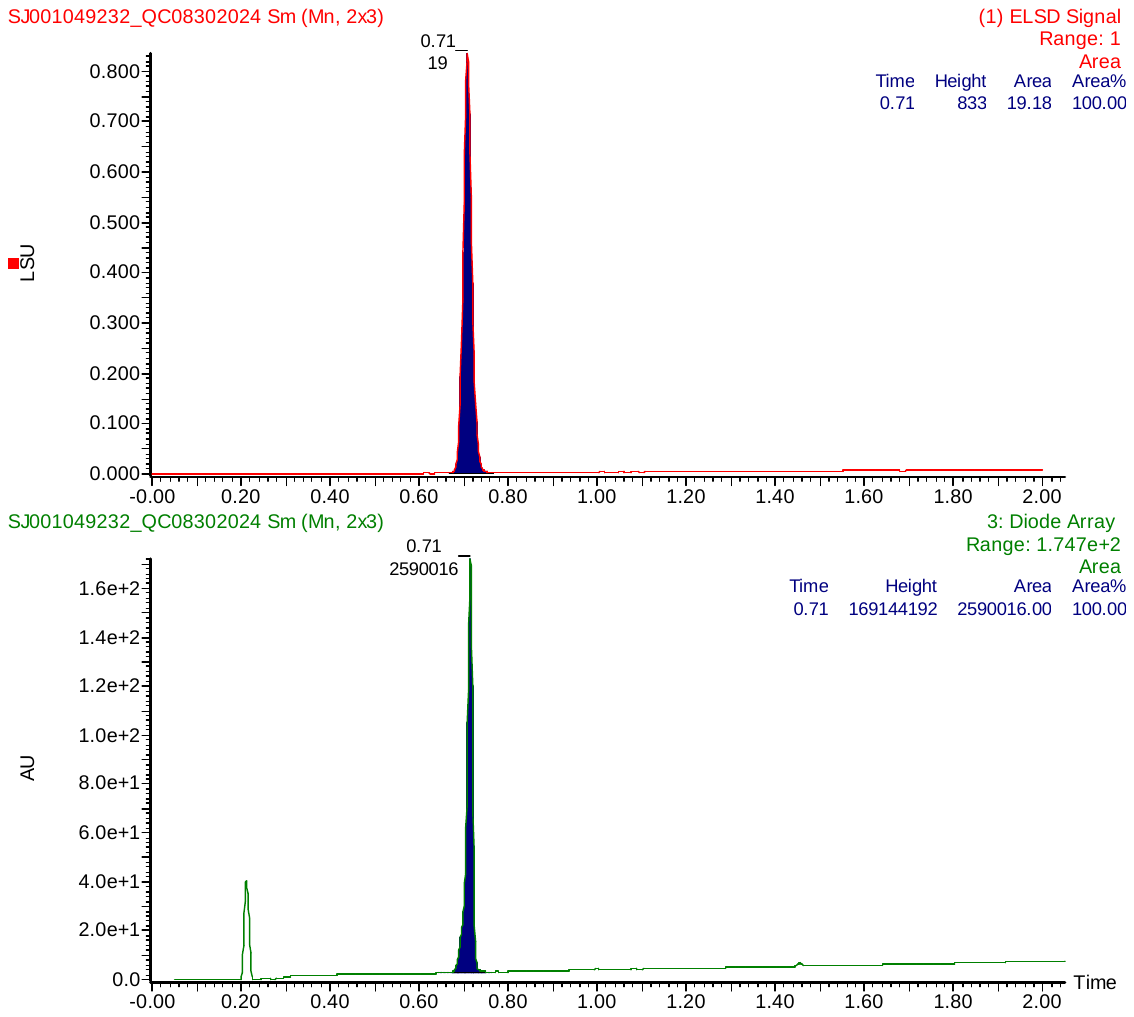

**Figure S57.** LC Chromatogram of **16** detected by ELSD and UV diode array.

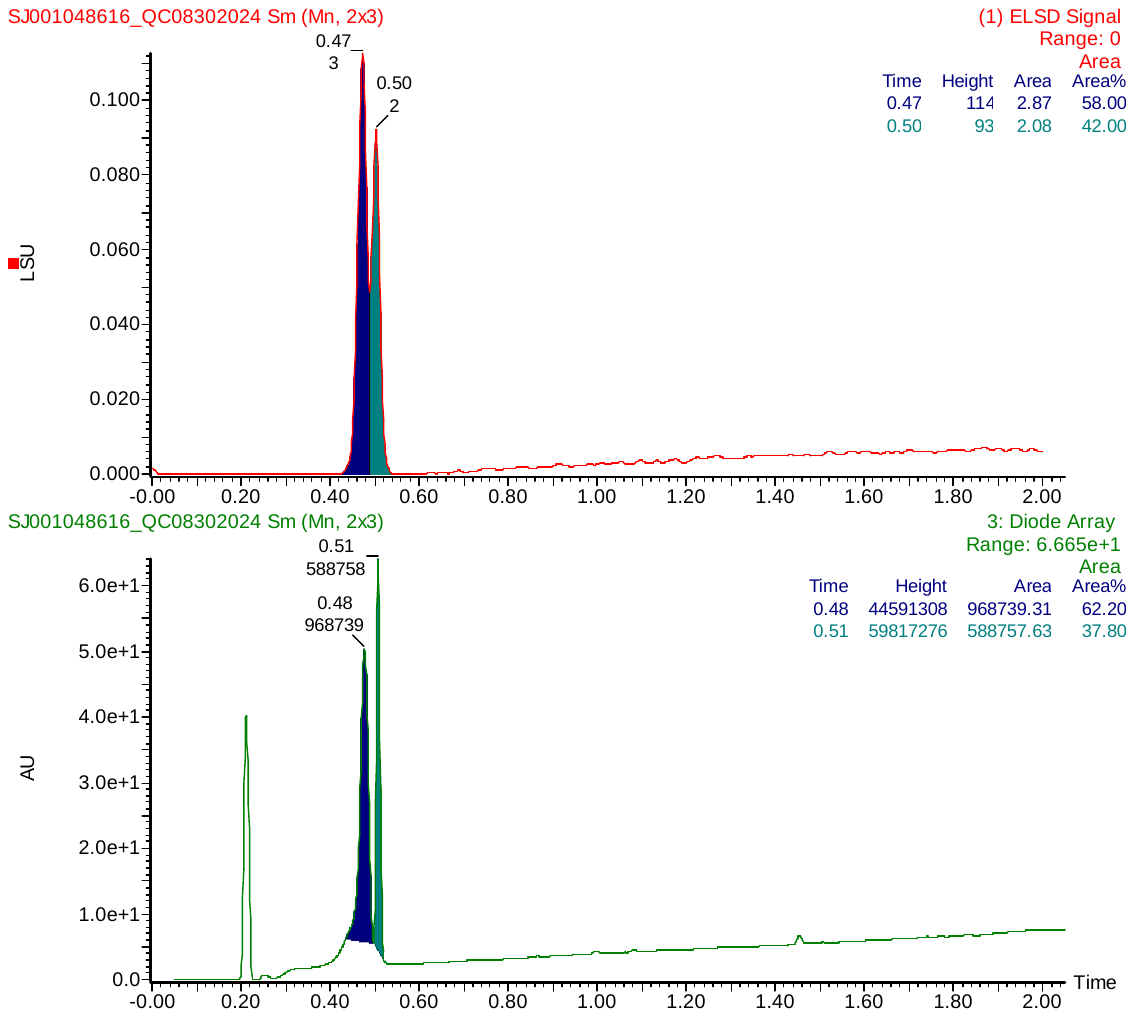

**Figure S58.** LC Chromatogram of **17** detected by ELSD and UV diode array.

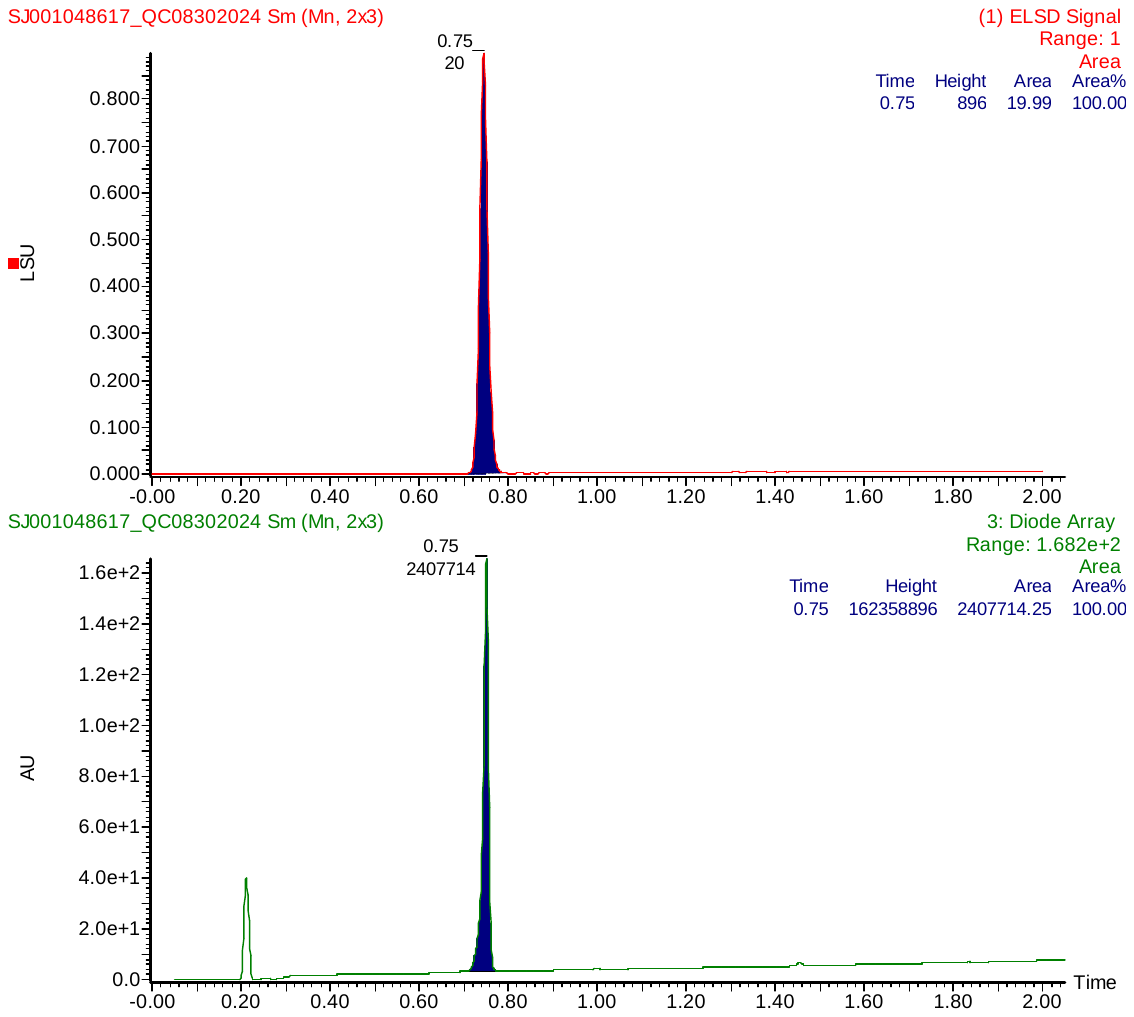

**Figure S59.** LC Chromatogram of **18** detected by ELSD and UV diode array.

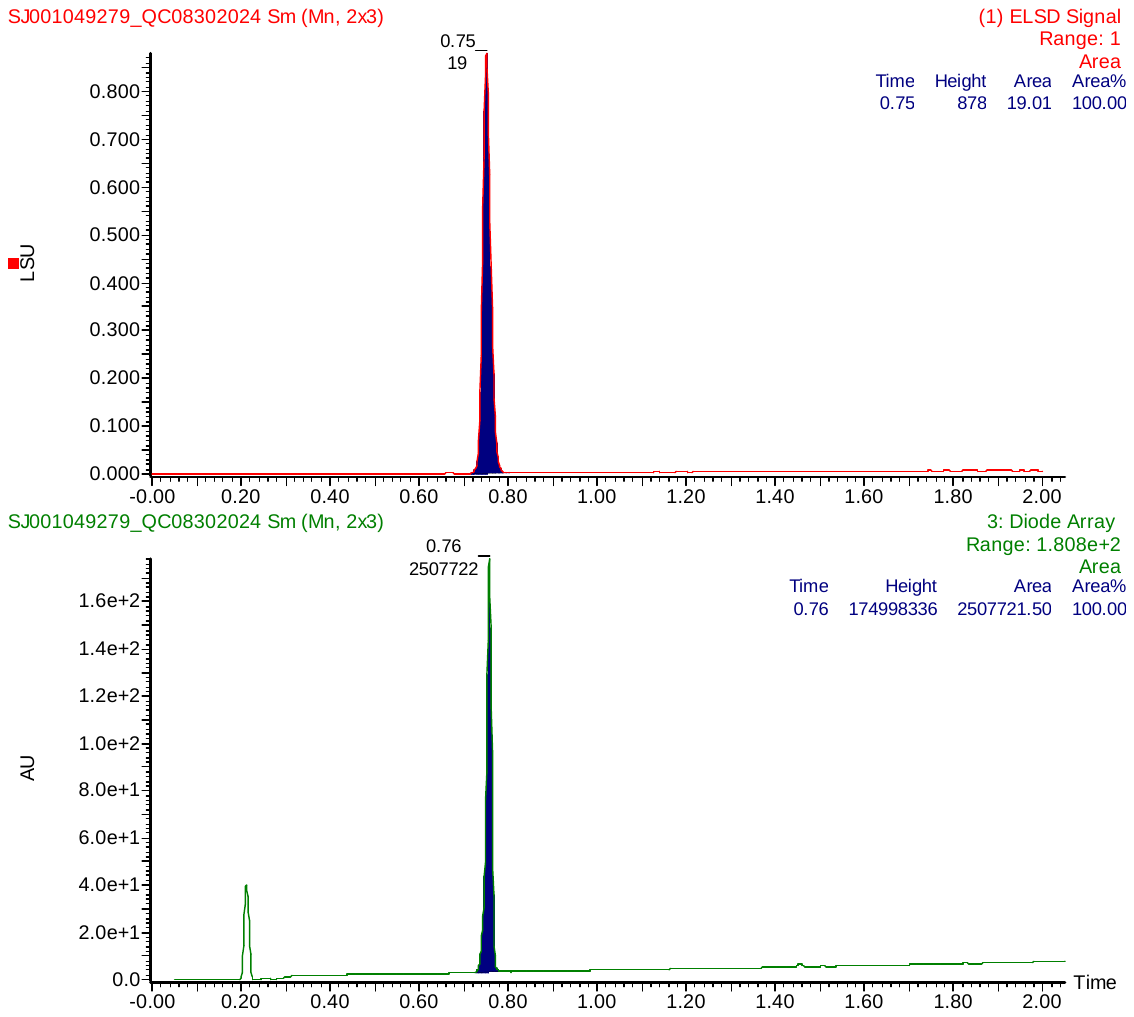

**Figure S60.** LC Chromatogram of **19** detected by ELSD and UV diode array.
